# A guide to the *in vitro* reconstitution of Cdc42 GTPase activity and its regulation

**DOI:** 10.1101/2023.04.24.538075

**Authors:** Sophie Tschirpke, Frank van Opstal, Ramon van der Valk, Werner K-G. Daalman, Liedewij Laan

**Affiliations:** Bionanoscience Department, Delft University of Technology, Delft, the Netherlands

## Abstract

Cdc42 is a small Rho-type GTPase and the main regulator of cell division in eukaryotes. It is surrounded by a large network of regulatory proteins. To understand the processes around cell division, in-depth understanding of Cdc42 and its regulation is required. *In vitro* reconstitutions are a suitable tool for such detailed mechanistic studies, as they allow a high level of control over the conditions and components used and. For these Cdc42 and its regulators need to be expressed, purified, and tested for their activity. There are many methods described for this, but their details, possible difficulties, and points of failure are rarely discussed. This makes *in vitro* studies on Cdc42 less accessible to scientists that have a background different from biochemistry. We here present our experience with working with Cdc42 *in vitro*. We describe the recombinant expression and purification behaviour of 12 Cdc42, six Cdc42-mNeonGreen^*SW*^ and four Cdc42-sfGFP^*SW*^ constructs in *E. coli*. We explore Cdc42 dimerisation *in vitro* and assess its activity using GTPase Glo assays and Flag-pulldown assays. GTPase Glo assays turn out to be a reliable tool to quantitatively asses GTPase activities, wheareas pulldown experiments are more error prone. We find that most Cdc42 constructs, with the exception of those with an N-terminal Twin-Step-tag, show a similar GTPase activity and interaction with the GDP/GTP exchange factor Cdc24. We close with using enterokinase and TEV protease to generate untagged Cdc42. Enterokinase also cuts Cdc42 in an undesired position. TEV protease leads to the desired product, which retains its GTPase activity but shows a reduced Cdc24 interaction. The work presented here acts as a guide for scientists desiring to work with Cdc42 *in vitro* through describing Cdc42’s properties in detail and examining assays that can be used to study its behaviour or act as activity checks.

**Graphical abstract:** 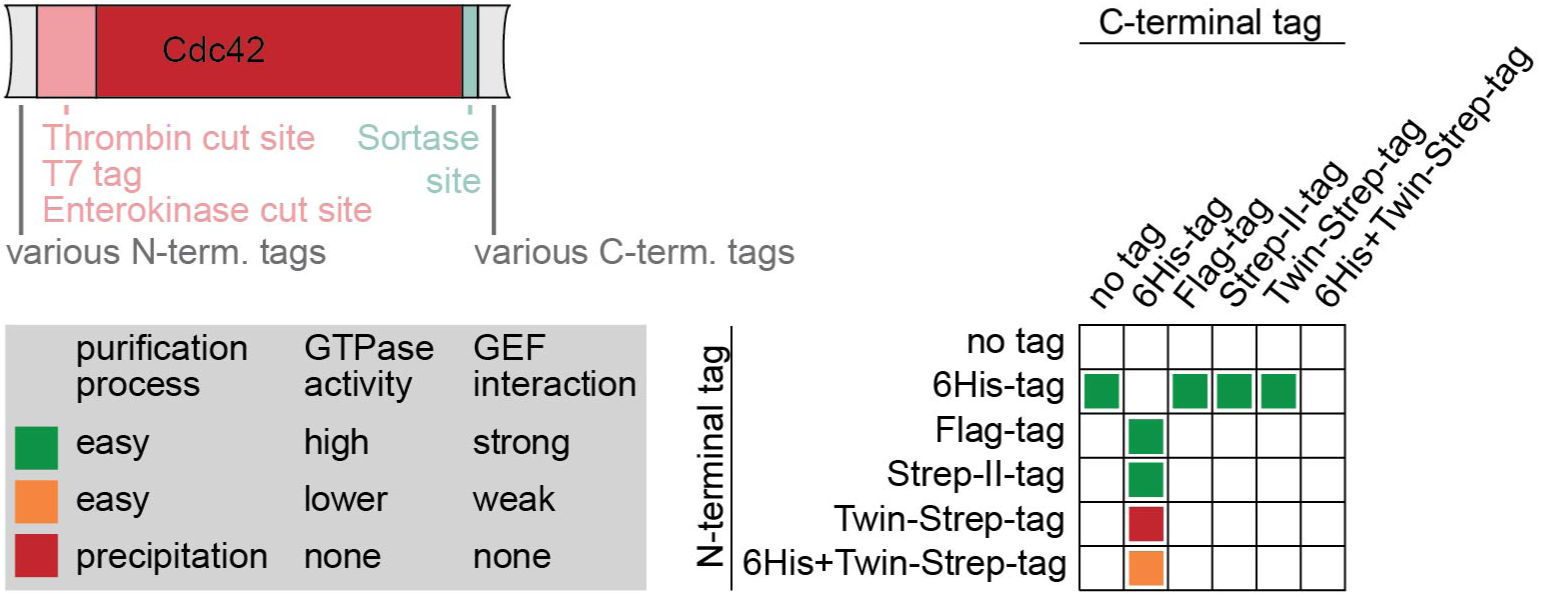

## Introduction

Cellular processes arise through complex interactions of many components. *In vivo* investigations give insights into the overall cellular function of a component, and *in vitro* assays elucidate the specifics of single interactions and component functions. Bridging the gap between highly controlled single to a few component *in vitro* assays and highly complex *in vivo* processes are *in vitro* reconstitutions (for example (Bieling et al., 2016; Roth, Gârlea, Vleugel, Mulder, & Dogterom, 2019; Bezeljak, Loya, Kaczmarek, Saunders, & Loose, 2020; Kohyama, Merino-Salomón, & Schwille, 2022)). Here the interplay between several components, synergies, and emergent behaviours can be characterised. Factors such as concentrations, component ratios, and temperature can be manipulated and their effect on the overall system can be observed - giving rise to an an in-depth understanding of the system and a base towards building a synthetic cell. With increasing amount of added components the complexity of the system increases - it moves towards a cell-like state - but it also becomes experimentally more challenging (Vendel, Tschirpke, Shamsi, Dogterom, & Laan, 2019); for example because an altered property of a single component can change the overall behaviour of the entire system. Thus, a high level of control and knowledge over the components is needed. The system surrounding the Rho-type GTPase Cdc42 is an interesting case for *in vitro* reconstitutions: Cdc42 is the main regulator of polarity establishment and cell division in eukaryotes (Diepeveen, Gehrmann, Pourquié, Abeel, & Laan, 2018). It is part of a complex network of polarity proteins, including GDP/GTP exchange factors (GEFs), GTPase activating proteins (GAPs), a guanine nucleotide dissociation inhibitor (GDI), scaffold proteins, and other regulatory proteins, that all interact with each other and with Cdc42 in particular (Gao et al., 2011; Costanzo et al., 2016; Daalman, Sweep, & Laan, 2020). Cdc42’s properties and the protein-protein interactions of the polarity network have been studied *in vivo* and in isolation *in vitro* (Park & Bi, 2007), elucidating aspects of the network’s properties. For example, several of Cdc42’s binding partners, properties of its GTPase activity, and kinetics of its membrane interaction were be examined through *in vitro* studies using one to three proteins in isolation (Supplement S 1). However, despite these detailed information accumulated over decades of study the molecular mechanism driving Cdc42 accumulation is still heavily debated (Vendel et al., 2019; Goryachev & Leda, 2017). One reason for the ongoing debate is the high level of interconnection and redundancy present in the yeast polarity system, which makes this system difficult to model and work with. *In vitro* reconstitutions, allowing for a certain, yet controllable, level of complexity, could show the molecular mechanism of Cdc42 accumulation (Vendel et al., 2019).

For *in vitro* reconstitutions Cdc42 constructs (with purification tags) need to be designed, expressed, purified, stored until use, and tested for their activity. One of the first decisions that thus needs to be made is that of native or recombinant protein expression/purification. In native purification the protein of interest (POI) is expressed in its native organism, in the recombinant approach a different organism is used. Native purification gives the most biologically relevant protein, but expression and purification protocols are usually less established and more difficult to follow, and the obtained POI batch can contain small amounts of co-purified POI binding partners and consists of a mixture of POI with various forms of post-translational modifications (for example, for *Saccharomyces cerevisiae* Cdc42: ubiquitinylation (Swaney et al., 2013; Back, Gorman, Vogel, & Silva, 2019), phosphorylation (Lanz et al., 2021), farnesylation and geranyl-geranylation (Caplin, Hettich, & Marshall, 1994). The presence of post-translationally modified protein is also an advantage of a native purification: if the post-translational modification is an essential feature of the POI, its presence is required. Previous *in vitro* studies on Cdc42 obtained the protein using (1) *Escherichia coli* based expression systems (Golding, Visco, Bieling, & Bement, 2019), (2) insect cell expression systems (Zheng, Cerione, & Bender, 1994; Zheng, Bender, & Cerione, 1995; Zhang & Zheng, 1998; Zhang, Zhang, Collins, Johnson, & Zheng, 1999; Kozminski et al., 2003; Johnson, Erickson, & Cerione, 2009, 2012), or (3) through purification of membrane-bound Cdc42 from yeast (Das et al., 2012; Rapali et al., 2017). We chose to purify *S. cerevisiae* Cdc42 recombinantly from *Escherichia coli* due to the method’s advantages: (1) Recombinant purification from *E. coli* is widely used and protocols are established and easy to follow. *E. coli* grows fast and reliably and protein can be over-expressed in a high yield. (2) As Cdc42 is not native in *E. coli*, it will have little to no binding-partners that it can co-purify. One can thus be more certain that the obtained Cdc42 batch is only Cdc42. This is especially important if Cdc42 ought to be used to quantitatively study its interaction with its binding partners. If the Cdc42 batch contains already small amounts of binding partners, claims about the effect of a binding partner on Cdc42 and the concentration-dependence of this effect are less accurate. We like to point out that Cdc42 purified in this way does not possess its C-terminal prenyl group, which is responsible for anchoring Cdc42 to the membrane and binding to the GDI Rdi1 (Cox & Der, 1992; Koch et al., 1997). This group would need to be added later in an *in vitro* reaction (Golding et al., 2019; Tschirpke, Spitzbarth, Hettema, Eelkema, & Laan, 2024; Popp & Ploegh, 2011).

There are many frequently used methods for protein expression, purification, and assays that assess protein activity and their interactions, but their details, possible difficulties, and points of failure are rarely discussed. This makes *in vitro* studies, and reconstitutions in particular, with Cdc42 less accessible to scientist that have a background different from biochemistry. Here we set out to make the *in vitro* use of (*Saccharomyces cerevisiae*) Cdc42 more accessible to a broad audience, through describing our work and experience with various Cdc42 constructs. The goal of this paper is not to map out all properties of all constructs used, but to showcase what steps and difficulties working with Cdc42 *in vitro* entails. We include positive as well as negative results and point out heterogeneous behaviours of Cdc42. We start by describing our Cdc42 construct design in detail. We then show the recombinant expression and purification behaviour of Cdc42 and Cdc42-sfGFP and -mNeongreen sandwich fusions in *E. coli* and discuss Cdc42 dimerisation *in vitro*. We then used GTPase and pulldown assays to (1) examine our Cdc42 constructs on their functionality - their GTPase activity and interaction with two binding partners, and (2) to assess how reliable and straightforward these assays are to test protein properties. We close with exploring how suitable two cleavage enzymes, enterokinase and TEV protease, are for cleaving tags off Cdc42.

## Results

### Cdc42 constructs

Purification of proteins for *in vitro* studies is generally done by attaching an N- or C-terminal purification tag to the POI. The rational behind the placement of purification tags and their effect on the protein’s expression/purification behaviour and properties are rarely discussed. This contributes to making *in vitro* studies less accessible for non-biochemists. Further, tag cleavage is not always easy or possible. To cleave off a tag a recognition site for a cleavage enzyme needs to be placed in-between the POI and purification tag. Cleavage enzymes have a relatively high sequence specificity, but can also cleave within the POI causing degradation products. The enzyme’s cleavage behaviour and efficiency need to be tested for every POI to ensure proper matching. After the cleavage reaction the tagged, untagged, degraded POI species and cleavage enzyme need to be separated, adding another undesirable and yield-reducing purification step. In other cases the cleavage of the purification tag is not possible. For example when the tag is used to bind the POI to beads or a modified surface to study protein-protein interactions.

Cdc42 is a small Rho-type GTPase of 191 amino acids (AAs)/21 kDa, consisting mainly of a Rho-domain, that is responsible for its catalytic activity, and a short flexible N-terminal region made of the polybasic region (PBR) directly followed by the CAAX box. The PBR is a five AA long unstructured region of mostly positively charged amino acids that has been linked to dimerisation and supports binding to negatively charged membranes (Williams, 2003). The CAAX-box is a four AA sequence at the C-terminus, at which the protein gets post-translationally modified: a prenyl group (geranylgeranyl or farnesyl) gets appended, allowing the protein to bind to membranes (Cox & Der, 1992).

So-far, *in vitro* studies with 6His-tagged (Zhang & Zheng, 1998; Zhang, Zhang, Wang, & Zheng, 2000; Kozminski et al., 2003; Johnson et al., 2012; Golding et al., 2019; Sonal, Heermann, & Schwille, 2022), GST-tagged (Zheng et al., 1995; Bose et al., 2001; Kozminski et al., 2003; Das et al., 2012), and 6His-Strep-II-tagged (Rapali et al., 2017) Cdc42 were conducted. Of these references, only one reported to have cleaved off the tag (Zhang & Zheng, 1998). All of these constructs have the purification tag placed on the N-terminus. It is likely the case because *in vivo* Cdc42 is prenylated at its C-terminus. Even though not all mentioned studies used prenylated Cdc42, all left the C-terminus unmodified. We set out to explore the effect of N- and C-terminal additions, including various purification tags, on Cdc42. We constructed three types of constructs:

- Type H: Here only a single 6His-tag is directly appended to the N-terminus of Cdc42 (Fig. 1). This construct has the least amount of additions and therefore is expected to be least influenced by them.
- Type S: Here only N-terminal additions are added; a purification tag followed by a thrombin site, T7 tag, and enterokinase site (Fig. 1). Thrombin and enterokinase sites allow the cleavage of the N-terminal region, thus facilitating for the removal of the purification (and T7) tag, giving the possibility to generate an untagged protein. The T7 tag is an 11-residue peptide from the leader sequence of the T7 bacteriophage gene 10 (F. Studier & Moffatt, 1986) and aids protein expression in *E. coli*.
- Type D: Here both the N-terminal additions from type S plus C-terminal additions are added. On the C-terminus a sortase site (sortase A from *Staphylococcus aureus*) followed by a second purification tag is added (Fig. 1). The sortase site allows the removal of the C-terminal tag and the ligation of a peptide probe to the protein in a single reaction step (Popp & Ploegh, 2011). With this, a protein prenylation moity or other modification could be added. The C-terminal purification tag can be used for an additional affinity chromatography step during protein purification, and to separate labelled from unlabelled protein after a sortase-mediated labelling reaction. To asses C-terminal Cdc42 additions further, we designed a Cdc42 construct with an alternative membrane-binding domain (Cdc42-BC). Here the first two basic cluster (BC) regions of the yeast protein Bem1 (51 AA) got inserted into the protein’s C-terminus in between its PBR and CAAX box. The BCs are a 23 to 74 AAs long unstructured region of mostly positively charged AAs that were shown to anchor Bem1 to negatively charged membranes *in vitro* (Meca et al., 2019). We designed this construct to further asses how unstructured (linker) regions on Cdc42’s C-terminus affect its behaviour. We did not add structured regions, as the addition of a C-terminal amphipathic helix from Rit to Cdc42 resulted in folding issues of the protein *in vitro* (unpublished data by P. Schwille group (MPI Martinsried, Germany)). (The addition was in reference to Bendezu *et al*., which showed that this alternative membrane binding domain leads to viable cells *in vivo* (Bendezú et al., 2015).).

**Figure 1.**
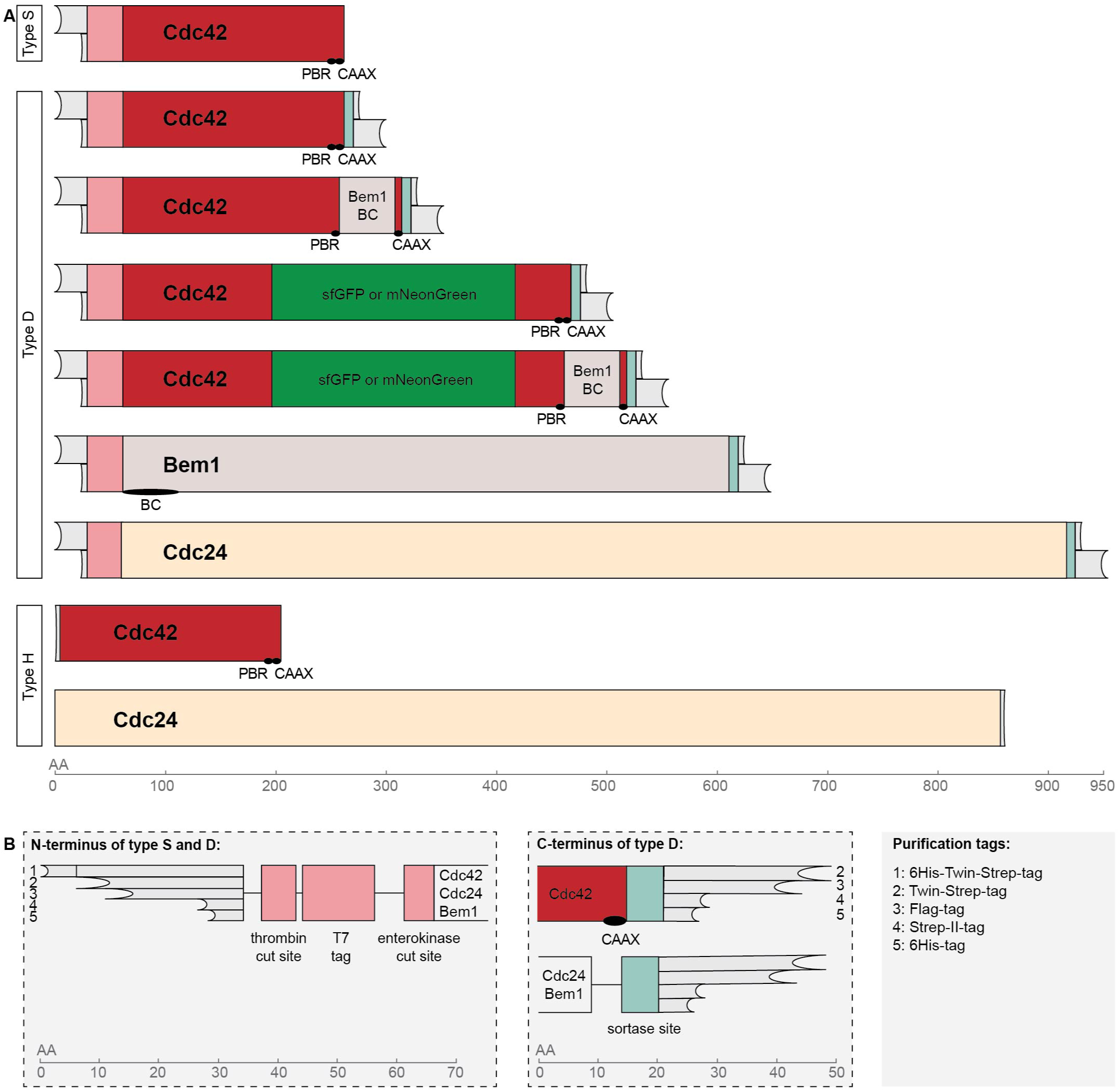
Schematic overview of the protein constructs. (A) Illustration of the general size and outline of double-tagged (type D) and single-tagged constructs (type S) and of constructs to which only a single 6His-tag got added (type H). (B) Zoom-in of the N- and C-terminal tag regions from type S and type D and illustration of the size difference of the purification tags. All constructs are drawn roughly up to scale, in terms of their number of amino acids (AA). The position of domains of interest are indicated. An overview of the specific constructs is given in Tab. 2.

As a standard we added a 6His-tag (H), placed N- and C-terminally, to all our constructs. The 6His-tag it is a small often-used tag that facilitates His-affinity chromatography (His-AC) - an easy, fast, and cheap purification method. Type D constructs contained additionally a Flag- (F), Strep-II- (S), or Twin-Strep- (SS) tag. All of these are also rather small in size and can be used for affinity chromatography. The Strep-II- and Twin-Strep-tag differ only in so far from each other as the Twin-Strep-tag is made from two repeats of the Strep-II-tag that are spaced with a linker (Tab. 1). The Twin-Strep-tag binds by an order of magnitude tighter to Strep-Tactin and is therefore more effective when used for purification purposes (Schmidt et al., 2013). Sequences and sizes of these purification tags are given in Tab. 1.

**Table 1.**
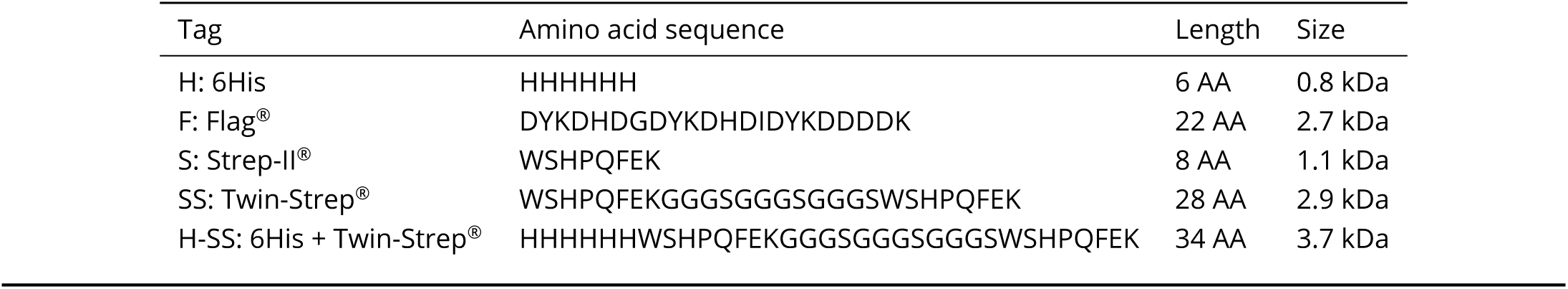
Purification tag specifications. For more information regarding the Strep-II^®^-tag and Twin-Strep^®^-tag see (Schmidt et al., 2013), and regarding the Flag^®^-tag see (Hopp et al., 1988).

Cdc42 expression in *E. coli* does not result in prenylated Cdc42, as *E. coli* lacks the machinery responsible for it. If needed, such a group can still be added in an *in vitro* reaction: Cdc42 constructs type H and S contain a C-terminal CAAX box, which can be prenylated using purified farnesyltransferase or geranylgeranyl-transferase type I (Golding et al., 2019). Type D constructs contain a sortase site with which a prenyl group can be ligated to the protein’s C-terminus (Popp & Ploegh, 2011).

To expand out investigations on Cdc42 further, we designed fluorescent Cdc42 fusions. In principle, a fluorophore could be attached to either Cdc42’s N- or C-terminus. Cdc42 fusions with N-terminal fluorophores have been controversial: N-terminal fusions of Cdc42 with fluorescent proteins have been shown to lead to disfunctional proteins *in vivo* (Bendezú et al., 2015), but two studies used such constructs successfully *in vitro* (Sartorel et al., 2018; Sonal et al., 2022). C-terminal additions on Cdc42 are not well-studied. We are only aware of the unsuccessful attempt to add a C-terminal amphipathic helix. However, it has been shown that fast-folding fluorophores can be inserted into a solvent-exposed loop of Cdc42 (Bendezú et al., 2015). We used this approach to create sandwich fusions of Cdc42 and sfGFP or mNeonGreen (Fig. 1), two proteins that are known to fold quickly (Pédelacq, Cabantous, Tran, Terwilliger, & Waldo, 2006; Shaner et al., 2013).

An overview of the herein used constructs and their abbreviations is given in Tab. 2.

**Table 2.** List of protein constructs and specification of their attributes. Protein sequences are stated in Supplement S6. If not stated otherwise, Cdc42 constructs contain the proteins natural CAAX sequence (CAIL). All ’pRV’ plasmids were created as part of this work. ^1^: pDM272 was received from D. McCusker (University of Bordeaux) and was published in (Rapali et al., 2017). Abbreviations: EK: enterokinase.

| Abbreviation | Type | N-term. tag | C-term. tag | Other modification | Plasmid | Protein size |
| --- | --- | --- | --- | --- | --- | --- |
| **Cdc42** |  |  |  |  |  |  |
| H-Cdc42 | H | 6His |  | CAAX sequence = CAIA | pRV035 | 22 kDa |
| H-Cdc42:CAIA | S | 6His |  | CAAX sequence = CAIA | pRV050 | 26 kDa |
| H-Cdc42:CTIS | S | 6His |  | CAAX sequence = CTIS | pRV086 | 26 kDa |
| H*-Cdc42:CTIS | S | 6His |  | TEV instead of EK site, CAAX sequ. = CTIS | pRV104 | 26 kDa |
| H-Cdc42-F | D | 6His | Flag |  | pRV007 | 29 kDa |
| F-Cdc42-H | D | Flag | 6His |  | pRV073 | 29 kDa |
| H-Cdc42-S | D | 6His | Strep-II |  | pRV024 | 28 kDa |
| H-Cdc42-SS | D | 6His | Twin-Strep |  | pRV068 | 29 kDa |
| S-Cdc42-H | D | Strep-II | 6His |  | pRV030 | 28 kDa |
| SS-Cdc42-H | D | Twin-Strep | 6His |  | pRV058 | 29 kDa |
| H-SS-Cdc42-F | D | 6His + Twin-Strep | Flag |  | pRV087 | 32 kDa |
| S-Cdc42-BC-H | D | Strep-II | 6His | + BC from Bem1 | pRV051 | 33 kDa |
| **Fluorescent Cdc42** |  |  |  |  |  |  |
| H-Cdc42-mNeon-F | D | 6His | Flag | + mNeonGreen<sup>SW</sup> | pRV010 | 55 kDa |
| F-Cdc42-mNeon-H | D | Flag | 6His | + mNeonGreen<sup>SW</sup> | pRV074 | 55 kDa |
| H-Cdc42-mNeon-S | D | 6His | Strep-II | + mNeonGreen<sup>SW</sup> | pRV052 | 54 kDa |
| H-Cdc42-mNeon-SS | D | 6His | Twin-Strep | + mNeonGreen<sup>SW</sup> | pRV067 | 56 kDa |
| S-Cdc42-mNeon-H | D | Strep-II | 6His | + mNeonGreen<sup>SW</sup> | pRV057 | 54 kDa |
| S-Cdc42-mNeon-BC-H | D | Strep-II | 6His | + BC from Bem1 + mNeonGreen<sup>SW</sup> | pRV093 | 61 kDa |
| H-SS-Cdc42-mNeon-F | D | 6His + Twin-Strep | Flag | + mNeonGreen<sup>SW</sup> | pRV090 | 58 kDa |
| F-Cdc42-sfGFP-H | D | Flag | 6His | + sfGFP<sup>SW</sup> | pRV101 | 57 kDa |
| H-Cdc42-sfGFP-SS | D | 6His | Twin-Strep | + sfGFP<sup>SW</sup> | pRV100 | 56 kDa |
| S-Cdc42-sfGFP-H | D | Strep-II | 6His | + sfGFP<sup>SW</sup> | pRV099 | 55 kDa |
| H-SS-Cdc42-sfGFP-F | D | 6His + Twin-Strep | Flag | + sfGFP<sup>SW</sup> | pRV102 | 59 kDa |
| **Cdc24 and Bem1** |  |  |  |  |  |  |
| Cdc24-H | H |  | 6His |  | pDM272 <sup>1</sup> | 98 kDa |
| H-Cdc24-SS | D | 6His | Twin-Strep |  | pRV065 | 103 kDa |
| S-Cdc24-H | D | Strep-II | 6His |  | pRV031 | 105 kDa |
| SS-Cdc24-H | D | Twin-Strep | 6His |  | pRV060 | 105 kDa |
| H-SS-Cdc24-F | D | 6His + Twin-Strep | Flag |  | pRV088 | 108 kDa |
| H-Bem1-F | D | 6His | Flag |  | pRV009 | 70 kDa |
| S-Bem1-H | D | Strep-II | 6His |  | pRV032 | 68 kDa |
| SS-Bem1-H | D | Twin-Strep | 6His |  | pRV061 | 70 kDa |
| H-SS-Bem1-F | D | 6His + Twin-Strep | Flag |  | pRV089 | 73 kDa |

### Cdc42 expresses robustly and can be purified in a high yield using His-AC

First, we examined expression levels of Cdc42. We assed Cdc42 constructs which only are 6His-tagged (type H), have a larger additional N-terminal region (type S), and contain both N- and C-terminal additions (type D) (Fig. 1). Cdc42 was placed under an IPTG inducible promotor so that it’s expression can be induced through addition of that chemical. We tested three expression conditions:

’f’: a strong and fast expression at elevated temperatures, induced by a high amount of IPTG (3 h at 37°C with 1 mM IPTG).

’s’: a low and slow expression at lower temperatures, induced by a smaller amount of IPTG (18 h at 18°C with 0.2 mM IPTG).

’AI’: a self-inducing combined approach, called auto-induction (3 h at 37°C + 18 h at 18°C) (F. W. Studier, 2005).

All Cdc42 constructs contained a 6His-tag, therefore expression levels could be analysed by anti-His Western blotting (Fig. 2a-c). Both type S and D expressed at all conditions in roughly equal amounts (Fig. 2a). The presence of a C-terminal tag or variations in Cdc42’s CAAX box did not alter the expression levels. Further, the position of the tags in type D did not affect Cdc42 expression (Fig. 2b). Type H did not express in any condition (Fig. 2a). The N-terminal region of constructs of type S and D, but not of type H, contains a thrombin cut site, T7 tag, and enterokinase cut site. If the direct placement of the 6His-tag onto Cdc42’s N-terminus interferes with protein folding, the additional elements could act as a spacer, circumventing the issue. The T7 tag could also directly be responsible for facilitating Cdc42 expression, as it aids protein expression in *E. coli*.

**Figure 2.**
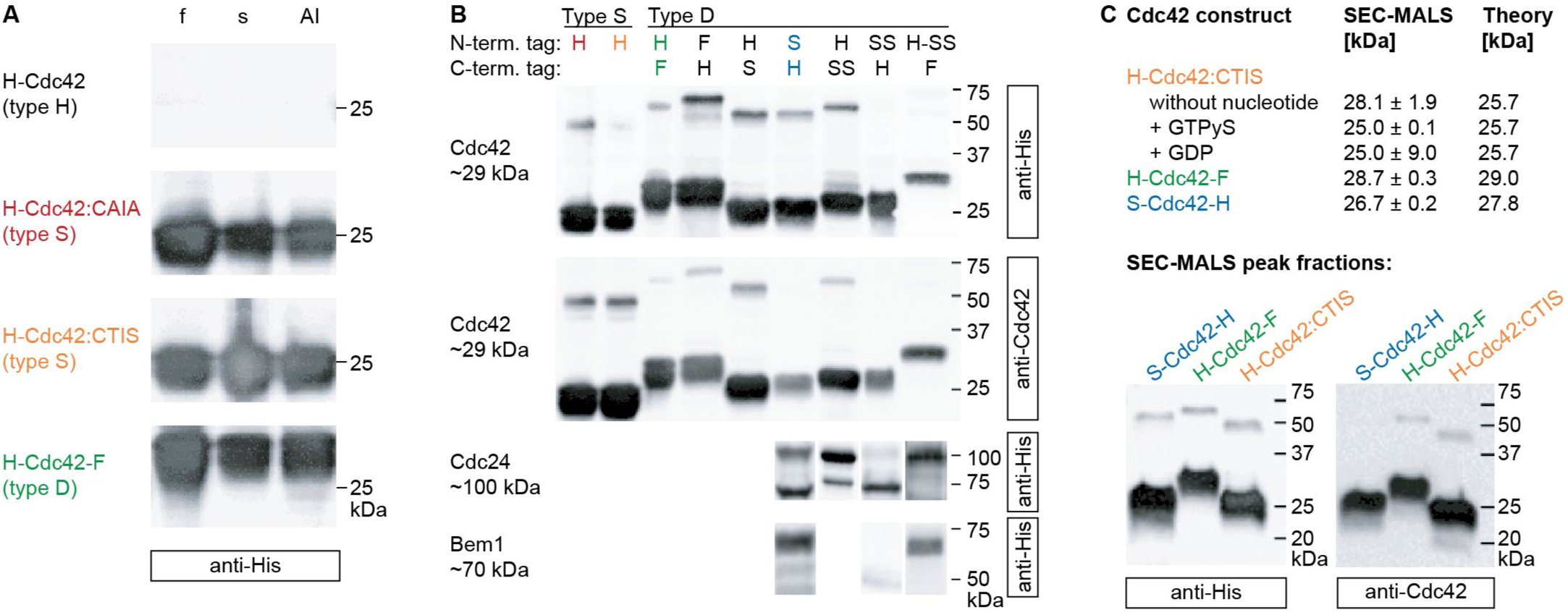
Cdc42 constructs type S and D, but not type H, express in a high amount and show dimer bands on SDS-Page. (A) Cdc42 constructs type S and D, but not type H, express in all three conditions (’f’: 3 h 37°C 1.0 mM IPTG, ’s’: 18 h 18°C 0.2 mM IPTG, ’ÁI’: 3 h 37°C + 18 h 18°C (F. W. Studier, 2005), Expression levels of Cdc42 were assessed by anti-His Western blotting. (B) Cdc42 expresses well, independent of which N- or C-terminal tags were added. Cdc24 and Bem1 with an N-terminal Twin-Strep-tag do not express. Expression levels were assessed by anti-His and anti-Cdc42 Western blotting. Cdc42 constructs were expressed in condition ’f’ and Cdc24 and Bem1 constructs were expressed in condition ’s’. (C) Cdc42 shows dimer bands on SDS-Page, but dimers can not be detected in SEC-MALS. Molecular weight of purified Cdc42 constructs determined by SEC-MALS in comparison to the expected mass based on the amino acid sequence (’Theory’). Anti-His and anti-Cdc42 Western blots of the peak fractions from SEC-MALS runs.

We then purified Cdc42 type S and D constructs. All constructs could be purified in a high yield in a single step His-AC (3-7 mg per 1 L expression volume) (Fig. 3). After purification only one construct, SS-Cdc42-H, was highly unstable and precipitated even at low concentrations (0.5 mg/mL). This was only the case for SS-Cdc42-H, all other constructs (including S-Cdc42-H, H-Cdc42-SS, and H-SS-Cdc42-F) showed no precipitation and were stable up to at least 2 mg/mL. This suggests that the N-terminal Twin-Strep-tag destabilises Cdc42. We observed a similar trend for the expression of two other yeast proteins; Cdc24 and Bem1. Cdc24 and Bem1 constructs, that were double-tagged in the same fashion as described for Cdc42, expressed when tagged with an N-terminal Strep-II-tag, C-terminal Twin-Strep-tag, or N-terminal 6His-Twin-Strep-tag, but did not, or only to greatly reduced level, express when tagged with an N-terminal Twin-Strep-tag (Fig. 2b). We suspect that the negative effect of the N-terminal Twin-Strep-tag is thus not Cdc42-specific, but inherent to the properties it exhibits when placed at a protein’s N-terminus.

**Figure 3.**
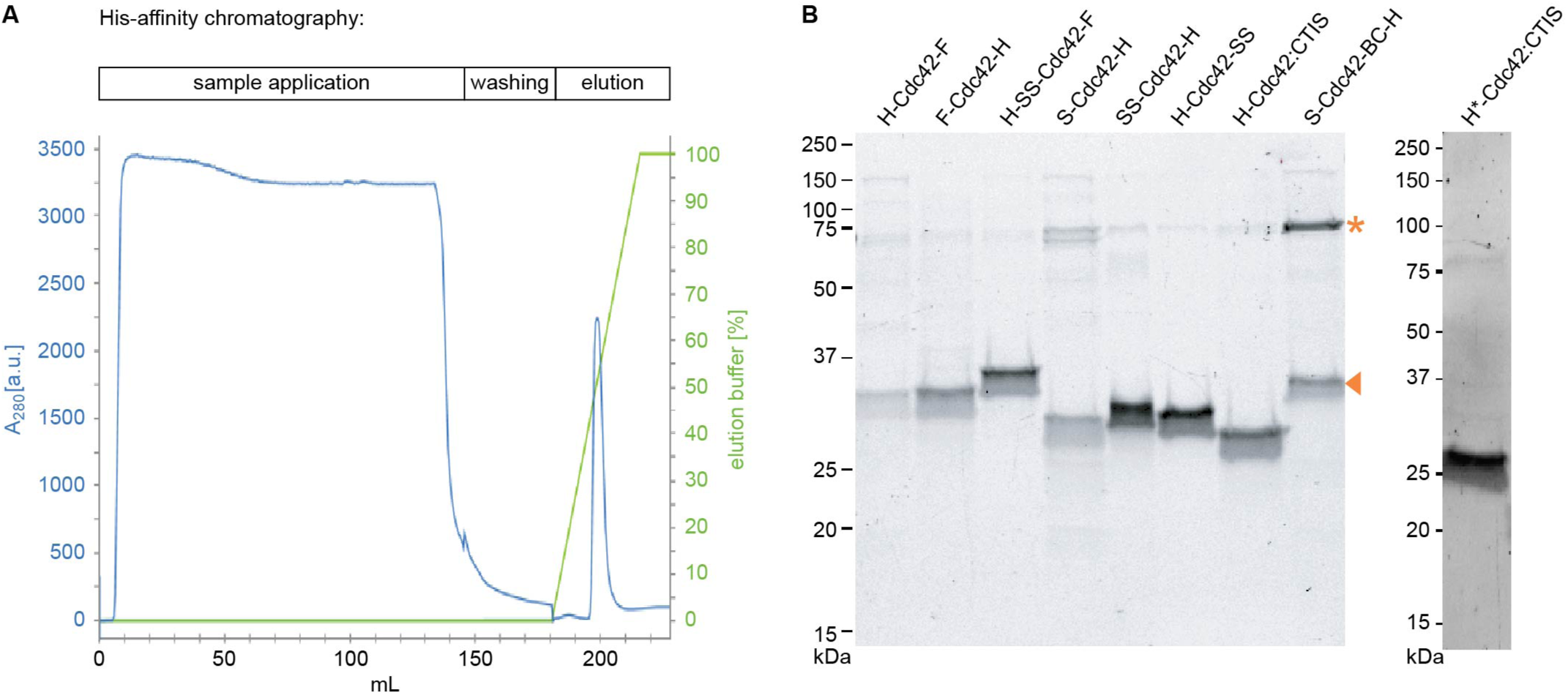
Cdc42 can be purified in a high yield in a single His-affinity chromatography step. (A) Purification profile of H-SS-Cdc42-F, purified by His-AC. The protein was eluted in a gradient of His-AC washing buffer (50 mM Tris-HCl (pH=8.0), 1 M NaCl, 5 mM imidazole, 1 mM 2-mercaptoethanol) and His-AC elution buffer (50 mM Tris-HCl (pH=8.0), 100 mM NaCl, 500 mM imidazole, 1mM 2-mercaptoethanol). The purification profiles of Cdc42 constructs shown in (B) looked alike. (B) SDS-Page of purified Cdc42 after a single step His-AC. The band of the full-length protein is indicated by an orange arrow. The asterix indicates the height of Cdc42 dimers.

We further observed that Flag-tagged protein constructs of type D run slightly higher than their expected size on SDS-Page (Fig. 2b, 3). We can imagine that this is an artefact of how the Flag-tag moves through an SDS-Page gel: the Flag-tag is filled with charged residues and contains very few hydrophobic residues. The lack of hydrophobic residues (and the presence of charged residues) makes it difficult for SDS to bind to the tag. This could allow the Flag-tag to fold back into its native structure. A folded structure will migrate differently through the gel, and in the case of the Flag-tag, slower that one would expect for a peptide of a similar size.

Taken together, Cdc42 expresses highly in a broad range of conditions and it can easily and in a high yield be purified in a single-step His-AC. The majority of constructs with N- and N-plus C-terminal additions (type S and D) behave equally well. The only exceptions were H-Cdc42 and SS-Cdc42-H. H-Cdc42 did not express at all, suggesting that a linker region between 6His-tag and Cdc42, or the T7 lead specifically, is required. The N-terminal Twin-Strep-tag caused Cdc42 precipitation and expression issues in Cdc24 and Bem1, indicating that the observed issues are tag-related.

### Cdc42 shows dimer bands on SDS-Page

We assessed Cdc42 expression levels of various constructs. Next to the expected Cdc42 band at 25-30 kDa, we observed a 50-60 kDa band for almost every construct in the anti-His and anti-Cdc42 Western blots (Fig. 2b). Its presence in the anti-Cdc42 Western blot confirms that this band is also indeed Cdc42, suggesting that Cdc42 forms dimers. Cdc42 belongs to the group of small GTPases, of which some have been shown to dimerise or oligomerise. Dimerisation has been linked to the PBR and (partial) removal of the PBR was shown to impede dimerisation. An illustration of this is given in Supplement S2 Tab. 1. Different GTPases have similar, yet slightly distinct PBR sequences. *S. cerevisiae* Cdc42 has the PBR sequence that has neither been linked to dimerisation nor to its absence. As the direct relation between the PBR sequence and protein dimerisation is still largely unknown, it is difficult to say, based on the PBR sequence, if Cdc42 dimerises. *In vivo* data suggests that it does not (Kang, Béven, Hariharan, & Park, 2010), but the absence of Cdc42 dimers *in vivo* does not necessarily exclude their existence *in vitro*: Cdc42 occupies a central role in the yeast cell cycle and has a lot of binding partners (Daalman et al., 2020), making it part of a lot of protein complexes. Cdc42 dimerisation was assessed though the fluorescence signal of two Cdc42-YFP fusions with YFP truncations: One Cdc42 copy was fused to the N-terminal part of YFP, the other Cdc42 copy to the C-terminal part. Fluorescence appears when both YFP fragments are brought together through association of two Cdc42 molecules (Kang et al., 2010). Hence, only dimeric Cdc42 leads to a YFP signal, monomeric Cdc42 and Cdc42 in hetero-complexes does not. If only a small fraction of Cdc42 dimers are Cdc42-Cdc42 homodimers (compared to Cdc42-effector protein heterodimers), not enough YFP signal is generated to observe these homodimers.

To further investigate this possibility, we ran purified Cdc42 sample on a size-exclusion chromatography (SEC) column and used multi-angle light scattering (MALS) to determine the molecular weight of the protein in each peak (Fig. 2c). Cdc42 ran in one peak and the SEC-MALS molecular weight corresponded to that of a monomer, independent of construct type or added nucleotide. We subsequently analysed the protein of these peak fractions by anti-His and anti-Cdc42 Western blotting. Again, the blots showed bands of the size of a dimer. Interestingly, the presence of dimers on SDS-Page was again influenced by the Strep-tag. Constructs with an N-terminal Twin-Strep-tag (SS-Cdc42-H, H-SS-Cdc42-F) formed no dimers (Fig. 2b), and S-Cdc42-H showed dimers in the anti-His, but not in the anti-Cdc42 Western blot (both in the expression test (Fig. 2b) and SEC-MALS samples (Fig. 2c)). C-terminal tags did not induce or influence dimer formation; type S constructs, and type D constructs with all possible C-terminal tags (6His, Flag, Strep-II, Twin-Strep) formed dimers (Fig. 2b). Thus, N-terminal Strep-II and Twin-Strep-tags seem to interfere with Cdc42 dimerisation under denaturing conditions. The origin of this remains elusive.

Taken together, we conclude that Cdc42 does not form stable long-lasting dimers *in vitro*. From this data we can not exclude that it forms transient and weakly bound dimers, as such complexes would not sustain themselves under the constant flow under which SEC is performed. Our data shows that it dimerises under denaturing conditions. This can be an artefact that has no translation to the behaviour of the folded protein, or mean that Cdc42 has some, still unexplored, potential to dimerise.

### Cdc42-mNeongreen^*SW*^/-sfGFP^*SW*^shows a more varied expression and purification behaviour than Cdc42

We explored the expression and purification behaviour of fluorescent Cdc42 sandwich fusions. Double-tagged Cdc42 was mostly unaffected by its specific N- and C-terminal tags (with the exception of an N-terminal Twin-Strep-tag). We assessed if this was also true for sandwich fusions of Cdc42 and sfGFP or mNeonGreen (Bendezú et al., 2015). We conducted expression tests (condition ’f’ and ’s’) and analysed them by anti-His Western blotting (Fig. 4). We tested combinations of N- and C-terminal 6His-, Flag-, Strep-, and C-terminal Twin-Strep-tags. We did not conduct tests with constructs with an N-terminal Twin-Strep-tag, as they already lead to issues with Cdc42 alone.

**Figure 4.**
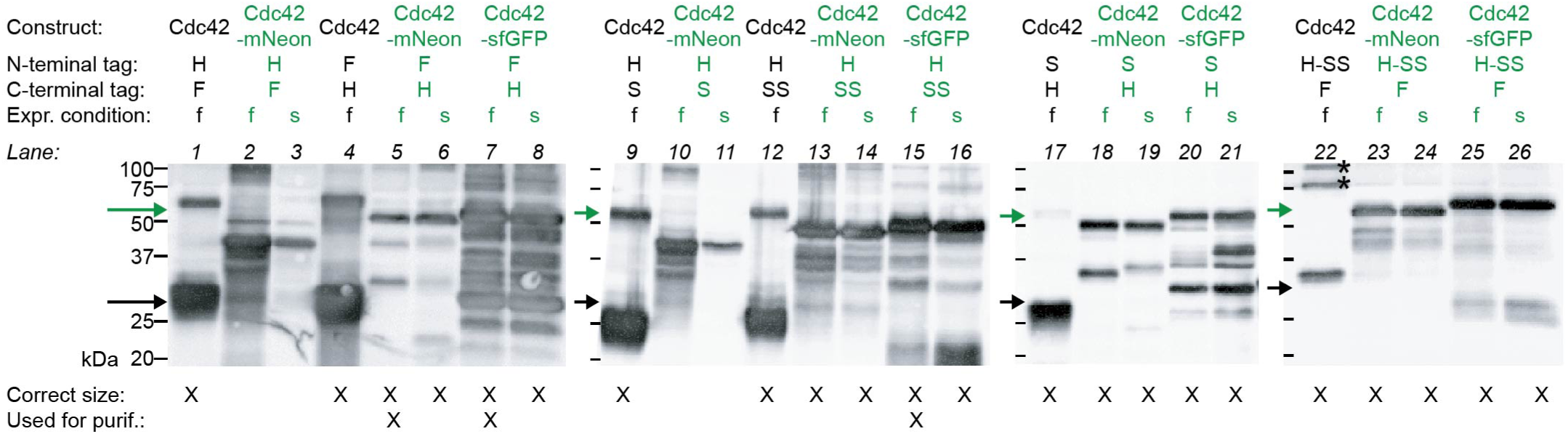
Most Cdc42-sfGFP^*SW*^ and -mNeonGreen^*SW*^ constructs express, but all show degradation. Expression levels (’f’: 3 h 37°C 1.0 mM IPTG, ’s’: 18 h 18°C 0.2 mM IPTG) were analysed by anti-His Western blotting. The black arrow indicates the size of monomeric Cdc42 and the green arrow indicates that of the fluorescent Cdc42 fusions. Bands annotated with an asterisk (lane 22) are not from the protein in that lane, but bands of the protein standard that migrated to this lane during the blotting process.

Most fusion-constructs showed bands at the expected size of 55 kDa (Fig. 4, green arrow). Only H-Cdc42-mNeon-F and H-Cdc42-mNeon-S did not express the full-size protein but only parts of it (Fig. 4 lane 2, 3, 10, 11). This is surprising as their Cdc42 equivalents showed no expression problems at all (Fig. 4 black arrow, lane 1 and 9). In contrast to Cdc42, most Cdc42-mNeon^*SW*^ /-sfGFP^*SW*^ fusions showed additional lower bands, that can originate from degradation or translation processes that terminated prematurely. This is not surprising, as the fluorophore sequence was inserted directly into the Cdc42 sequence, potentially making the fusion more susceptible to degradation.

In most cases both Cdc42-mNeonGreen^*SW*^ and Cdc42-sfGFP^*SW*^ fusions expressed with roughly the same amount and to same level of degradation, through significantly less strong than Cdc42. An exception is F-Cdc42-sfGFP-H (Fig. 4 lane 7 and 8), that expressed at a much higher amount than its mNeonGreen equivalent (Fig. 4 lane 5 and 6). It also shows higher amounts of degradation, even though it is possible that the degradation bands are only more pronounced due to the generally higher expression levels.

In most cases no big difference between the slow (’s’) and fast (’f’) expression condition could be observed; the fast condition lead to slightly more protein than the slow condition, but also showed a proportionally higher amount of degradation bands. An exception is S-Cdc42-sfGFP-H (Fig. 4 lane 20 and 21) where less degradation is present in the fast compared to the slow condition. The opposite is true for the mNeonGreen equivalent (Fig. 4 S-Cdc42-mNeon-H, lane 18 and 19). Here the slow condition expressed roughly the same amount of protein with less degradation.

In contrast to Cdc42, the expression behaviour of fluorescent Cdc42 fusions is very heterogeneous. Full-size protein can be produced with several, but not all, N- and C-terminal tag combinations. Degradation bands are observed in all cases but to different degrees, depending on the combination of (1) fluorophore used (sfGFP or mNeonGreen), (2) expression condition (’f’ or ’s’), and (3) N- and C-terminal tags. From our data no apparent relation between these factors could be drawn.

Next, we purified a subset of the constructs used in the expression test. We chose the constructs based on our applications and did not do entire screens. The data shown here therefore acts as exemplary evidence. We purified F-Cdc42-sfGFP-H, H-Cdc42-sfGFP-SS, F-Cdc42-mNeon-H, and S-Cdc42-mNeon-BC-H.

F-Cdc42-sfGFP-H could be purified in a high yield (5 mg per 1 L expression volume) in a single step His-AC (Fig. 5). The other constructs required multiple chromatography techniques and gave a very low yield (<0.1 mg per 1 L expression volume). Addition of detergents to the buffers increased the yield, but not to a substantial degree. We suspect that the high expression levels of F-Cdc42-sfGFP-H, which are already visible in the expression test (Fig. 4 lane 7,8), lead to the high yield. An extended discussion of the purification of these constructs is given in Supplement S3.

**Figure 5.**
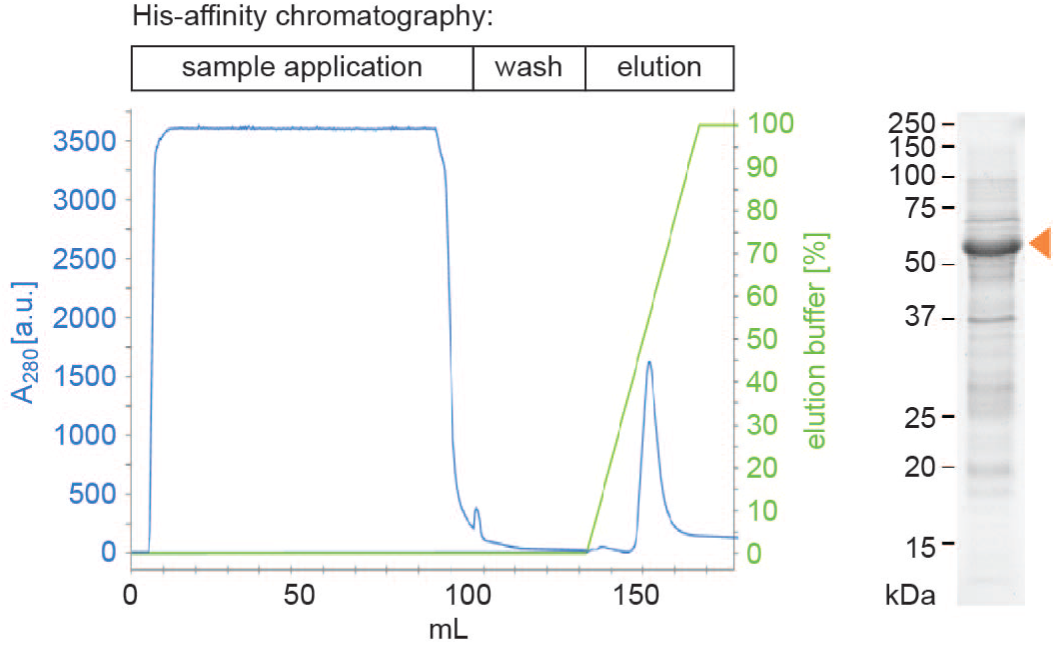
F-Cdc42-sfGFP-H can be purified in a single His-affinity chromatography step. The protein was eluted in a gradient of His-AC washing buffer (50 mM Tris-HCl (pH=8.0), 1 M NaCl, 5 mM imidazole, 1 mM 2-mercaptoethanol) and His-AC elution buffer (50 mM Tris-HCl (pH=8.0), 100 mM NaCl, 500 mM imidazole, 1mM 2-mercaptoethanol). SDS-Page of the elution peak, the band of the full-length protein is indicated by an orange arrow.

We further observed for all constructs that the lysis step was incomplete (about 30-50% of cells lysed), even though the same protocol resulted in 100% lysed cells if a non-fluorescent Cdc42 construct was used. The lysis could be slightly improved when the number of lysis rounds with the high pressure homogenizer were doubled from five to ten. The origin of this remains elusive.

Based on this data, we suggest to use F-Cdc42-sfGFP-H due to its easy purification procedure which results in a high yield. If another construct is required, we advise to screen for one that has the highest yield after His-AC, as we found this to be the limiting factor.

### Cdc42’s GTPase activity and interaction with the GEF Cdc24 can be reliably assessed using GTPase-Glo assays

Cdc42 is a small GTPase and can therefore hydrolyse GTP. To test how active the various Cdc42 constructs are, we performed GTPase assays using the Promega GTPase Glo^TM^ assay. Here serial dilutions of Cdc42 were incubated with GTP for a certain time, after which the reactions were stopped and the amount of remaining GTP was measured (see materials and methods). To compare the GTPase activity of different Cdc42 constructs, we determined GTP hydrolysis cycling rates *k*. These rates encompass the entire GTPase cycle, which can be described in three steps (Fig. 6a): (1) Cdc42 binds to a free GTP. (2) GTP gets hydrolysed by Cdc42. (3) Cdc42 releases GDP.

**Figure 6.**
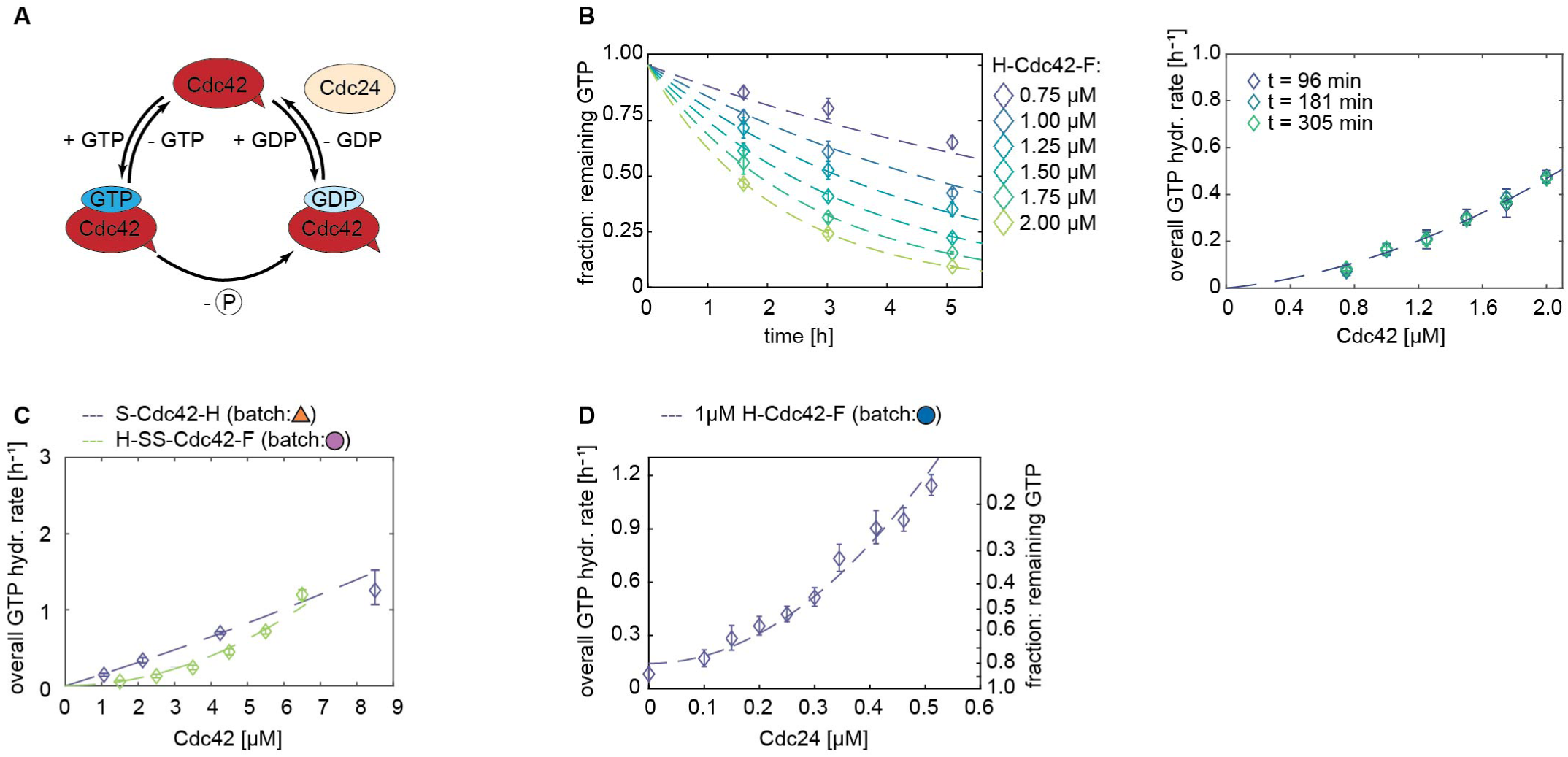
Cdc42’s GTPase activity and interaction with the GEF Cdc24 can be quantified using GTPase Glo assays (Promega). (A) Schematic illustration of the GTPase cycle. (B) GTPase Glo assay: The amount of remaining GTP declines exponentially over time. Dashed lines are fits (Eq. 1). (C) H-SS-Cdc42-F shows cooperative behaviour (high *k*_2_ and *k*_1_ ≈ 0) and S-Cdc42-H exhibits almost no cooperativity (high *k*_1_ and *k*_2_ ≈ 0). (D) The overall GTP hydrolysis cycling rate scales non-linearly with the Cdc24 concentration.

If one construct shows decreased rates *k* it indicates that at least one of these steps is occuring at a slower rate, suggesting that this batch has less active Cdc42 or a higher ratio of inactive to active protein. If batches of a certain construct show this behaviour repeatedly, the N- and/or C-terminal additions are likely interfering with its functionality.

First, we examined how the GTP concentration changes for different Cdc42 concentrations over time. We performed experiments to measure the amount of remaining GTP after incubation times of 1.5 h, 3 h, and 5 h. The graph of the amount of remaining GTP over time (Fig. 6b) shows that the GTP hydrolysis process can be described by an exponential decline (Eq. 1). The assay data of all three incubation times gave the same fit (Fig. 6b), suggesting that one data set from one assay, i.e. one incubation time, is sufficient. In this assay we used Cdc42 concentrations of 0.75 to 2.00 µM. For assays comparing the GTPase activity of various Cdc42 constructs we used wider concentration ranges (1 - 9 µM Cdc42 in combination with incubation times of 1 -1.5 h) to ensure a better fit quality.

We determined the GTP hydrolysis cycling rates of Cdc42 using

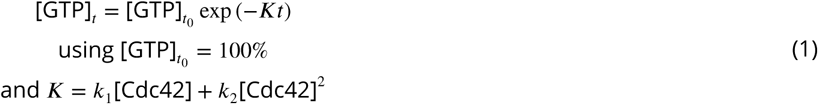

We will refer to *K* as the ‘overall GTP hydrolysis cycling rate’ in the following. In this equation *k*_1_ describes the GTP hydrolysis cycling rate of a single Cdc42 molecule and *k*_2_ includes any effects due to Cdc42 dimerisation or cooperativity. Introduction of this two-parameter fit was needed as some Cdc42 constructs showed cooperative behaviour (high *k*_2_ and *k*_1_ ≈ 0), whereas others showed almost none (high *k*_1_ and *k*_2_ ≈ 0) (Fig. 6c).

Next to Cdc42’s GTPase activity, we also investigated its interaction with the GEF Cdc24. As a GEF Cdc24 boosts the release of GDP from Cdc42 (GTPasy cycle step (3), Fig. 6a) and thereby increases the cycling speed of the GTPase cycle.

Assays with Cdc24 dilutions show that Cdc24 is highly active; addition of sub-µM concentrations of Cdc24 to 1 µM Cdc42 were sufficient to boost the reaction cycle by a factor of about three (Fig. 6d).

Further, the data showed that the GTP decline does not depend linearly on the Cdc24 concentration, but is better approximated by a quadratic term. This suggests cooperativity. Previous work showed that Cdc24 has the capability to oligomerise via its DH domain (Mionnet, Bogliolo, & Arkowitz, 2008). We expect dimers and oligomers to have an increased GEF activity. This could, for example, be facilitated by Cdc24’s C-terminal PB1-domain, which has been suggested to reduce Cdc24’s GEF activity through intramolecular interactions (Shimada, Wiget, Gulli, Bi, & Peter, 2004). Cdc24 oligomerisation could interfere with this self-interaction and thereby increase the proteins GEF activity. We note that previous this assumption could be seen to contradict previous *in vitro* work: Cdc24 peptides, consisting only of Cdc24’s DH and PH domain, showed that these peptides exhibit GEF activity. The GEF activity was not changed when oligomerisation was inhibited (through mutations) or amplified (Mionnet et al., 2008). These findings would exclude that Cdc24 oligomers exhibit an increased GEF activity. However, one has to be careful when applying these findings to full-size Cdc24: For one, not full size protein, but only peptides were used. Other domains that are not directly involved in oligomerisation or GEF funtion can still affect these properties. For example, the PB1 domain was suggested to reduce Cdc24 GEF activity in a self-inhibitory fashion (Shimada et al., 2004). Next, samples representing a heightened oligomerisation state were produced through addition of an additional oligomerisation domain to the peptides. This domain was not related to Cdc24 and could be triggered to oligomerise through the addition of a chemical. It is thus questionable if the GEF activity of these oligomerised peptides relates to the GEF activity of oligomeric Cdc24.

To determine Cdc42-Cdc24 interaction rates *k*_3_, we incubated Cdc42:Cdc24 mixtures, as well as samples containing only Cdc42, with GTP for 1-1.5 h and measured the amount of remaining GTP. We had observed that rates for Cdc42 can vary slightly between assays (Fig. 7a). To account for this variance, we introduced the parameter *c*_*Cdc42orr*_, that maps all factors that lead to variations between assays onto the Cdc42 concentration. The assay data, including samples containing only Cdc42 and Cdc42 - effector protein mixtures, was fitted with

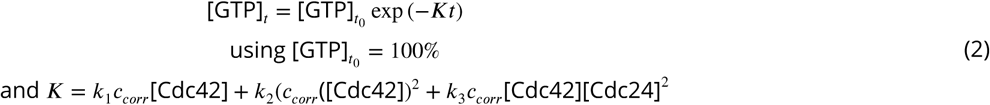

to determine *c*_*corr*_ and *k*_3_ (using *k*_1_ and *k*_2_ values determined earlier (Eq. 1)). Values of *c*_*corr*_ are 0.7 - 1.6, with the majority close to 1.0 (see Supplement S4).

**Figure 7.**
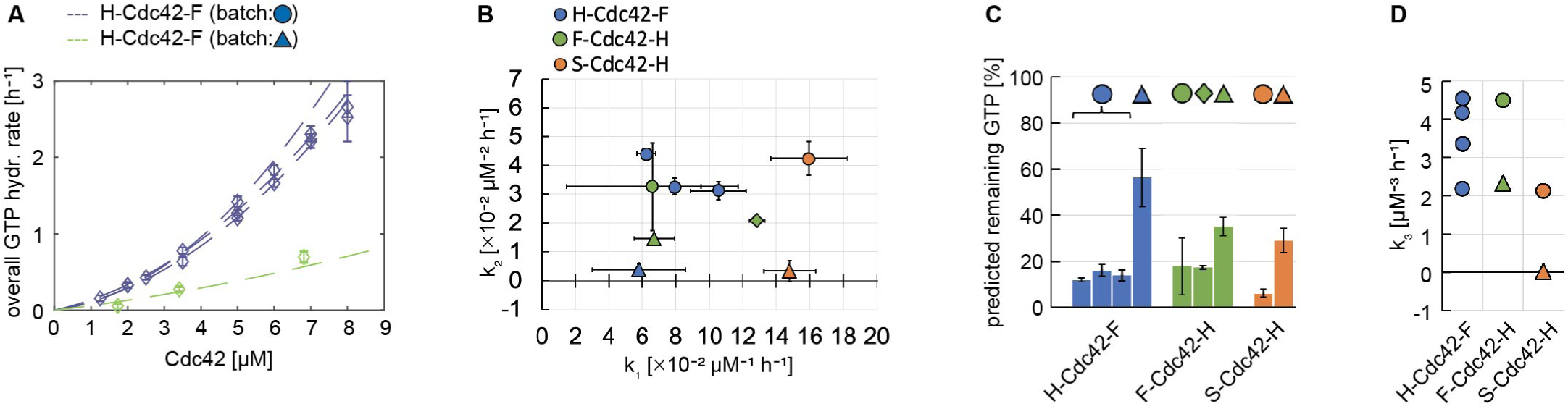
Cdc42 purification batches can show variations in their GTPase activity and GEF interaction or be non-functional. (A) Seperate GTPase assays show a similar GTPase activity for Cdc42, but purification batches can vary in their activity. (B) GTP hydrolysis cycling rates *k*_1_ and *k*_2_ for different Cdc42 constructs and purification batches. (C) Predicted amount of remaining GTP for Cdc42 constructs (5 µM) after an incubation time of 1.5 h, calculated using rates *k*_1_, *k*_2_ shown in (B). Symbols indicate if rates of a specific purification batch were used. (D) Cdc42-Cdc24 interaction rates *k*_3_. (A-D): GTP hydrolysis cycling rates *k*_1_ and *k*_2_ and Cdc24 interaction rates *k*_3_ for different Cdc42 constructs. Data points of the same colour refer to the rates obtained from separate measurements/assays. Data points of the same colour but a distinct shape (circle, triangle, rhombus) represent different purification batches of the same construct. Summary graphs of the data, values, and overview figures are given in Supplement S4.

We used these assays to examine the variation (1) within assays, (2) between purification batches of the same Cdc42 construct, and (3) between Cdc42 constructs.

#### The variation between assays is generally small

We conducted several assays with H-Cdc42-F. In all assays the overall GTP hydrolysis cycling rate *K* increases similarly with Cdc42 concentration; data points of fits from three assays overlay (Fig. 6a). Variations observed can be due to small concentration differences introduced though pipetting of small volumes (as are required for this assay), small temperature and shaker speed fluctuations during the GTPase reaction, and/or intrinsic changes in the Cdc42 protein due to other external conditions. Fitting each measurement data with *k*_1_ and *k*_2_ results in values that vary a bit (Fig. 7b, blue dots). However, they describe a similar GTPase behaviour: We used the rates and Eq. 1 to calculate how much GTP would remain in a simulated GTPase assay with equimolar amounts of Cdc42. All three *k*_1_ and *k*_2_ pairs lead to similar amounts of predicted remaining GTP (Fig. 7c, blue bars with blue dot). We assume that the variation in *k*_1_ and *k*_2_ is due to fitting quality. Measurements using more data points, or restricting the fit to either *k*_1_ (no cooperativity) or *k*_2_ (cooperativity) would result in less variation. As the aim of this assay was to compare different constructs, and as constructs showed a heterogeneous tendency towards cooperativity (Fig. 6c), we considered the observed variations in *k*_1_ and *k*_2_ unideal but necessary. We will use them as a baseline variability to compare purification and construct variations with.

We further used H-Cdc42-F to asses the variability of the Cdc42-Cdc24 interaction rate *k*_3_ between assays. Variations in *k*_3_ were bigger than those in *k*_1_/*k*_2_: For H-Cdc42-F, values of 2.2 to 4.5 µM^−3^ℎ^−1^ were observed (Fig. 7d, blue dots). As Cdc24 boosts the GTPase cycle in a strong fashion that depends non-linearly (quadratically in our fits) on its concentration (Fig. 6d), small activity changes of Cdc24 and dilution inaccuracies between assays affect *k*_3_ quadratically. For example, if the intended Cdc24 concentration was 0.2 µM, but in fact 0.1 µM was present in the assay, *k*_3_ will decrease by a factor of four. To accurately determine *k*_3_ one thus needs to perform several assays. To compare different Cdc42 constructs, one ideally would add them all to the same assay. If this is not feasible, addition of a reference Cdc42 construct to all assays is advantageous, as then *k*_3_ values can be normalised/compared to *k*_3_ of the reference construct. If this was not done, the degree of *k*_3_ variation needs at least to be considered. Constructs can then only be considered to have a stronger/weaker GEF interaction, if their *k*_3_’s are significantly above or below the observed variation.

#### Some purification batches are not functional

We assessed two purification batches for H-Cdc42-F and S-Cdc42-H, and three purifications for F-Cdc42-H (Fig. 7b-d, indicated by distinct symbols of the same colour). All batches were expressed and purified using the same protocol, and we expected to see only minor differences.

Surprisingly, one purification batch of H-Cdc42-F showed a hugely reduced GTPase activity (Fig. 7a). We assume that this batch contained either protein with a reduced activity or a substantial portion of inactive protein. We discarded this batch from further analysis.

One of three purification batches of F-Cdc42-H had a reduced GTPase activity (Fig. 7b,c, green triangle, and Supplement S4). It was less severe than the reduction observed for H-Cdc42-F. Its GEF interaction rate *k*_3_ was also similar to that of the more active F-Cdc42-H batch (Fig. 7d), given the observed variability of *k*_3_. We conclude that this batch is a bit less active that it ideally could be, but still shows sufficient activity. We also observed a reduced GTPase activity for one purification of S-Cdc42-H (Fig. 7b,c, orange triangle, and Supplement S4), and the reduction was similar to that in F-Cdc42-H. In contrast to F-Cdc42-H, this Cdc42 batch did not interact with Cdc24 (Fig. 7d), suggesting that Cdc42’s GTPase activity is independent from its ability to interact with Cdc24. One can thus not deduce full protein functionality from GTPase activity data alone.

The decrease in GTPase activity of certain purifications is visible in this data set as a drop in *k*_2_: *k*_1_ of the less active batch is within error of the more active batch, but *k*_2_ drops close to zero (Fig. 7b). A lack of cooperative behaviour could explain the reduced overall activity; Cdc42 molecules in these batches would still possess their baseline GTPase activity, but lack the ability to interact and enhance each others GTPase activity, thus leading to an overall reduced activity. On the other hand, the observed drop in *k*_2_ could also be a fitting artefact due to the decreased Cdc42 activity; a drop in activity in the used Cdc42 concentration range can make the data appear more linear than it actually is (Supplement S1). Use of a wider concentration range for these constructs would show if they have indeed less cooperative behaviour or if the perceived drop is an artefact. The data we collected is limited and of exemplary nature, and further experiments are required to claim that is is indeed the case.

#### All Cdc42 constructs, with the exception of SS-Cdc42-H, are active

Finally, the different constructs can now be compared. GTPase data (Supplement S4) were fitted to determine GTP hydrolysis cycling rates *k*_1_ and *k*_2_ (Fig. 8a) and GEF interaction rates *k*_3_ (Fig. 8c). The amount of remaining GTP in a simulated GTPase assay with equimolar amounts of each construct using rates *k*_1_ and *k*_2_ is shown in Fig. 8b.

**Figure 8.**
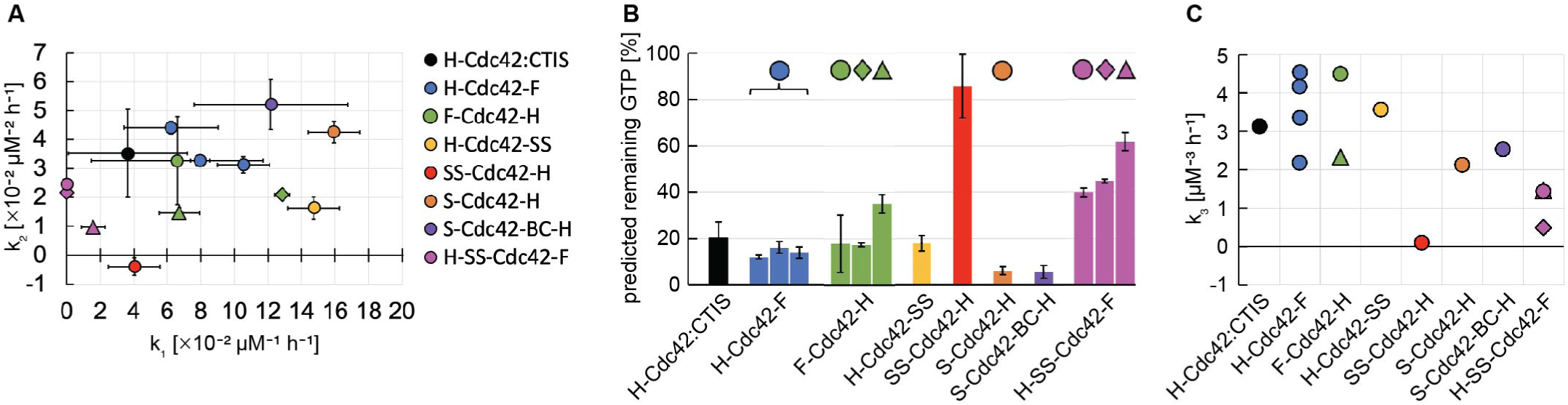
All Cdc42 constructs of type D, with the exception of SS-Cdc42-H and H-SS-Cdc42-F, show a similar GTPase activity and GEF interaction. (A): GTP hydrolysis cycling rates *k*_1_ and *k*_2_ of different Cdc42 constructs. Data points of the same colour refer to the rates obtained from separate measurements/assays. Data points of the same colour but a distinct shape (circle, triangle, rhombus) represent different purification batches of the same construct. (B): Predicted amount of remaining GTP for Cdc42 constructs (5 µM) after an incubation time of 1.5 h, calculated using rates *k*_1_, *k*_2_ shown in (A). Symbols indicate if rates of a specific purification batch were used. (C) Cdc42-Cdc24 interaction rates *k*_3_. (A-C): Summary graphs of the data, values, and overview figures are given in Supplement S4.

Most double-tagged constructs (type D: H-Cdc42-F, F-Cdc42-H, H-Cdc42-SS, S-Cdc42-H, S-Cdc42-BC-H) show similar GTPase activities and a weak cooperativity (both *k*_1_ and *k*_2_ are unequal to zero) (Fig. 8a,b), and GEF interaction rates that fall within the assay variability observed for H-Cdc42-F (Fig. 8c). Both GTPase activity as well as GEF interaction are similar to the only N-terminally tagged H-Cdc42:CTIS (type S). Even S-Cdc42-BC-H, which contained further C-terminal modifications, does not show any reduced activity (Fig. 8). This suggests that C-terminal additions do not hinder Cdc42’s functionality (GTPase activity and GEF interaction). SS-Cdc42-H has almost no GTPase activity and does not interact with Cdc24 (Fig. 8 red dots/bar). SS-Cdc42-H showed already precipitation issues during the purification process and this assay confirms that the protein is barely functional. (We show only one data point/one assay for this construct. However, several additional assays were performed to ensure that the observed behaviour is not an artefact. The same behaviour was observed repeatedly and in most assays no GTPase activity was observed. The rates could only be extract from one assay where a significantly longer incubation time was used.) H-SS-Cdc42-F also has a lower GTPase activity and a lower GEF interaction (0.5 - 1.5 µM^−3^ℎ^−1^) (Fig. 8, purple symbols). It is more active than SS-Cdc42-H but less than the other constructs. This is true for all three purification batches tested. In the same assay that we used to assess the GEF interaction of H-SS-Cdc42-F, we had also added H-Cdc42-F as a control. In this assay H-Cdc42-F had a GEF interaction rate of 4.5 µM^−3^ℎ^−1^. The low *k*_3_ of H-SS-Cdc42-F is thus not due to assay variation, but describes a reduced or weaker interaction with Cdc24. This data shows that SS-Cdc42-H is not active and that H-SS-Cdc42-F has a reduced activity. The 6His-tag upstream of the Twin-Strep-tag seems to restore Cdc42 functionality thus partially, but not fully.

Taken together, our data shows that the GTPase activity of Cdc42 can be assessed with the GTPase Glow assay (Promega). The variation between assays containing only Cdc42 is small. Cdc42 is heterogeneous in its tendency cooperativity, and a lack of any cooperativity might be a potential indication for a less functional protein. We observe a stronger assay variation for assays assessing the Cdc42 Cdc24 interaction. To effectively compare different Cdc42 constructs, one thus has to account for this assay variation through assessing it, adding a reference to every assay, or by only comparing constructs, which rates were measured in the same assay. We further see that not all purification batches give fully functional protein and that constructs can have a GTPase activity but show no GEF interaction. Thus, every purification batch has to be assessed for its activity. Our data is not sufficient to provide quantitative measures that aid the decision process of if a construct is sufficiently active, but acts as exemplary evidence to guide the process and to highlight outcomes that can be observed. Our data suggests that mostly unstructured C-terminal additions to Cdc42 do not interfere with its GTPase activity and GEF interaction, but that the N-terminal Twin-Strep tag does. Placement of a 6His-tag prior to the Twin-Strep-tag partially, but not fully, restores Cdc42 functionality.

### Pulldown assays are less robust to study weakly bound Cdc42 effector interactions as the Cdc42-Bem1 complex

We then wanted to explore other Cdc42 - effector interactions. The GTPase assay worked well to investigate the Cdc42 Cdc24 interaction, but is limited to only those interactors of Cdc42 that affect its GTPase cycle. Pulldown assays, which directly assess protein - protein binding, can be used to study a wider range of interactions. Here protein A is tethered to a matrix using its affinity tag (e.g. a Flag-tag). Protein B, which does not contain this affinity tag, is added. The matrix is washed several times. Then protein A is eluted. If protein B is in this elution as well, it suggests that it is bound to protein A (Fig. 9a). We conducted Flag-pulldown experiments with Bem1 and Cdc42, using Flag-tagged Bem1 (H-Bem1-F) and Cdc42 constructs that do not contain a Flag-tag.

**Figure 9.**
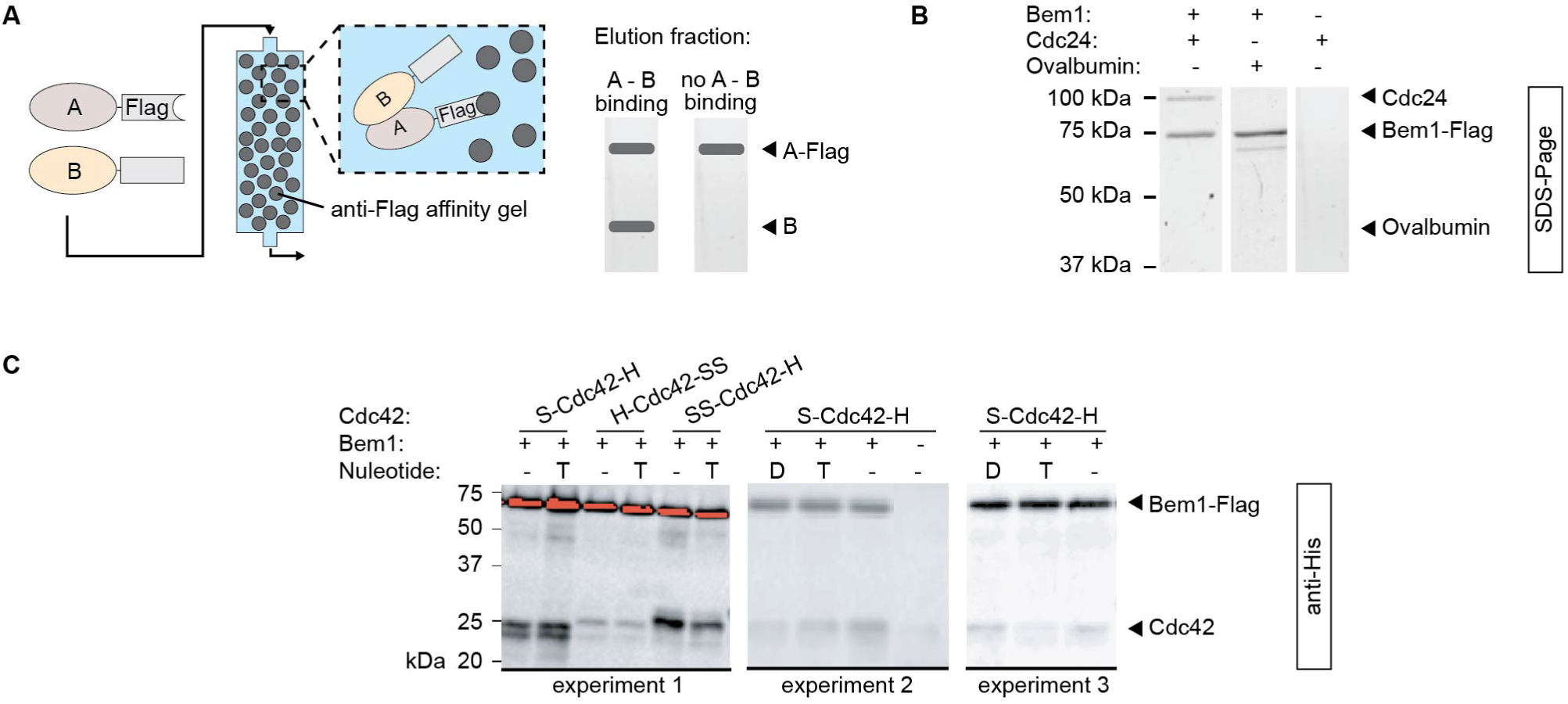
Flag-pulldown experiments are suitable to study strongly binding proteins (Cdc24-Bem1), but less reliable for weakly interacting proteins (Cdc42-Bem1). (A) Schmematic illustration of Flag-pulldown experiments. (B) SDS-Page of elution fractions of Flag-pulldown experiments with H-Bem1-F, Cdc24-H, and Ovalbumin. (C) Anti-His Western blots of elution fractions of three Flag-pulldown experiments with H-Bem1-F and Cdc42 preloaded with no nucleotide, GDP (D), or GTPγS (T).

We observed that binding between Bem1 and Cdc42 occurred, but that it was so weak that Cdc42 could only be detected using Western blotting (Fig. 9c). In comparison, pulldown experiments with the same Bem1 construct and Cdc24, another binding partner of Bem1, showed so high amounts of Cdc24 in the elution fraction that it was visible on SDS-Page, indicating stronger binding. Cdc24 alone did not bind to the anti-Flag affinity gel. The Bem1 - Cdc24 interaction was also specific: Ovalbumin, a protein not interacting with Bem1, was not present in the elution fraction (Fig. 9b).

We used several Strep-tagged Cdc42 versions (Fig. 9c): S-Cdc42-H, H-Cdc42-SS, and SS-Cdc42-H. All of these Cdc42 constructs bound to Bem1. The blotting signal for H-Cdc42-SS was weaker in comparison to S-Cdc42-H and SS-Cdc42-H. This is surprising, as we expected the barely functional SS-Cdc42-H to not, or only very weakly, bind to Cdc42. SS-Cdc42-H either still shows some remaining functionality (ability to bind to Bem1), or all our binding events are due to nonspecific interactions (and thus only detectable with Western blots). We tested if Strep-tagged Cdc42 (S-Cdc42-H) would also stick to the anti-Flag affinity gel, thus causing the presence of Cdc42 in all elution samples. In absence of H-Bem1-F, a small amount of S-Cdc42-H was binding to the anti-Flag affinity gel. Its signal in the Western blot however was significantly weaker than when Cdc42 was incubated with anti-Flag affinity gel in presence of H-Bem1-F (Fig. 9c). Out data suggests that there is weak binding between Cdc42 and Bem1, but we can not exclude that this is due to non-specific interactions between both proteins. In order to test this, one would need to add another control in which Bem1 and another His-tagged protein, that is not a binding partner of Bem1, are used. We did conduct this control due to the lack of a suitable His-tagged protein.

Bem1 was shown to specifically bind to GTP-bound, but not GDP-bound, Cdc42 (Bose et al., 2001). We conducted two separate Flag-pulldown experiments in which 1 nmol Cdc42 got pre-loaded with no nucleotide, 100 nmol GDP, or 100 nmol GTPγS (a non-hydrolysable variant of GTP). The intensities of the Western blot signal for Cdc42 varied with the nucleotide type, but also per experiment: In one experiment more Cdc42-GDP (than Cdc42-GTP or nucleotide free Cdc42) bound to Bem1, but in another experiment it was nucleotide-free Cdc42 (Fig. 9c). We thus could not observe a significant effect of the nucleotide state on the Cdc42-Bem1 interaction. This further points to the conclusion that something in our Cdc42 - Bem1 interaction assays is not properly functioning. We also spotted several small experimental differences between the pulldown assays shown here and those conducted by Bose *et al*.: Bose *et al*. conducted similar pulldown experiments as shown here, but used GST-tagged (26 kDa) instead of a Flag-tagged (3 kDa) proteins (and subsequently GST agarose instead of anti-Flag affinity gel). Given the size of the GST-tag, it can not be excluded that this tag is destabilising the Bem1-Cdc42 interaction, thereby exaggerating the effect of a conformational state of Cdc42 on Bem1 binding. Pulldowns with (1) GST-Cdc42 and Bem1, and (2) GST-Bem1 and Cdc42, both showed the same result. Thus, the GST-tag would need to be positioned in both cases in such a way that it destabilise the interaction, questioning this hypothesis. Another experimental difference is the way in which Cdc42 was locked in one nucleotide state: We pre-loaded Cdc42 with excess GDP or GTPγS, Bose *et al*. used Cdc42-mutants that are locked in a GDP- or GTP-state. One interpretation is that GDP/GTP-loaded Cdc42 and Cdc42 mutants are structurally distinct in a way that affects Cdc42-Bem1 binding. Alternatively, it could indicate that our pre-loading was insufficient and that it did not bring the majority of Cdc42 into its GDP-bound conformation.

In conclusion, we find that pulldown assays are a great tool to study the interaction of strongly interacting proteins. Here binding can be analysed by SDS-Page, and controls for non-specific binding can easily be added. In contrast, weaker protein - protein interactions, as the Cdc42 - Bem1 interaction, are more difficult to study using pulldown experiments. We observed a very weak, but potentially non-specific, interaction between Cdc42 and Bem1, that also did not depend on the nucleotide state of Cdc42 (contradicting (Bose et al., 2001). Western blots are very sensitive and can easily be overinterpreted. Controls to correct for signal from Cdc42 sticking to the matrix, and from non-specific protein-protein binding (Bem1 plus a protein that is not a binding partner of Bem1) are required. With sufficient preparation they can be a suitable tool, but are, in our experience, much more difficult and error-prone than GTPase assays.

### TEV protease, but not enterokinase, is suitable to cleave tags off Cdc42

We created Cdc42 constructs with both an N- and C-terminal tag (type D), and one which only contains an N-terminal tag (type S). The effect of C-terminal additions (sortase site and purification tag) can be determined by comparing type D with type S. To asses the the effect of the N-terminal additions (incl. purification tag), an Cdc42 version lacking that region needs to be created. To generate such a Cdc42 version, we used the existing enterokinase site in H-Cdc42:CTIS (type S) to cleave the N-terminal region. We conducted a condition screen varying amounts of enterokinase used, reaction time and temperature (Supplement S5 Fig. 1a). In all conditions the majority of the cleavage products are smaller than the expected product; bands at ∼14 kDa and ∼8 kDa show a stronger intensity than the band of the expected cleavage product at ∼20 kDa (Supplement S5 Fig. 1a). The enterokinase cleavage sequence is ‘Asp-Asp-Asp-Asp-Lys’, but it was reported that it can also cleave after a site of an acidic amino acid followed by a basic amino acid (Shahravan, Qu, Chan, & Shin, 2008; Light, Savithri, & Liepnieks, 1980). We found two sequences that fit those requirements and that can explain the observed fragments (Supplement S5 Fig. 2).

As enterokinase showed an undesirable cleavage behaviour, we designed another Cdc42 construct in which the enterokinase site was replaced with a TEV cleavage site (H*-Cdc42:CTIS). We tested two commercially available enzymes (TEV (1): TEV protease (Sigma Aldrich), TEV (2): AcTEV (Invitrogen)) and one in-house purified TEV protease (TEV (3): pSF1818). We again conducted a condition screen varying amounts of protease used, reaction time and temperature (Supplement S5 Fig. 1b). In all conditions only the desired cleavage product was observed. Increasing the amount of protease used resulted in more cleavage product in the case of TEV (1) and TEV (2), but not with TEV (3). Prolonged reaction times did only marginally increase the yield, and reactions at room temperature and at 4°C showed equal amounts of product (Supplement S5 Fig. 1b).

We cleaved the N-terminal 6His-tag from H*-Cdc42:CTIS in two large-scale reactions (using TEV (1) or TEV (3)), and used His-AC to separate cleavage product from uncleaved Cdc42 and protease. To ensure all uncleaved Cdc42 binds to the column, the sample/flow-through got loaded thrice onto the column. The flow-through contained the cleavage product (Fig. 10a). Anti-His Western blots show that a tiny fraction of uncleaved Cdc42 is still present in the flow-through (Fig. 10b). In accordance to the observation of Cdc42 dimers in expression tests before (Fig. 2), we again observed higher bands, correspond to Cdc42 dimers, in the Western blots (Fig. 10b). Both Cdc42 with the N-terminal region and Cdc42 without showed dimers on the blots. These dimers are thus due to properties of Cdc42 and not due to N- or C-terminal additions.

**Figure 10.**
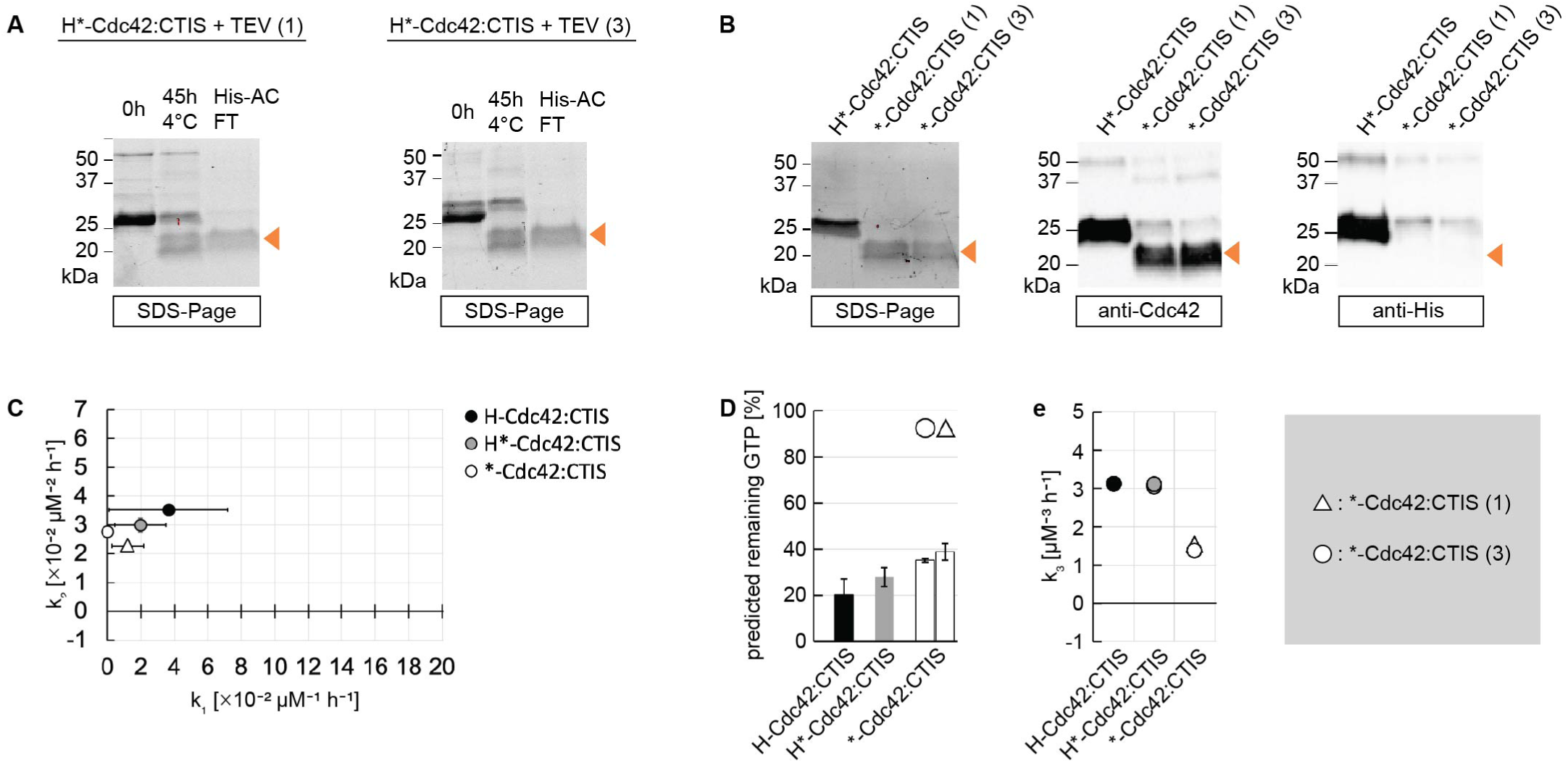
N-terminal tags can be cleaved of Cdc42 with TEV protease: the cleavage product maintains its GTPase activity but shows a reduced interaction with the GEF Cdc24. (A) N-terminal tags can be cleaved off Cdc42 with TEV protease and the cleavage product can be purified using His-AC (FT: flow through). (B) SDS-Page and Western blots of the purified cleavage reaction product of reactions shown in (A). (C) GTP hydrolysis cycling rates (*k*_1_, *k*_2_) of different Cdc42 constructs. (D) Predicted amount of remaining GTP for Cdc42 constructs (5 µM) after an incubation time of 1.5 h, calculated using rates *k*_1_, *k*_2_ shown in (C). Symbols indicate if a specific protein batch was used. (E) Cdc42-Cdc24 interaction rates *k*_3_. (c-E) Summary graphs of the data, values, and overview figures are given in Supplement S4.

We tested the cleavage educts and products on their GTPase activity and ability to interact with the GEF Cdc24. H*-Cdc42:CTIS, containing an N-terminal TEV site, is similar in both of these two properties to H-Cdc42:CTIS, which contains an N-terminal enterokinase site (Fig. 10c-e). Thus, the cleavage site sequence (enterokinase vs. TEV protease cut site) does not influence protein function. The cleavage products of both reactions with TEV protease (*-Cdc42:CTIS) had a slightly reduced GTPase activity (Fig. 10c,d), though the magnitude of this reduction lies within assay variability and is thus likely not due to protein properties (see Supplement S1). It’s interaction with Cdc24, however, was significantly reduced (*k*_3,*-Cdc42:CTIS_ = 1.4−1.5 µM^−3^ℎ^−1^, Fig. 10e). We measured the GEF interaction of H-Cdc42-F in the same assay: *k*_3,H-Cdc42-F_ = 4.5 µM^−3^ℎ^−1^. The low *k*_3_ of *-Cdc42:CTIS is thus not due to assay variation, but describes a reduced or weaker interaction with Cdc24. It is possible that the N-terminal additions boost Cdc42’s ability to interact with Cdc24 or that during the cleavage reaction the protein got damaged.

In conclusion, enterokinase is not a suitable enzyme for cleaving additions/tags off Cdc42. TEV protease might a better option, as it does not lead to undesired cleavage products and works robustly in many conditions. The cleavage product can easily be isolated using His-AC, even though a very small fraction of 6His-tagged Cdc42 remains mixed with the cleavage product after His-AC. However, the cleaved product shows a significantly reduced Cdc24 interaction. This indicates that during both cleavage reactions the protein got damaged or that the N-terminal additions to Cdc42 aid GEF interaction. If the protein got damaged, it questions if TEV protease is indeed a suitable cleavage enzyme for Cdc42.

## Conclusion and Discussion

Here, we shared our experiences of working with *S. cerevisiae* Cdc42 *in vitro*, to make its *in vitro* use more accessible to a broader audience. We discussed our construct design and the expression and purification of Cdc42 and Cdc42-mNeonGreen^*SW*^ /-sfGFP^*SW*^ constructs. We explored two assays for Cdc42 activity checks (GTPase Glo assay (Promega), and Flag-pulldown assay), and investigated tag cleavage with enterokinase and TEV protease.

Most Cdc42 constructs expressed in several conditions and could be purified in a high yield. The exceptions were H-Cdc42 (construct type H, where only a 6His-tag was added N-terminally) and SS-Cdc42-H (type D). H-Cdc42 did not express at all, and we suspect that either a linker region between an N-terminal tag and Cdc42, or the T7 tag in particular, is required for Cdc42 expression in *E. coli*. SS-Cdc42-H expressed well, but precipitated after purification even at low concentrations and showed almost no GTPase activity. As neither S-Cdc42-H, F-Cdc42-H nor H-Cdc42-SS exhibited these issues, we conclude that the N-terminal Twin-Strep-tag is responsible for this. We also observed negative effects of the N-terminal Twin-Strep-tag for Cdc24 and Bem1 constructs, which did not express well or at all. These two protein are larger and less tightly folded than Cdc42, making them more susceptible to degradation and/or folding issues. We think that these early and easily detectable expression issues already indicate a general problem with the construct design: in the case of the bigger and less stable Cdc24 and Bem1 it already showed up on the expression level. In the case of the smaller and more stable Cdc42 it was only observable after purification. We suggest to not use an N-terminal Twin-Strep-tag on (any) protein constructs. Our data shows that even a stable and easy to work with protein (as Cdc42 is) can have a few conditions where it does not behave (express or be stable), creating unpredictable barriers of entry for its *in vitro* use.

These issues were exacerbated for Cdc42-mNeonGreen^*SW*^ and -sfGFP^*SW*^. Here additional constructs (H-Cdc42-mNeon-F, H-Cdc42-mNeon-S) did not express the full-size protein, even though their Cdc42 counterparts did. It seems that purification tags influence the expression and degradation levels for those constructs. Cdc42 is a rather small and tightly folded protein. The sandwich fusions of Cdc42 and sfGFP and mNeonGreen (Bendezú et al., 2015) consist of two folded proteins parts. Because the sequence of sfGFP/mNeonGreen is inserted into that of Cdc42, Cdc42 can only completely fold if the fluorophore is fully folded. As long as both protein parts are not full folded, the entire construct is more susceptible to degradation. It is therefore not surprising to see almost no degradation bands in the Cdc42 expression tests, but many degradation bands in the Cdc42-mNeonGreen^*SW*^ /-sfGFP^*SW*^ expression tests. However, the reason why the two tag combinations, that hinder expression of the full-size protein, exactly have their observed effect remains elusive to us.

After purifying a protein, it ought to be tested for its activity. We used the GTPase Glo assay to investigate the GTPase activity of Cdc42 constructs and their interaction with the GEF Cdc24, and assesd Flag-pulldown assays on their usability to study the interaction of Cdc42 with its other interactors.

The GTPase assay is a suitable assay to test the GTPase activity of Cdc42; it includes all steps in the GTPase cycle and can measure the effect of GEFs and GAPs as well. Our data showed that there was little variation between assays assessing the same Cdc42 purification batch. There can, however, be a large variation between purification batches of the same Cdc42 construct (even though the same purification protocol and conditions were used). This means that: (1) One should test all Cdc42 purification batches on their activity, as purification batches can vary a lot. (2) When comparing different Cdc42 constructs, one should purify several batches of each construct and test each. Otherwise construct differences can be confused with what are in fact purification batch differences. (3) One should be critical/reserved in assuming that kinetic rates and other protein-specific parameters derived from other Cdc42 batches are exactly applicable to one’s protein. The exact numbers of such rates can be both context specific (e.g. depend on buffer components, temperature, …) and purification batch specific.

We also used the GTPase assay to study the Cdc42 - Cdc24 (GEF) interaction. Cdc24 was highly active, and sub-micromolar additions boosted Cdc42 GTPase cycle drastically. We suspect that this leads to the larger variation of Cdc24’s effect between assays; small dilution errors and activity changes of Cdc24 affect the entire GTPase cycle stronger than similar errors and changes in Cdc42. To quantify the Cdc42 - Cdc24 interaction, one should thus conduct the assays a sufficient number of times to get a sense of the ‘normal’ variation. When using this assay to quantitatively compare the GEF interaction of several Cdc42 constructs, one should thus add all Cdc42 constructs to the same assay. If this is not feasible, the addition of a reference Cdc42 construct to all assays is advantageous, as then rate values can be normalised to the rate of the reference construct in each assay.

Considering these assay and purification batch variations, we conclude that most Cdc42 constructs showed a similar GTPase activity and GEF interaction. Surprisingly, all C-terminal additions did not impede Cdc42’s function. So even though so-far no C-terminally tagged Cdc42 was used in *in vitro* studies, these seem to be acceptable additions. On the contrary, Cdc42’s N-terminus seemed to be more delicate: Constructs with an N-terminal Twin-Strep-tag were not active at all and those with an N-terminal 6His-Twin-Strep-tag showed both a reduced GTPase activity and GEF interaction. Additionally, Cdc42 construct type H did not express at all, and Cdc42 where the N-terminal tag got cleaved showed also a reduced GEF interaction. When designing new Cdc42 constructs, we thus advise to be more careful when placing N-terminal additions, and to check literature for evidence that the intended additions are not interfering with Cdc42’s functionality. If such data is not available, we advise to design several variations of this construct and test them in parallel.

In contrast to GTPase assays, we found Flag-pulldown assays to be potentially more difficult for studying protein functionality. They are versatile for studying any protein - protein interaction, as only two differently tagged proteins are needed. In our experience they also gave clear results when working with two strongly interacting proteins (e.g. Cdc24 and Bem1). This must, however, not be true for Cdc42 - effector interactions. We tested the Cdc42 - Bem1 interaction, and could not reproduce previous results (Bose et al., 2001). The Western blot data showed blotting intensities that varied per experiment and would be easy to misinterpret, when not done repeatedly. We think this assay can be a useful tool, but requires several controls and replicas to ensure genuine outcomes.

Lastly, we tested two cleavage enzymes to cleave the N-terminal tag of Cdc42 to produce Cdc42 without additions. Our data showed that enterokinase is not a suitable enzyme, as it cleaves also within Cdc42. TEV protease showed a better cleavage behaviour. The cleavage product, however, showed a reduced GEF interaction. This reduction could have several causes: (1) Due to external circumstances Cdc42 misfolded (as we saw during the purification of one S-Cdc42-H construct) during the cleavage reaction. (2) The reaction with TEV protease lead to the changes in Cdc42, and the TEV enzyme might not be a good choice for cleaving tags off Cdc42. (3) The N-terminal additions on Cdc42 enhance its ability to interact with Cdc24, facilitating the reaction. From our data we can not identify which of these causes is most likely and additional experiments are needed. These include repeating the cleavage with TEV protease, using another cleavage enzyme, and expressing only C-terminally tagged Cdc42. Depending on the cause, this speaks for or against cleaving N-terminal additions off Cdc42: If Cdc42’s N-terminus is sensitive and changes/modifications easily result in protein misfolding and a reduced activity, N-terminal tags ought to be left unchanged, thus avoiding the possibility of damaging the protein. If N-terminal tags/additions on Cdc42 facilitate its GEF interaction, they should be cleaved. This would also mean that all previously conducted *in vitro* studies using N-terminally tagged Cdc42 potentially show an effect facilitated by additions/tags, as other Cdc42 - effector interactions could be equally effected by the tag. Studies comparing and parameterising Cdc42 constructs with N- and/or C-terminal tags would thus be beneficial to the entire community, setting the foundation for all other *in vitro* studies. It would also be interesting to see if only C-terminally tagged Cdc42 expresses and if it is functional. Such a construct would avoid all possible issues of the N-terminus and can be a great choice if post-translational modifications are not needed.

To make working with Cdc42, and any protein in general, more accessible for a broader spectrum of scientists, it will also be hugely beneficial if publications state and explain the used construct designs and their effects on the protein behaviour, as well as show data of constructs and experiments that “failed”.

We here explored several Cdc42 construct designs, including Cdc42 with different N- and C-terminal tags, and Cdc42-fluorophore sandwich fusions. What can they be used for?

1. Establishing robust protocols for *in vitro* prenylation of Cdc42 (Fig. 11a): All our Cdc42 constructs are recombinantly expressed in *E. coli* and lack Cdc42’s C-terminal post-translational prenylation, which enables Cdc42 to bind to membranes. Obtaining prenylated Cdc42 is still a barrier to entry. Only a few groups succeeded, through obtaining the protein using (1) insect cell expression systems (Zheng et al., 1994, 1995; Zhang & Zheng, 1998; Zhang et al., 1999; Kozminski et al., 2003; Johnson et al., 2009, 2012), (2) through purification of membrane-bound Cdc42 from yeast (Das et al., 2012; Rapali et al., 2017), (3) through *in vitro* prenylation of purified unprenylated Cdc42 from *E. coli* (Golding et al., 2019), or (4) through using cell-free expression systems (without purifying the protein) (Sonal et al., 2022). These four methods are powerful tools, but may not be accessible for everyone. Insect cell expression systems require culturing facilities that are not available at every research location, purification from yeast remains not as reproducible as needed, and *in vitro* prenylation of Cdc42 requires additional purification and testing of the prenylation enzyme, whose activity might vary among different purification batches, and cell-free expression systems leave a lot of other proteins in the reaction mixture, requiring similar purification steps as *in vivo* expressions. Our H-Cdc42:CAAX constructs can be used to establish robust and reproducible protocols for *in vitro* prenylation (farnesylation or geranylgeranylation) of recombinantly expressed Cdc42. Further, we prenylated double-tagged Cdc42 (of which the C-terminal tag is a 6His-tag (F-Cdc42-H, S-Cdc42-H)) in a Sortase-mediated reaction (Fig. 11a) (Tschirpke, Spitzbarth, et al., 2024): Sortase A (from now on referred to as ‘Sortase’) is an enzyme that in one reaction cleaves a C-terminal group off a protein and ligates another target group to the protein. Sortase is widely used as it accepts a wide range of target groups (Antos et al., 2017; Popp & Ploegh, 2011) and is commercially available (e.g. Sortase mutant with improved catalytic properties (Chen et al., 2016), available from BPS Bioscience). The cleavage and ligation occur in one incubation step, after which prenylated Cdc42 can be purified in a two-step purification (see Fig. 11a).
2. Fluorescent Cdc42 for microscopy studies (Fig. 11b): We here showed how Cdc42-fluorophore sandwich fusions can be expressed and purified. These fluorescent Cdc42 constructs can be used for any microscopy studies on Cdc42. Sandwich fusions are more ideal than Cdc42 where a fluorophore is added to its N- or C-terminus, as N-terminal fusions can compromise Cdc42’s functions *in vivo* (Bendezú et al., 2015) and Cdc42’s C-terminus is subject to post-translational prenylation. If membrane binding is required, our Cdc42-fluorophore sandwich fusions can be prenylated in a Sortase-mediated reaction as described above (Tschirpke, Spitzbarth, et al., 2024) (Fig. 11a). If fluorescent Cdc42 that does not bind to membranes is needed (e.g. Fig 11c), unprenylated Cdc42-fluorophore sandwich fusions or S-Cdc42-BC-H, fluorescently labelled in a Sortase-mediated reaction, can be used. The use of S-Cdc42-BC-H can be advantageous: (1) We showed that the flexible C-terminal BC sequence does not interferre with Cdc42’s GTPase activity or GEF interaction. It serves as a flexible linker region between Cdc42 and the fluorophore, preventing any possible functionality-impeding effects of the fluorophore on Cdc42. (2) S-Cdc42-BC-H can be fluorescently labelled with almost any fluorescent dye, enabling access to a wide spectral range.
3. Recruitment tests (Fig. 11c): To initiate budding, Cdc42 accumulates in one spot on the cell membrane. The molecular mechanism driving Cdc42 accumulation is still heavily debated (Vendel et al., 2019; Goryachev & Leda, 2017). It was proposed that a double-positive feedback loop drives Cdc42 accumulation: membrane-bound Cdc42 recruits Bem1 to the membrane, and in turn a Bem1-Cdc24 complex recruits more cytosolic Cdc42 to the membrane (Klünder, Freisinger, Wedlich-Söldner, & Frey, 2013). To experimentally test and quantify these feedback loops, total internal reflection fluorescence (TIRF) microscopy and fluorescently labelled TwinStrep-tagged proteins can be used: First a biotinylated membrane coated with streptavidin is made on a microscopy cover slip. Then a TwinStrep-tagged protein is added, which binds with high affinity to the streptavidin coated membrane (Schmidt et al., 2013). Then another fluorescently labelled, not-Strep-tagged protein is added. TIRF microscopy is used to assessed if the membrane-bound protein recruits the other protein to the membrane. To test, for example, the feedback loops proposed in the model by Klünder *et al*. (Klünder et al., 2013), H-Cdc42-sfGFP-SS can be tethered to the membrane and it can be explored if and to which extend it recruits Bem1. If recruitment of Cdc42 ought to be assessed F-Cdc42-sfGFP-H (or any fluorescent Cdc42 that does not contain a Strep- or TwinStrep-tag) can be used.
4. Investigations on Cdc42’s GTPase activity and its regulation (Fig. 11d): Cdc42’s GTPase activity is highly regulated by GEFs and GAPs, and the GTPase cycle involves three steps: GTP binding, GTP hydrolysis, and GDP release. Although the kinetics of the individual reactions steps and the influence of GEFs and GAPs were studied in the 1990s and 2000s (for an overview see Supplement S 1), the reaction cycle of yeast Cdc42 is still not entirely understood. For one, many studies used human (and truncated) Cdc42. Cdc42 is a highly conserved and human Cdc42 shows an 80% sequence identity to yeast Cdc42 (Diepeveen et al., 2018), but both Cdc42 proteins differ in their C-terminal region (PBR and CAAX domain). These C-terminal differences have been shown to influence Cdc42’s GTPase properties (Zhang & Zheng, 1998; Zhang et al., 1999, 2000). Further, for Cdc42-effector studies often only effector domains (instead of full-size proteins) were used and the concentration-dependent profile of an effector, as well as the interplay between effectors, was rarely studied. It has been suggested that the GAPs possess a different level of GAP activity (Smith, Givan, Cullen, & Sprague, 2002), but *in-vitro* studies quantifying the full-size proteins are lacking. Next to the concentration-dependent effect of Cdc42 GTPase regulators, their interplay is relevant as well: Rapali *et al*. showed that Bem1 enhances the GEF-activity of Cdc24 (Rapali et al., 2017) and we recently found that the GEF Cdc24 exhibits synergy with the GAP Rga2 (?, ?). We believe that in order to understand (and model) the GTPase behaviour of Cdc42, more in-depth *in-vitro* studies of its GTPase cycle, its regulation through individual effectors, as well as the interplay of effectors is required. All of our functional Cdc42 constructs (e.g. all H-Cdc24:CAAX constructs, H-Cdc42-F, F-Cdc42-H, …) and the GTPase assay and model described here can be used for such investigations.
5. Investigating the relationship between the PBR sequence and dimerisation and Cdc42’s GAP activity towards itself (Fig. 11e): Many small GTPases contain a C-terminal PBR which has been linked to di-/ oligomerisation (for an overview see Supplement S 2). However, the direct correspondence between the PBR’s amino acid sequence and oligomerisation remains elusive. Human Cdc42 has been shown to dimerise and to exhibit a GAP-activity towards itself (Zhang & Zheng, 1998; Zhang et al., 1999; Zhang, Gao, Moon, Zhang, & Zheng, 2001) (see Supplement S 1), due to an arginine residue in its PBR (Zhang et al., 1999). In the data presented here we observed a small non-linear increase of Cdc42’s GTPase activity with its concentration, suggesting that it could also exhibit a small GAP activity towards itself and potentially form dimers. Further studies, that systematically and thoroughly map out relationship between the amino acid sequence of the PBR and (1) protein dimerisation and (2) if and to which extend a protein shows GAP activity towards itself, are needed to unravel where the non-linearity originates from. It was shown that an arginine present in the PBR leads to protein exhibiting GAP activity towards itself (Zhang et al., 1999), but (1) if the position of this arginine affects its catalytic activity and (2) if other amino acids can lead to a similar/weaker effect (which is possible given that the molecular mechanism of GTP hydrolysis through GTPases is still discussed (Calixto, Moreira, & Kamerlin, 2020)) remains unclear. Our H-Cdc42:CAAX constructs, with systematic changes in their PBR, can be used to answer these questions.
6. Confirming binding partners of Cdc42 (Fig. 11f): Lastly, Flag- and TwinStep-tagged Cdc42 constructs can be used in Flag- or Strep-pulldown assays to confirm direct binding of yeast proteins to Cdc42. Both tags are significantly smaller than the commonly used GST-tag (Bose et al., 2001; Zheng et al., 1995) (Flag-, TwinStrep-tag: <5 kDa, GST-tag: 26 kDa) and introduce less steric hindrance between both proteins. However, we found that pulldown assays require significant controls and optimisation steps in order to reliably detect weakly binding protein-protein interactions. Applications are therefore limited to strongly interacting protein or require significant optimisation of the protocols used. Establishment of such robustly working protocols for protein-protein binding assays will be beneficial to the community, as they will enable more groups to verify protein-protein interactions *in-vitro*.

**Figure 11.**
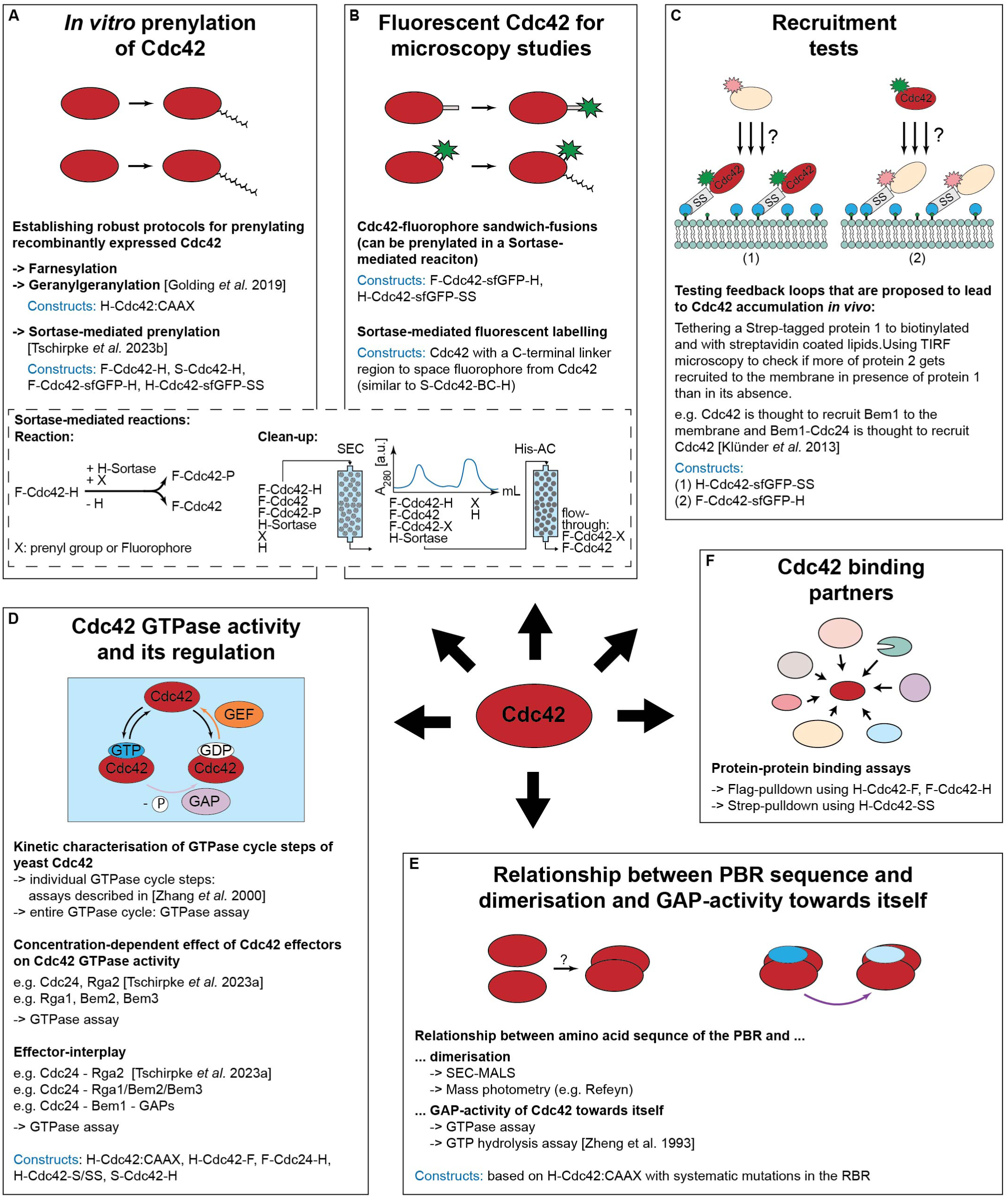
Overview of potential applications of the Cdc42 constructs of this publication. Abbreviations: GAP: GTpase activating protein. PBR: polybasic region. Stated references: Golding *et al*. 2019: (Golding et al., 2019). Klünder *et al*. 2013: (Klünder et al., 2013). Tschirpke *et al*. 2023a: (?, ?) Tschirpke *et al*. 2023b: (Tschirpke, Spitzbarth, et al., 2024). Zhang *et al*. 2000: (Zhang et al., 2000). Zheng *et al*. 1993: (Zheng et al., 1993).

Taken together, we here show how to recombinantly express and purify Cdc42, how to assess its GTPase activity, and outline possible use-cases for our constructs and describe open questions. We aspire that this work sets a basis for more investigations on detailed mechanism of Cdc42 and its regulation, to as a community move towards understanding how Cdc42 establishes polarity.

## Materials and Methods

### Plasmid construction

Genes of interest were obtained from the genome of *Saccharomyces cerevisiae* W303, or in the case of mNeonGreen and sfGFP from plasmids, and were amplified through PCR. The target vector was also amplified through PCR. Additionally, each PCR incorporated a small homologous sequences needed for Gibson assembly (Gibson et al., 2009). After Gibson assembly, the resulting mixture was used to transform chemically competent Dh5α and BL21 DE::3 pLysS cells and plated out onto a Petri dish containing Lysogeny broth agar and the correct antibiotic marker. The primers used for each PCR can be found in Supplement S7 Tab. 1. Gibson assembly resulted in plasmids found Tab. 2 (of which protein sequences are stated in Supplement S6) and Tab. 3.

**Table 3.** List of additional protein constructs/plasmids used throughout this publication.

| Plasmid | Description | Source |
| --- | --- | --- |
| pWKD011a | Cdc42-sfGFP<sup>SW</sup> | this work |
| pWKD017 | Cdc42-mNeonGreen<sup>SW</sup> | this work |
| pET28a-His-mcm10-Sortase-Flag | plasmid template for N-term. 6His- and C-term. Flag-tagged proteins | received from N. Dekker (TU Delft), based on pBP6 (Douglas & Diffley, 2016) |
| pSF1818 | TEV protease, referred to as TEV (3) in this publication | purified protein received from C. Dekker (TU Delft) |

### Buffer composition

If not mentioned otherwise, the buffers are of the composition stated in Tab. 4.

**Table 4.** Buffer composition.

| Buffer | Composition |
| --- | --- |
| Lysis buffer | 50 mM Tris-HCl (pH=8.0), 1 M NaCl, 5 mM imidazole, 1 mM 2-mercaptoethanol (Sigma Aldrich), supplemented with EDTA-free Protease inhibitor cocktail (Roche) and 1 mM freshly prepared PMSF. |
| His-AC washing buffer | 50 mM Tris-HCl (pH=8.0), 1 M NaCl, 5 mM imidazole, 1 mM 2-mercaptoethanol (Sigma Aldrich). |
| His-AC elution buffer | 50 mM Tris-HCl (pH=8.0), 100 mM NaCl, 500 mM imidazole, 1 mM 2-mercaptoethanol (Sigma Aldrich). |
| Strep-AC washing buffer | 50 mM Tris-HCl (pH=8.0), 100 mM NaCl, 1 mM 2-mercaptoethanol (Sigma Aldrich). |
| Strep-AC elution buffer | 50 mM Tris-HCl (pH=8.0), 100 mM NaCl, 10 mM desthiobiotin (Sigma Aldrich), 1 mM 2-mercaptoethanol (Sigma Aldrich). |
| SEC buffer | 50 mM Tris-HCl (pH=7.5), 100 mM NaCl, 10 mM MgCl<sub>2</sub>, 1 mM 2-mercaptoethanol (Sigma Aldrich). |

### Protein expression tests

Proteins were expressed in Bl21::DE3 pLysS cells, which carry the gene for the bacteriophage T7 RNA polymerase under the regulation of a Lactose dependent promoter. Expression of the T7 RNA polymerase, which for example can be induced by IPTG, results in the transcription and therefore expression of the genes of interest that are placed under the T7 promoter. This system is of advantage, as the T7 RNA polymerase transcribes 5-10× faster than E. coli RNA polymerase (F. Studier & Moffatt, 1986; Dubendorf & Studier, 1991). Four conditions were used:

1. Cells were grown in Lysogeny broth at 37°C until an OD_600_ of 0.7, the expression was induced through addition of 1.0 mM IPTG, after which cells were grown for 3 h at 37°C.
2. Cells were grown in Lysogeny broth at 37°C until an OD_600_ of 0.7, the expression was induced through addition of 0.2 mM IPTG, after which cells were grown for 18 h at 18°C.
3. Cells were grown in Studier Induction ZYP-5052 Medium for 3 h at 37°C, followed by 18 h at 18°C, and in accordance to the recommended protocol (F. W. Studier, 2005).

Cells were harvested through centrifugation. Cell pellets were resuspended in SDS loading buffer (Laemmli buffer, (Laemmli, 1970) and expression levels were analysed through SDS-Page and Western blotting.

### Protein purification

The expression and purification protocols for Cdc24, Cdc42, and Bem1 described here are based, with modifications, on the Cdc24 purification protocol described previously (Rapali et al., 2017).

The methods used are described in the following and protein-specific purification routes are stated thereafter.

#### Protein expression

The proteins were expressed in Bl21::DE3 pLysS cells. Cells were grown in Lysogeny broth at 37°C until an OD_600_ of 0.7. One of these two induction conditions was chosen:

1. Cells were grown in Lysogeny broth at 37°C until an OD_600_ of 0.7, the expression was induced through addition of 1.0 mM IPTG, after which cells were grown for 3 h at 37°C.
2. Cells were grown in Lysogeny broth at 37°C until an OD_600_ of 0.7, the expression was induced through addition of 0.2 mM IPTG, after which cells were grown for 18 h at 18°C.

Cells were harvested through centrifugation.

#### Lysis and His affinity chromatography (His-AC)

Cell pellets were resuspended in lysis buffer and lysed with a high pressure homogenizer (French press cell disruptor, CF1 series Constant Systems) at 4°C, using 5-10 rounds of exposing the sample to pressurisation. The cell lysate was centrifuged at 37000× g for 30 min and the supernatant was loaded onto a HisTrap^TM^ excel column (Cytiva)^1^. After several rounds of washing with His-AC washing buffer, the protein was eluted in a gradient of His-AC washing buffer and His-AC elution buffer.

#### Strep affinity chromatography (Strep-AC)

For some proteins Strep-AC was chosen for further purification. The His-AC elution peak was loaded onto a StrepTrap^TM^ HP column (Cytiva). After several rounds of washing with Strep-AC washing buffer, the protein was eluted in a gradient of Strep-AC washing buffer and Strep-AC elution buffer.

#### Size exclusion chromatography (SEC)

Some proteins were further purified by SEC using SEC buffer and a HiPrep 16/60 Sephacryl S-300 HR (Cytiva) column.

#### Dialysis and storage

All proteins were dialysed twice in SEC buffer. After the addition of 10% glycerol, samples were flash frozen in liquid nitrogen and kept at -80°C for storage.

#### Expression and purification conditions per protein

H-Cdc42-F, F-Cdc42-H, H-SS-Cdc42-F, S-Cdc42-H, S-Cdc42-BC-H, H-Cdc42-SS, H-Cdc42:CTIS, H*-Cdc42:CTIS, and F-Cdc42-sfGFP-H were expressed using expression condition (1) and purified in a single step His-AC using an AKTA Pure (Cytiva) machine. The purification profile of H-SS-Cdc42-F, as an example for Cdc42 purification, is shown in Fig. 3a and that of F-Cdc42-sfGFP-H in Fig. 5.

H-Cdc42-sfGFP-SS was expressed using expression condition (1) and purified using His-AC followed by Strep-AC. Here lysis buffer, His-AC washing buffer, and His-AC elution buffer were supplemented with 0.1 v/v% Tween-20 (Sigma Aldrich), 0.1 v/v% NP-40 (Thermo Fischer Scientific), and 0.1 v/v% Triton-X-100 (Sigma Aldrich). The purifications were done using an AKTA Pure (Cytiva) machine. Purification profiles are shown in Supplement S3 Fig. 1.

F-Cdc42-mNeon-H was expressed using expression condition (1) and purified using His-AC followed by SEC using an AKTA Pure (Cytiva) machine. Purification profiles are shown in Supplement S3 Fig. 2.

S-Cdc42-mNeon-BC-H was expressed using expression condition (1) and purified using His-AC followed by SEC. Here lysis buffer, His-AC washing buffer, and SEC buffer were supplemented with 0.1 v/v% Tween-20 (Sigma Aldrich), 0.1 v/v% NP-40 (Thermo Fischer Scientific), and 0.1 v/v% Triton-X-100 (Sigma Aldrich). The purifications were done using an AKTA Pure (Cytiva) machine. Purification profiles are shown in Supplement S3 Fig. 3.

Cdc24-H was expressed using expression condition (2) and purified using His-AC followed by SEC using an AKTA Pure (Cytiva) machine.

H-Bem1-F was expressed using expression condition (2) and purified in a single-step His-AC using an AKTA Pure (Cytiva) machine. Fig. 3b and 12 shows all purified proteins on SDS-Page.

**Figure 12.**
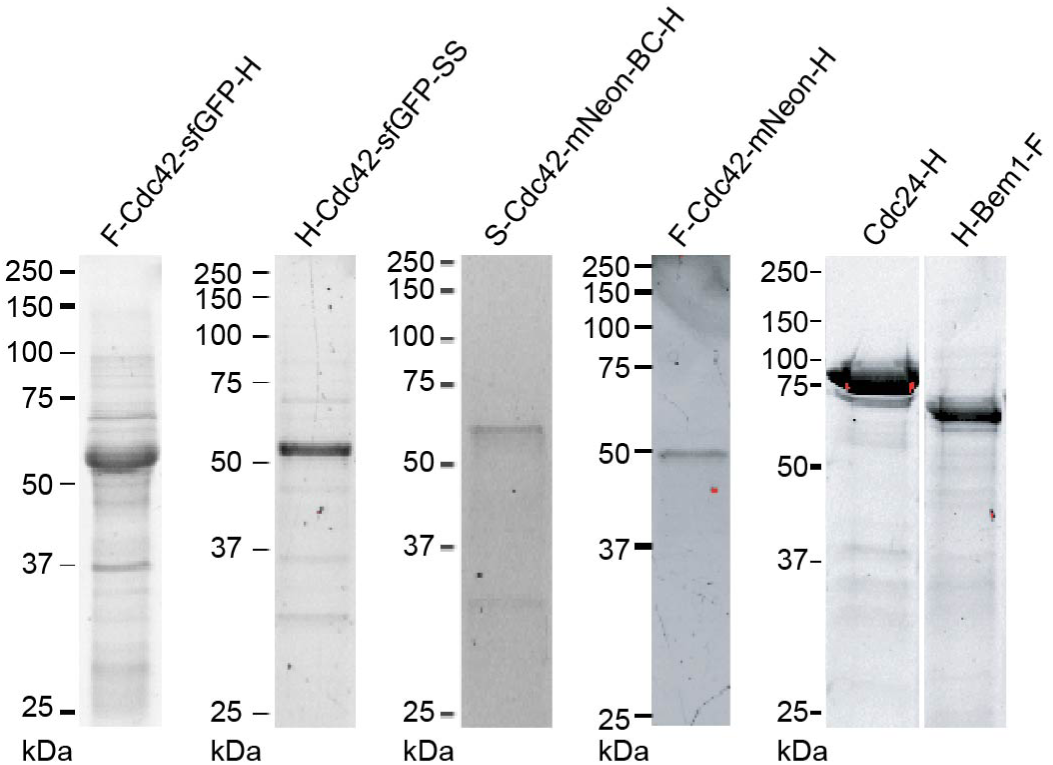
SDS-Page of purified Cdc24, Bem1, and fluorescent Cdc42. Purified Cdc42 is shown in Fig. 3b.

### GTPase activity assay

GTPase activity was measured using the GTPase-Glo^TM^ assay (Promega) as described previously (Mondal, Hsiao, & Goueli, 2015). In brief, 5 µL protein in SEC buffer (Tab. 4) was mixed with 5 µL of a GTP-solution (10 µM GTP, 50 mM Tris-HCl (pH=7.5), 100 mM NaCl, 10 mM MgCl_2_, 1 mM 2-mercaptoethanol (Sigma Aldrich), 1 mM dithiothreitol (VWR) in 384-well plates (Corning) to initiate the reaction. The reaction mixture got incubated at 30°C on an Innova 2300 platform shaker (New Brunswick Scientific) (120 rpm), before the addition of 10 µL Glo buffer and another 30 min incubation. The Glo buffer contains a nucleoside-diphosphate kinase that convert remaining GTP to ATP. Addition of 20 µL detection reagent, containing a luciferase/luciferin mixture, makes the ATP luminescent, which was read on a Synergy HTX plate reader (BioTek) in luminescence mode. The amount of hydrolysed GTP inversely correlates with the measured luminescence. Wells without protein (’buffer’) were used for the normalization and represent 0% GTP hydrolysis (Eq. 3). Reactions were carried out with at least 4 replicates (wells) per assay, and the average (’Lum.’) and standard deviation (’ΔLum.’) of each set was used to calculate the activity and error of each set.

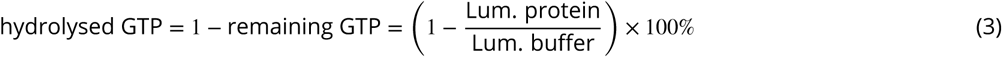

Error bars were calculated by using error propagation:

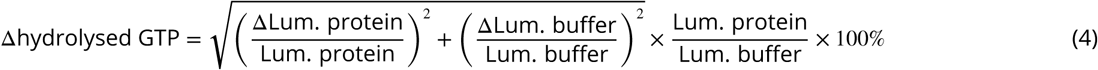

The amount of remaining GTP at the time of reaction termination (*t*_*term*_.) was calculated by

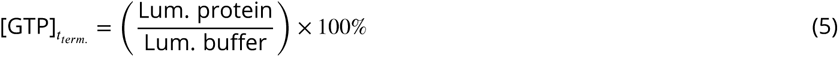

We developed a Cdc42 GTPase activity model for determining the GTPase cycling rates *k*. It is described in the following:

The GTP hydrolysis process involves three steps: (1) A GTP molecule from solution binds to Cdc42. (2) Cdc42 hydrolyses GTP. (3) Cdc42 releases GDP.

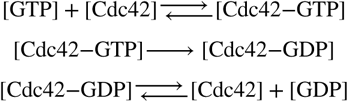

It can further be upregulated by effector proteins: GAPs have been shown to enhance GTP hydrolysis by Cdc42 (step 2), GEFs enhance the release of GDP from Cdc42 (step 3) (Martin, 2015; Chiou, Balasubramanian, & Lew, 2017) and the scaffold Bem1 enhances the GEF activity of Cdc24 (Rapali et al., 2017).

To quantitatively describe the GTPase reaction cycle, we coarse-grained the GTPase reaction steps with

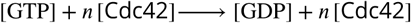

To account for possible Cdc42 dimerisation and cooperativity, we included the following reactions into the model: (1) We assume that Cdc42 can dimerise, as other small GTPases have been shown to dimerise (Zhang & Zheng, 1998; Zhang et al., 1999, 2001; Kang et al., 2010):

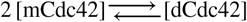

and both monomeric and dimeric Cdc42 can contribute to the overall GTP hydrolysis with different rates:

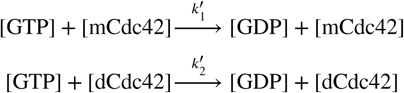

Assuming that the majority of Cdc42 is in its monomeric form ([mCdc42] < *C*_*d*_, with *C*_*d*_ as the concentration at which half of the total Cdc42 is dimeric), we can approximate

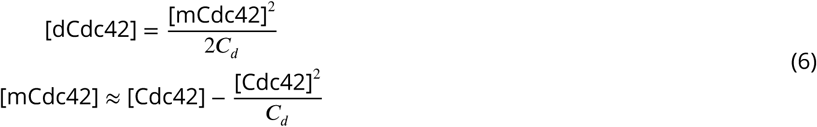

(2) Next to cooperativity from dimerisation, cooperativity can also emerge when Cdc42 proteins come in close contact with each other - they can affect each other’s behaviour without forming a stable homodimer, effectively functioning as an effector protein for themselves:

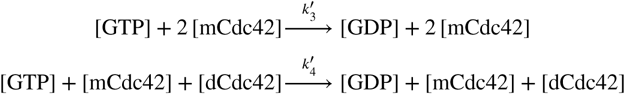

(3) Effector proteins, such as GAPs and GEFs, affect the speed of the GTP hydrolysis cycle:

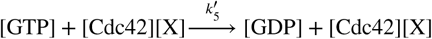

Here X is an effector protein.

Our data showed that the amount of remaining GTP follows an exponential decline over time:

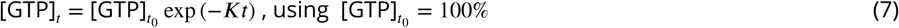

Considering reactions (1) - (3), we can thus define *K* in Eq. 7 as

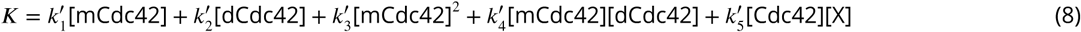

Using Eq. 6, and considering only up to second-order terms, results in

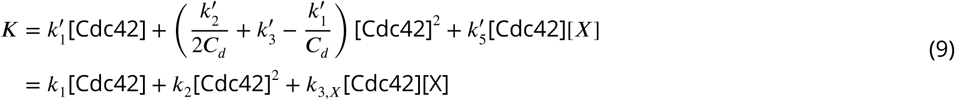

where *k*_1_ refers to GTP hydrolysis cycling rates of monomeric Cdc42, *k*_2_ includes effects of cooperativity and dimerisation and *k*_3_ represents the rate of Cdc42 - effector interaction. We will refer to *K* as ‘overall GTP hydrolysis rate’ in the following.

We used Eq. 9 with [X]=0 to determine the rates of Cdc42 alone. We then conducted assays with Cdc42 and an effector protein to determine *k*_3_. While doing so we needed to account for assay variability, i.e. for the observation that the rates for Cdc42 can vary between assays. Possible reasons for this include small concentration differences introduced though pipetting of small volumes (as are required for this assay), temperature and shaker speed fluctuations during the incubation step, and/or intrinsic changes in the protein activities due to other external conditions. To account for this variance, we introduced the parameter *c*_*corr*_. It maps all factors that lead to variations between assays onto the Cdc42 concentration.

The assay data, including samples containing only Cdc42 and Cdc42 - (effector) protein mixtures, was fitted with

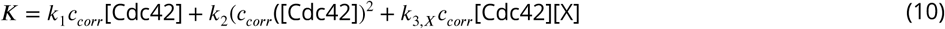

to determine *c*_*corr*_ and *k*_3,*X*_ (using *k*_1_ and *k*_2_ determined earlier).

Our data suggested that the GTP decline in assays Cdc42-Cdc24 mixtures does not depend linearly on the Cdc24 concentration (Eq. 10), but is better approximated by a quadratic term:

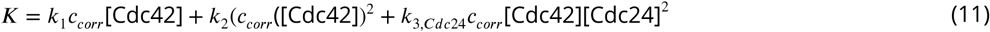

The GTPase assays conducted here involve many pipetting steps and can, in done without a lot of attention, be prone to errors. These result in assays with lots of variability and big error bars. Most GTPase assays resulted in data with small error bars. We used the following criteria to exclude assays from our analysis:

- The error of buffer wells (ΔLum. buffer) is above 10%. This generally means that the assay and all assay data has big error bars.
- The variability of the Cdc42 activity between assays using only Cdc42 and Cdc42 - Cdc24 mixtures is high, i.e. *c*_*corr*_ < 0.5 or *c*_*corr*_ > 1.6. Assays of only Cdc42 dilutions showed a low variability between assays. In Cdc42 - Cdc24 assays only one Cdc42 concentration was used. If it doesn’t show the same GTPase activity as in the assays with only Cdc42, *c*_*corr*_ deviates from one. This then affects the other fitting parameters and leads to more variability in *k*_3_. Ideally only data of *c*_*corr*_ close to one would be included. As we did not intent to provide a detailed parametrisation of the constructs used, but to use this assay to compare them and to point ot points of failure when working with Cdc42, we included a wider range of data. Most assays, nonetheless, had values of *c*_*corr*_ close to one (see Supplement S4).

Additionally, data points that showed less than 5% remaining GTP were excluded from the fit, as they correspond to almost no measurable luminescence.

A detailed experimental protocol and description of the fitting procedure used is given in (Tschirpke, Daalman, & Laan, 2024), with the software available at (Tschirpke, Daalman, & Laan, 2023).

### Flag pulldown assay

In this assay, Flag-tagged Bem1 (H-Bem1-F) and another protein, that does not contain a Flag-tag, was mixed. For each experiment, 0.2 nmol Bem1, and 1.0 nmol Cdc42 or 0.6-1.4 nmol Ovalbumin (as a neg. control, Gel Filtration HMW Calibration Kit (Cytiva)) were used. If Cdc42 was used, 1.0 nmol got first pre-loaded with either no nucleotide, 100 nmol GDP (Jena Bioscience), or 100 nmol GTPγS (Jena Bioscience) for 30 min at room temperature. Two proteins were incubated for 1 h at 30 °C. 100 µL anti-Flag^®^ M2 Affinity gel (Sigma Aldrich) was added and incubated for another 30 min at room temperature. The reaction mixture was added onto a Poly-Prep Chromatography column (Biorad) and was washed 3× with 1 mL of assay buffer (25 mM Tris-HCl (pH=7.2), 300 mM NaCl, 10% Glycerol, 0.01 v/v% NP-40 (Thermo Fischer Scientific); supplemented either with no nucleotide, 10-100 µM GDP, or 10-100 µM GTPγS). Proteins were eluted with 200 µL assay buffer supplemented with 0.6 mg∕mL 3× Flag^®^ peptide (Sigma Aldrich), and analysed by Western blotting.

### Reactions with enterokinase and TEV protease

For enterokinase screening reactions, 50 µg H-Cdc42:CTIS was mixed with 5, 10, or 20 U of enterokinase (16 U/µL) (P8070, New England Biolabs) in SEC buffer, supplemented with 2 mM CaCl_2_. After 1-3 h at room temperature or 24-48 h at 4°C reaction samples were taken and analysed by SDS-Page.

For TEV protease screening reactions, tree enzymes were assessed: TEV (1): 50 µg H*-Cdc42:CTIS was mixed with 5 or 50 U of TEV protease (10 U/µL) (T4455, Sigma-Aldrich) in SEC buffer. TEV (2): 50 µg H*-Cdc42:CTIS was mixed with 5 or 50 U of AcTEV^TM^ protease (10 U/µL) (Invitrogen) in SEC buffer. TEV (3): 50 µL H*-Cdc42:CTIS (63 µM) was mixed with 1, 5, or 10 µL of purified TEV protease (125 µM, pSF1818) in SEC buffer. The enzyme was received as a gift from C. Dekker lab (TU Delft). After 1-3 h at room temperature or 24-48 h at 4°C reaction samples were taken and analysed by SDS-Page.

For cleavage reactions, H*-Cdc42:CTIS (63 µM) was mixed with either TEV (1) (TEV protease, 10 U/µL, T4455, Sigma-Aldrich) in a 10 U/100 µg TEV/Cdc42 ratio, or with TEV (3) (125 µM, pSF1818) in a 5 µL/100 µL TEV/Cdc42 ratio, in SEC buffer. After an incubation of 45 h at 4°C, the reaction was stopped. The sample was loaded thrice onto a HisTrap^TM^ excel column (Cytiva). The flow-through contains Cdc42 of which the N-terminal 6His-tag got cleaved. Samples were analysed by SDS-Page and Western blotting.

### SDS-Page

SDS-Page gels (12-15% acrylamide) were prepared freshly. In brief, a solution of 375 mM Tris-HCl (pH=8.8), 30-37.5 v/v% 40% acrylamide solution (Biorad), 0.2 w/v% sodium dodecyl sulfate (Sigma Aldrich), 0.5 v/v% 2,2,2-Trichloroethanol (Sigma Aldrich), 0.1 w/v% ammonium persulfate (Sigma Aldrich), and 0.1 v/v% N,N,N’,N’-tetramethyl ethylenediamine (Sigma Aldrich) was prepared and casted into 1.00 mm mini-protean glass plates (Biorad), filling them up to 80%. To protect the gel surface from drying, a layer of isopropanol (Sigma Aldrich) was added. The gel was let solidify for 20 min, after which the isopropanol layer was removed. A solution of 155 mM Tris-HCl (pH=6.5), 10 v/v% 40% acrylamide solution (Biorad), 0.2 w/v% sodium dodecyl sulfate (Sigma Aldrich), 0.1 w/v% ammonium persulfate (Sigma Aldrich), and 0.1 v/v% N,N,N’,N’-tetramethyl ethylenediamine (Sigma Aldrich) was prepared and added to the existing gel layer, after which a well comb (Biorad) was added. The gel was let solidify for 20 min.

Cell and protein samples were mixed with SDS loading buffer (Laemmli buffer, (Laemmli, 1970)). Before loading onto the SDSPage gels, they were kept for 5 min at 95°C. Gels were run for 5 min at 130 V followed by 55min min at 180 V (PowerPac Basic Power Supply (Biorad)). Imaging was done on a ChemiDoc MP (Biorad) using the ‘Stain-free gels’ feature and automatic exposure time determination. Precision Plus Protein Unstained standard (Biorad) was used as a protein standard.

### Western blotting

After SDS-Page, the sample was transferred from the SDS-Page to a blotting membrane (Trans-Blot Turbo Transfer Pack, Bio-rad) using the ‘Mixed MW’ program of the Trans-Blot Turbo Transfer System (Bio-rad). The blotting membrane was incubated with Immobilon signal enhancer (Millipore) at 4°C for 18 h. The blotting membrane was incubated with primary antibody, diluted in Immobilon signal enhancer (see Tab. 5), at room temperature for 1 h. It was washed thrice with TBS-T (10 mM Tris-HCl (pH=7.5), 150 mM NaCl, 0.1 v/v% Tween-20 (Sigma Aldrich)). For each washing step the blotting membrane was incubated with TBS-T at room temperature for 20 min. The blotting membrane was incubated with secondary antibody, diluted in Immobilon signal enhancer (see Tab. 5), at room temperature for 1 h, after which it was again washed thrice with TBS-T. SuperSignal West Pico Mouse IgG Detection Kit (Thermo Scientific) was used for activation. Imaging was done on a ChemiDoc MP (Biorad) using the ‘Chemi Sensitive’ feature and automatic exposure time determination. Precision Plus Protein Unstained standard (Biorad) and Precision Plus Protein All Blue Standard (Biorad) were used as protein standards.

**Table 5.** Antibodies used for Western blotting.

| Blot | Primary antibody | Secondary antibody |
| --- | --- | --- |
| anti-His | His Tag Antibody, Mouse (OAEA00010, Aviva Systems Biology) (dilution: 1:4000) | IgG2b Antibody HRP conjugated (Goat Anti-Mouse) (OASA06620, Aviva Systems Biology) (dilution: 1:1000) |
| anti-Cdc42 | Anti-CDC42 antibody, Rabbit (ab64533, abcam) (dilution: 1:2000) | Polyclonal Goat Anti-Rabbit Immunoglobulins/HRP (Goat Anti-Rabbit) (P0448, Agilent Dako) (dilution: 1:500) |

## Abbreviations

AA: amino acid
GAP: GTPase activating protein
GDI: guanine nucleotide dissociation inhibitor
GEF: GDP/GTP exchange factor
His-AC: His-affinity chromatography
PBR: polybasic region
POI: protein of interest
SEC: size exclusion chromatography
SEC-MALS: size exclusion chromatography - multi-angle light scattering
Strep-AC: Strep-affinity chromatography
SW: sandwich fusion

## Data availability

The GTPase-Glo assay data underlying this publication is available at doi.org/10.4121/ddc72065-1082-4183-908d-a55b04d05cb2 (CC BY-SA 4.0) (Tschirpke, Opstal, Valk, Daalman, & Laan, 2023).

## Contributions

**S. Tschirpke:** Conceptualization, Methodology, Validation, Investigation, Writing - Original Draft, Writing - Review & Editing, Visualization, Project administration. **F. van Opstal:** Investigation, Validation. **R. van der Valk:** Investigation, Validation. **W. K.-G. Daalman:** Conceptualization, Investigation, Software, Formal analysis, Validation, Methodology. **L. Laan:** Writing - Review & Editing, Project administration, Funding acquisition, Supervision.

## Acknowledgments

We thank C. de Agrela Pinto for conducting SEC-MALS with our samples and analysing the data thereof. We thank D. McCusker (University of Bordeaux) for the plasmid pDM272, N. Dekker (TU Delft) for the plasmid pET28a-His-mcm10-Sortase-Flag, and C. Dekker lab (TU Delft) for a sample of purified TEV protease (pSF1818).

L. Laan gratefully acknowledges funding from the European Research Council under the European Union’s Horizon 2020 research and innovation programme (grant agreement 758132) and funding from the Netherlands Organization for Scientific Research (Nederlandse Organisatie voor Wetenschappelijk Onderzoek) through a Vidi grant (016.Vidi.171.060).

## Supporting information captions

**S3 Appendix.** Purification of Cdc42-sfGFP^*SW*^ and -mNeonGreen^*SW*^.

**S6 Appendix.** Amino acid sequences of proteins.

## Supplement S1

**S1 Figure 1.**
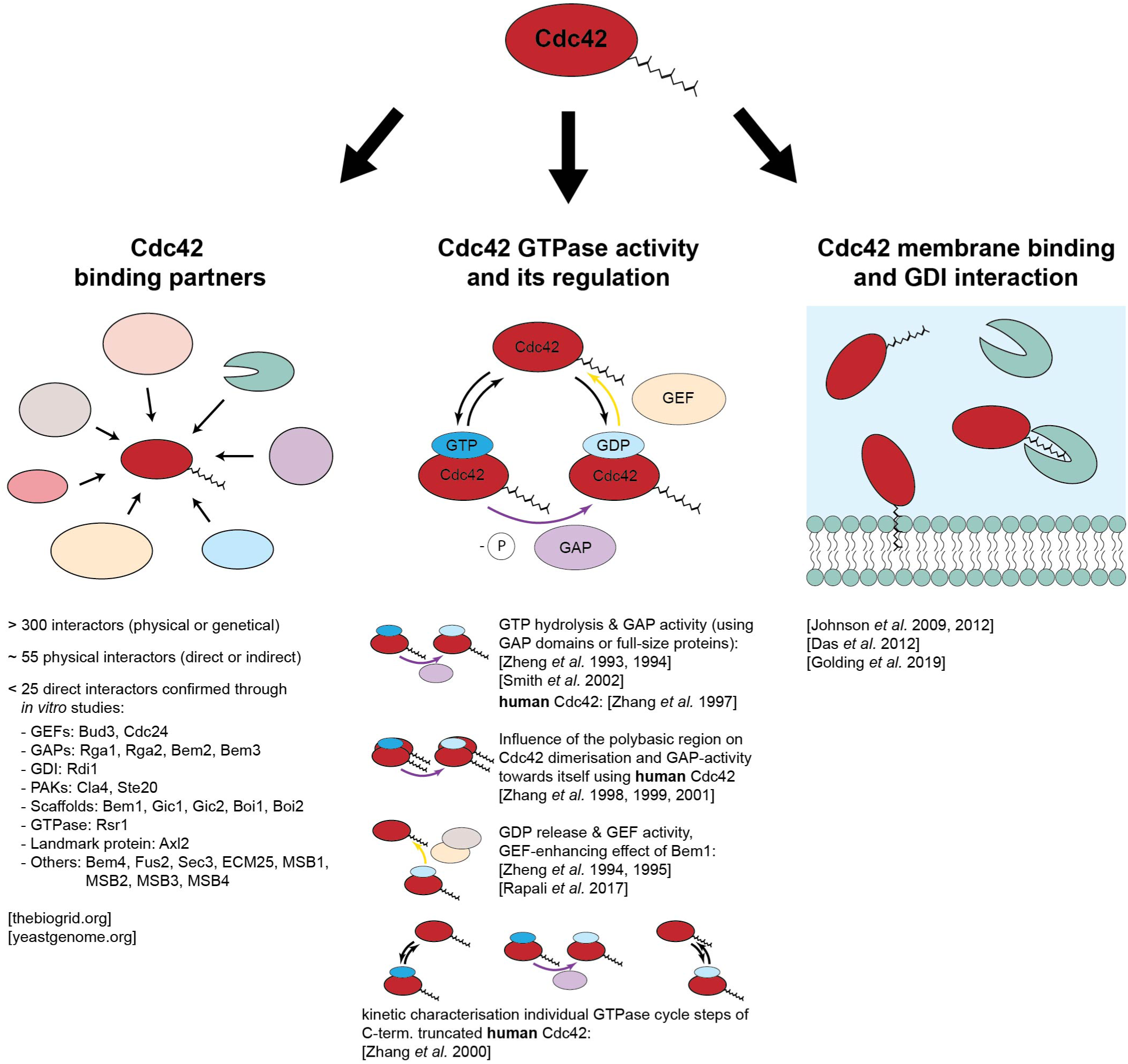
Overview of *in vitro* studies investigating Cdc42’s properties. Abbreviations: GAP: GTPase activating protein. GDI: guanine nucleotide dissociation inhibitor. GEF: GDP/GTP exchange factor. PAK: p21 activated kinase. Stated references: Das *et al*. 2012: (Das et al., 2012). Golding *et al*. 2019: (Golding et al., 2019). Johnson *et al*. 2009: (Johnson et al., 2009). Johnson *et al*. 2012: (Johnson et al., 2012). Rapali *et al*. 2017: (Rapali et al., 2017). Smith *et al*. 2002: (Smith et al., 2002). Zhang *et al*. 1997: (Zhang, Wang, & Zheng, 1997). Zhang *et al*. 1998: (Zhang & Zheng, 1998). Zhang *et al*. 1999: (Zhang et al., 1999). Zhang *et al*. 2000: (Zhang et al., 2000). Zhang *et al*. 2001: (Zhang et al., 2001). Zheng *et al*. 1993: (Zheng et al., 1993). Zheng *et al*. 1994: (Zheng et al., 1994). Zheng *et al*. 1995: (Zheng et al., 1995).

## Supplement S2

**S2 Table 1.**
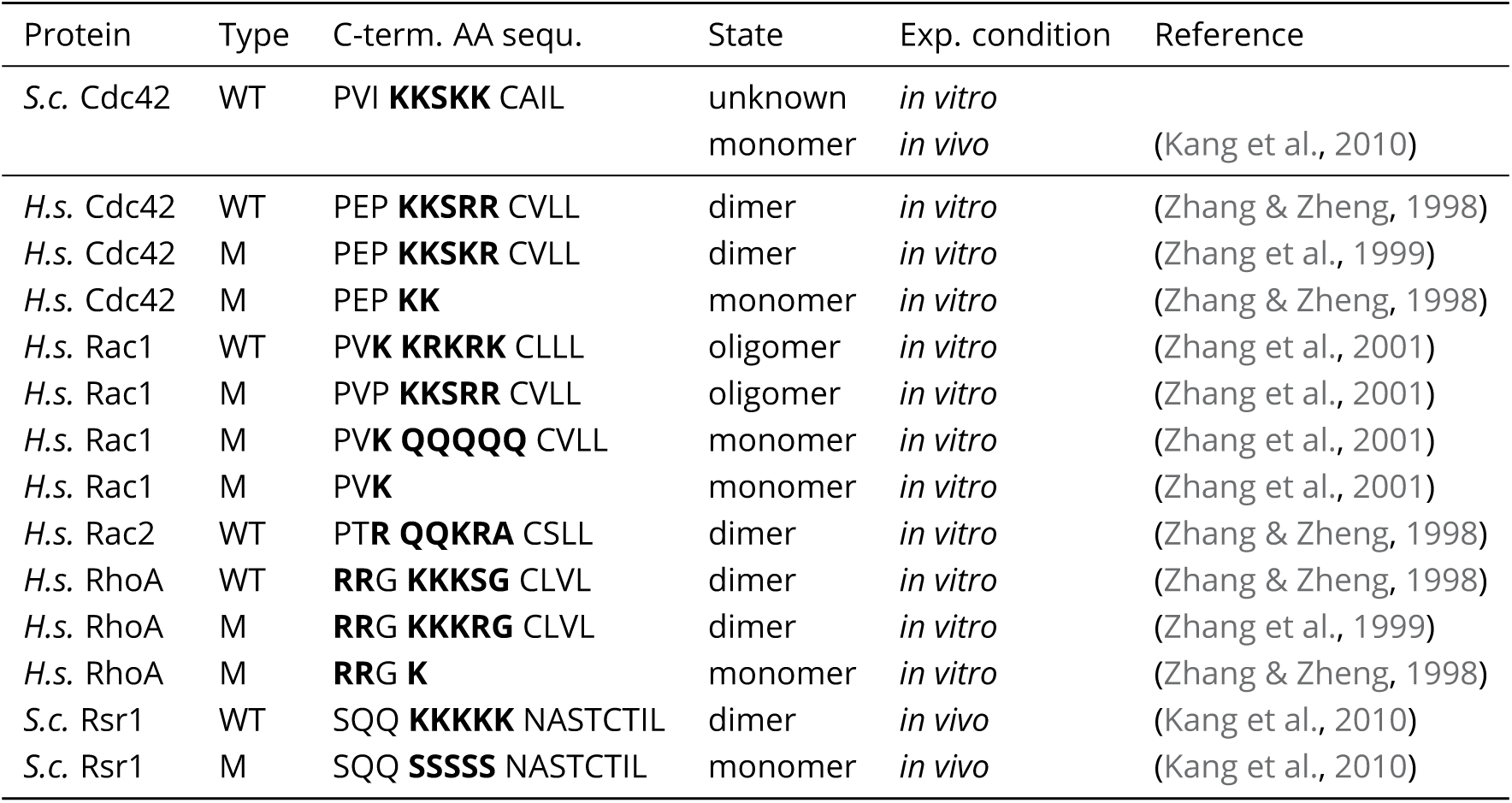
Relationship between polybasic regions (PBRs) and protein dimerisation. Amino acid (AA) sequences of the C-terminal region of selected small GTPases in relation to the ability of these GTPases to dimerise/oligomerise. The PBR and other basic AA (K: lysine, R: arginine) are highlighted in bold. Abbreviations: *S.c.*: *S. cerevisia*, *H.s.*: *H. sapiens*, WT: wild-type, M: mutant.

## Supplement S3

We purified F-Cdc42-sfGFP-H, H-Cdc42-sfGFP-SS, F-Cdc42-mNeon-H, and S-Cdc42-mNeon-BC-H. F-Cdc42-sfGFP-H could be purified in a high yield (30 mg per 6 L expression) in a single step His-AC (Fig. 5), and the other constructs required further purification and gave a significantly lower yield. An overview of the expression and purification conditions used is given in S3 Tab. 1.

H-Cdc42-sfGFP-SS was purified using His-AC followed by Strep-AC (S3 Fig. 1). The final product was pure, but the yield was low. We observed that the addition of detergents doubled the yield (S3 Tab. 1).

F-Cdc42-mNeon-H was purified using His-AC followed by SEC-AC (S3 Fig. 2). After His-AC F-Cdc42-mNeon-H was not pure, in addition to the 50 kDa protein band a ∼70 kDa impurity is visible on SDS-Page (S3 Fig. 2c). The protein was further cleaned by SEC. Here the 50 kDa protein started to elute before a 70 kDa impurity. The peak overlapped strongly with the 70 kDa impurity. Much of the protein fractions therefore had to be discarded, resulting in a low yield.

For S-Cdc42-mNeon-BC-H a similar route was chosen (S3 Fig. 3). After His-AC the elution peak fraction showed also a impurity at ∼70 kDa. In SEC the 70 kDa impurity eluted before the 50 kDa S-Cdc42-mNeon-BC-H. We suspect that this time the order changed to the expected one because detergents were added to the buffer. Because the amount of protein in the His-AC elution peak was little, this purification also resulted in a small yield.

Based on this data we suggest to use detergents in the buffers, as this increases the yield during affinity chromatography and could help in SEC. However, the addition of detergents does not lead to the yield obtained for F-Cdc42-sfGFP-H.

In general, if a protein ought to be purified that does not express in a high yield and requires SEC, we advise to significantly scale-up the expression volume and and use large purification columns: We used up to 12 L medium per expression, and purified it in a single His-AC using a 5 mL column (HisTrap^TM^ excel column (Cytiva)). The bigger column volume allows for faster flow rates and reduces the overall purification time, even when large sample volumes are used. Elution using a steep gradient (or even step elution) concentrates all protein into a small volume, which can immediately be used for SEC. If ideal SEC conditions are unknown, a smaller SEC column (e.g. Superdex 200 Increase 10/300 GL (Cytiva), maximum sample volume of 500 µL) can be used to screen conditions. Smaller columns accommodate less sample volume, but have shorter run-times. Once a good condition is found, we advise to switch to a larger SEC column (e.g. HiPrep 16/60 Sephacryl S-300 HR (Cytiva), maximum sample volume of 5 mL), so that the entire, or at least half of the, His-AC elution peak can be loaded onto the SEC column without any intermediate concentration steps. If the sample after SEC is low in concentration, or samples from several SEC ought to be pooled and concentrated, they can be loaded onto a His-AC column again and eluted in a step elution. This strategy avoids intermediate concentration steps, which, in our experience, reduce the yield up to 30% due to protein sticking to the spin-concentator.

**S3 Table 1.** Overview of expression conditions, volumes, purification steps, and final yields of protein for fluorescent Cdc42 constructs. ’X’ indicates that the respective purification method was used, (det.) indicates that the buffers were supplemented with a detergent mixture (0.1% Tween-20, 0.1% NP-40, 0.1% Triton-X-100). ‘f’ refers to a strong and fast expression at elevated temperatures, induced by a high amount of IPTG (3 h at 37°C with 1 mM IPTG).

| Construct | Expression condition | Expression volume | His-AC | Strep-AC | SEC | Final yield |
| --- | --- | --- | --- | --- | --- | --- |
| F-Cdc42-sfGFP-H | f | 6 L | X |  |  | ~30 mg |
| H-Cdc42-sfGFP-SS | f | 12 L | X | X |  | ~0.5 mg |
| H-Cdc42-sfGFP-SS | f | 12 L | X (det.) | X (det.) |  | ~1 mg |
| F-Cdc42-mNeon-H | f | 6 L | X |  | X | ~0.5 mg |
| S-Cdc42-mNeon-BC-H | f | 6 L | X (det.) |  | X (det.) | ~0.5 mg |

**S3 Figure 1.**
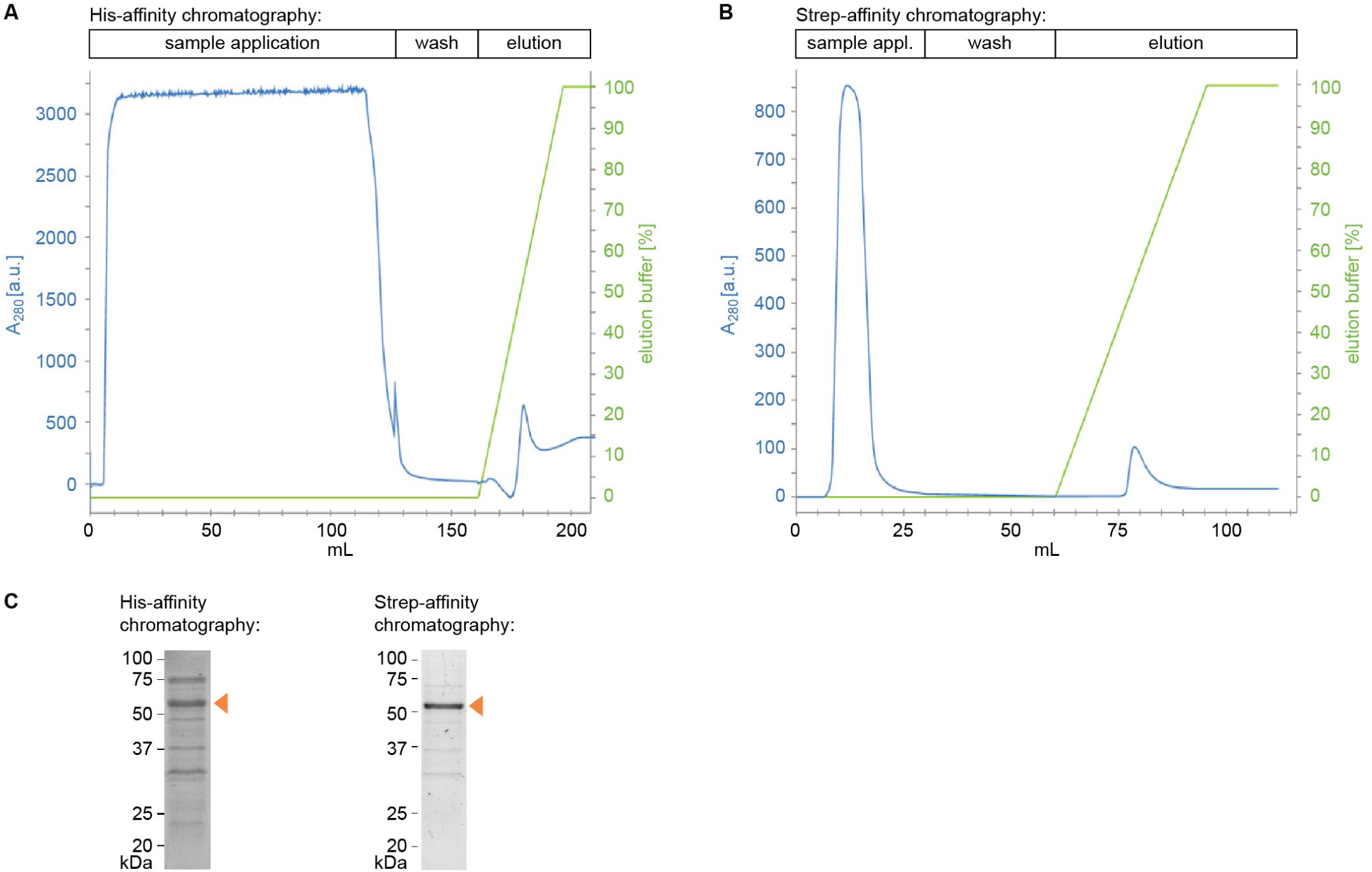
Purification profile of H-Cdc42-sfGFP-SS. H-Cdc42-sfGFP-SS was purified by His-affinity chromatography (A) followed by Strep-affinity chromatography (B). (C) SDS-Page of the elution peaks, the band of the full-length protein is indicated by an orange arrow.

**S3 Figure 2.**
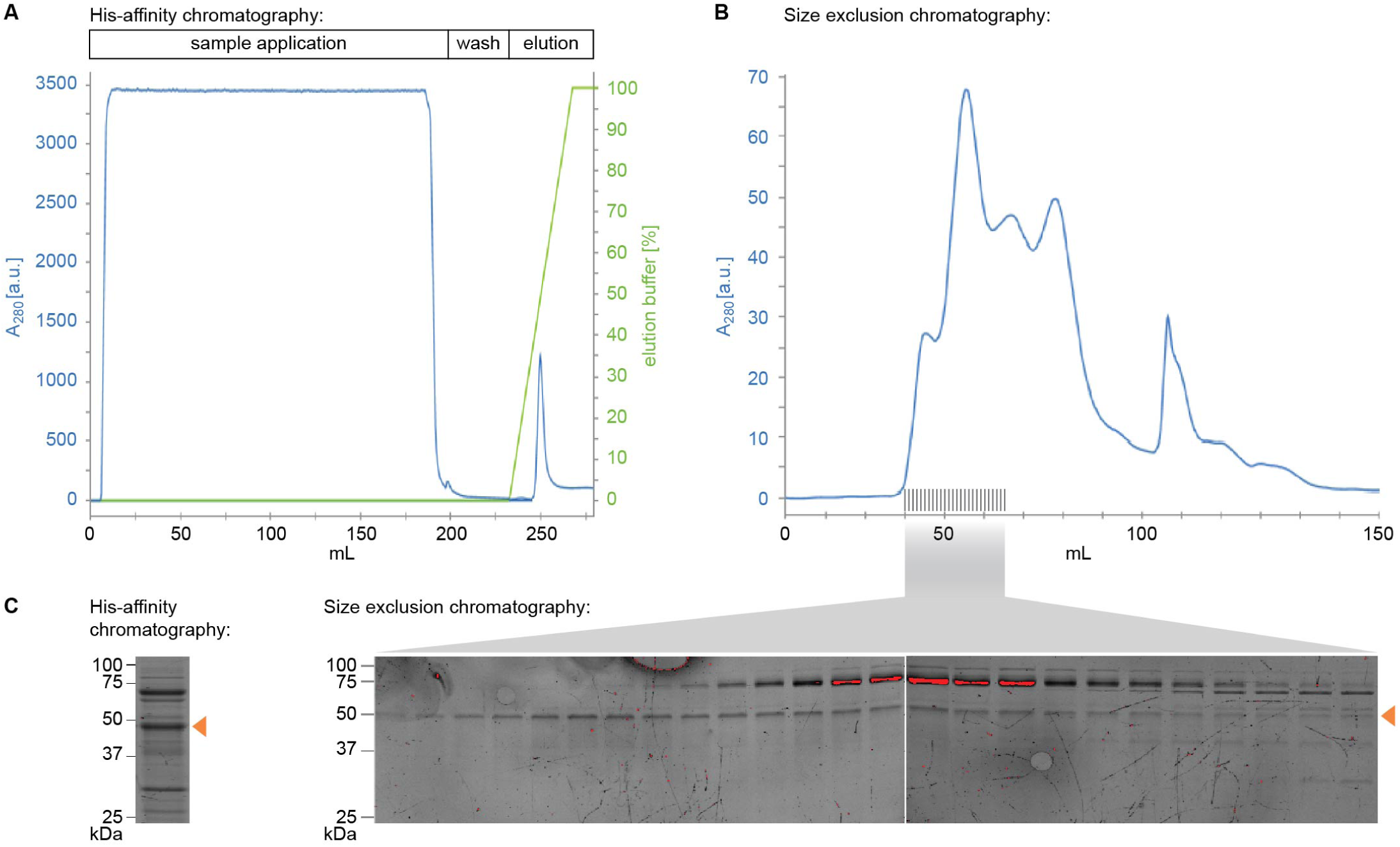
Purification profile of F-Cdc42-mNeon-H. F-Cdc42-mNeon-H was purified by His-affinity chromatography (A) followed by size exclusion chromatography (B). (C) SDS-Page of the elution peaks, the band of the full-length protein is indicated by an orange arrow.

**S3 Figure 3.**
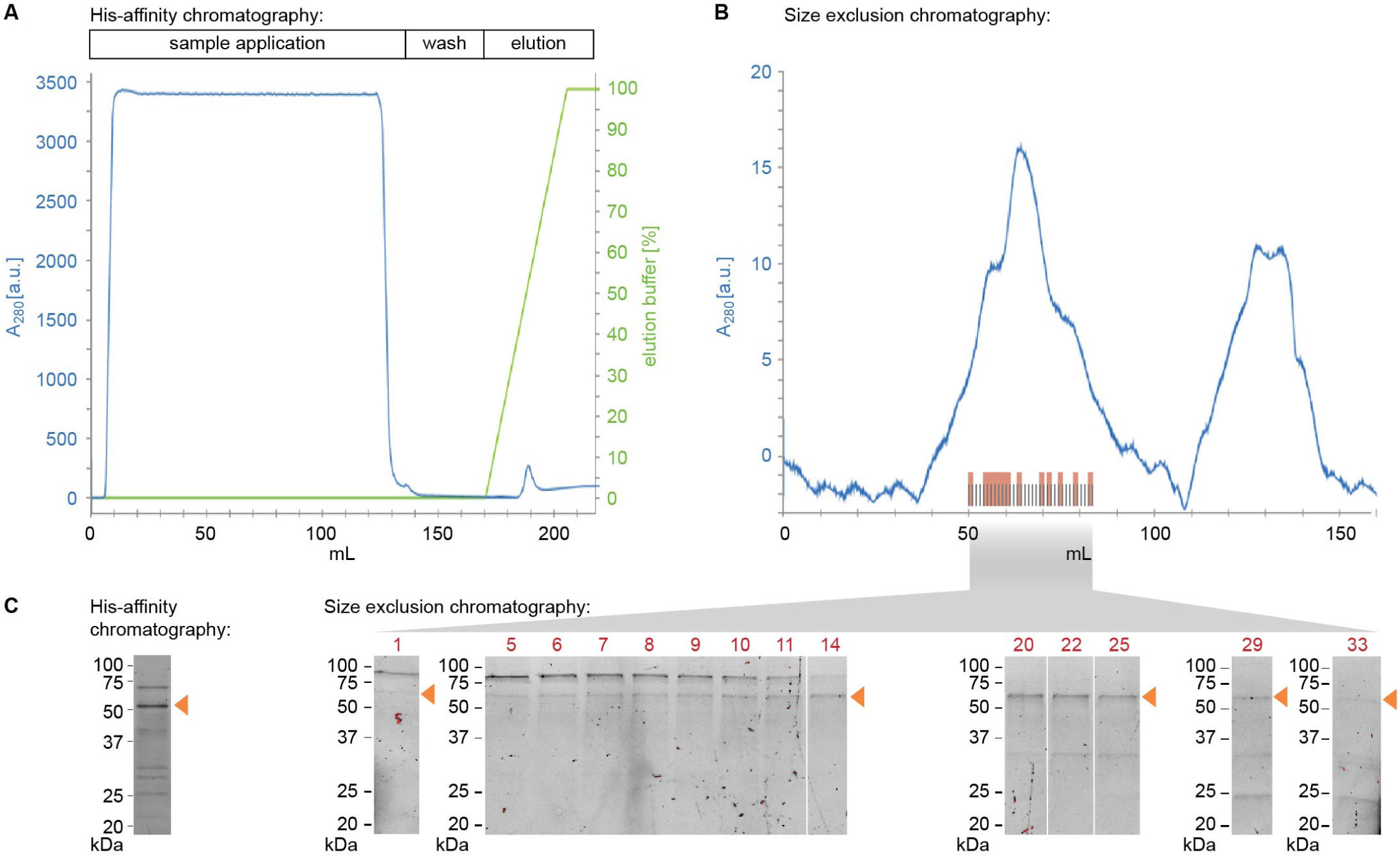
Purification profile of S-Cdc42-mNeon-BC-H. S-Cdc42-mNeon-BC-H was purified by His-affinity chromatography (A) followed by size exclusion chromatography (B). (C) SDS-Page of the elution peaks. The band of the full-length protein is indicated by an orange arrow.

## Supplement S4

**S4 Figure 1.**
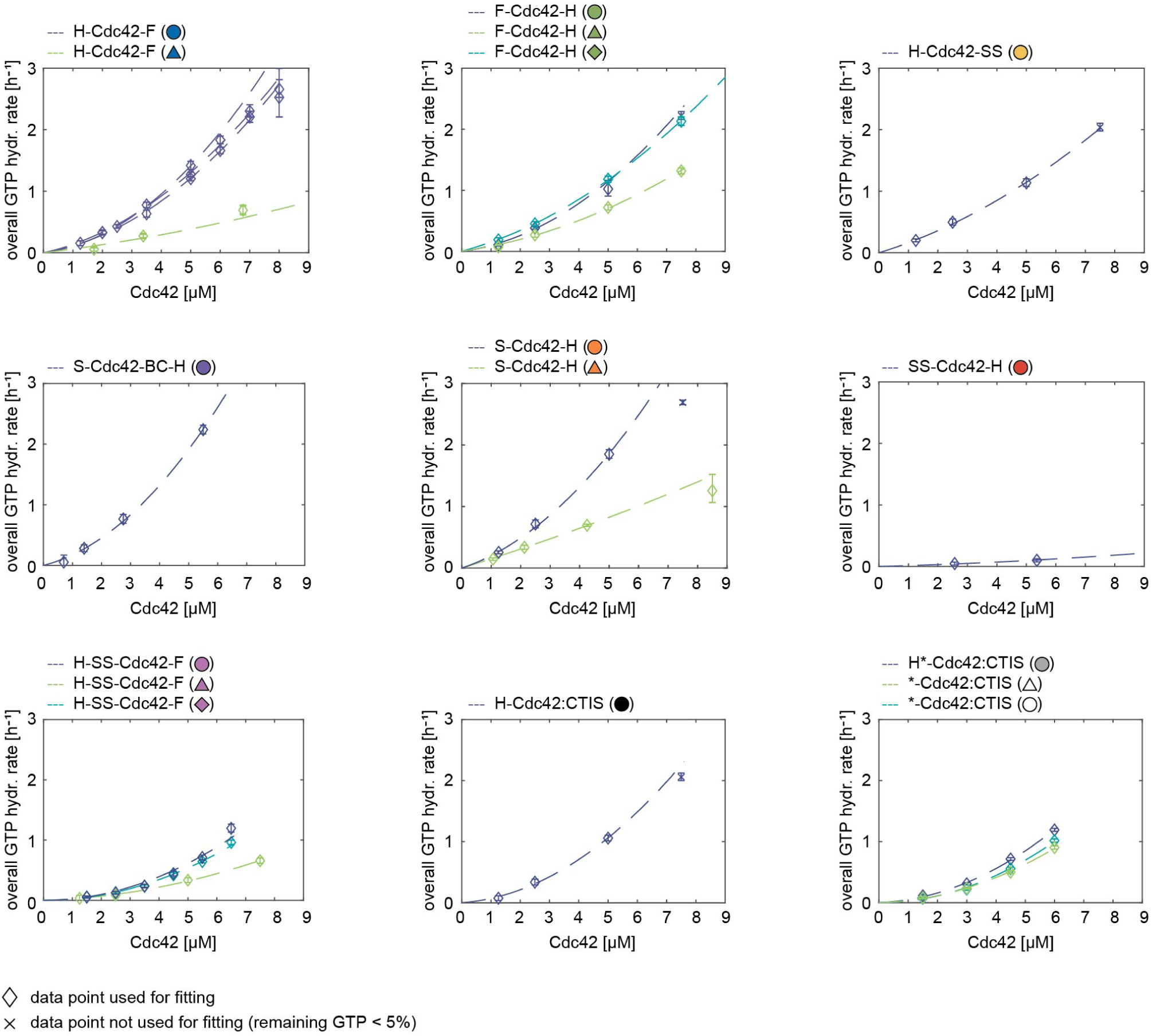
GTPase activity data and fits for Cdc42 constructs.

**S4 Figure 2.**
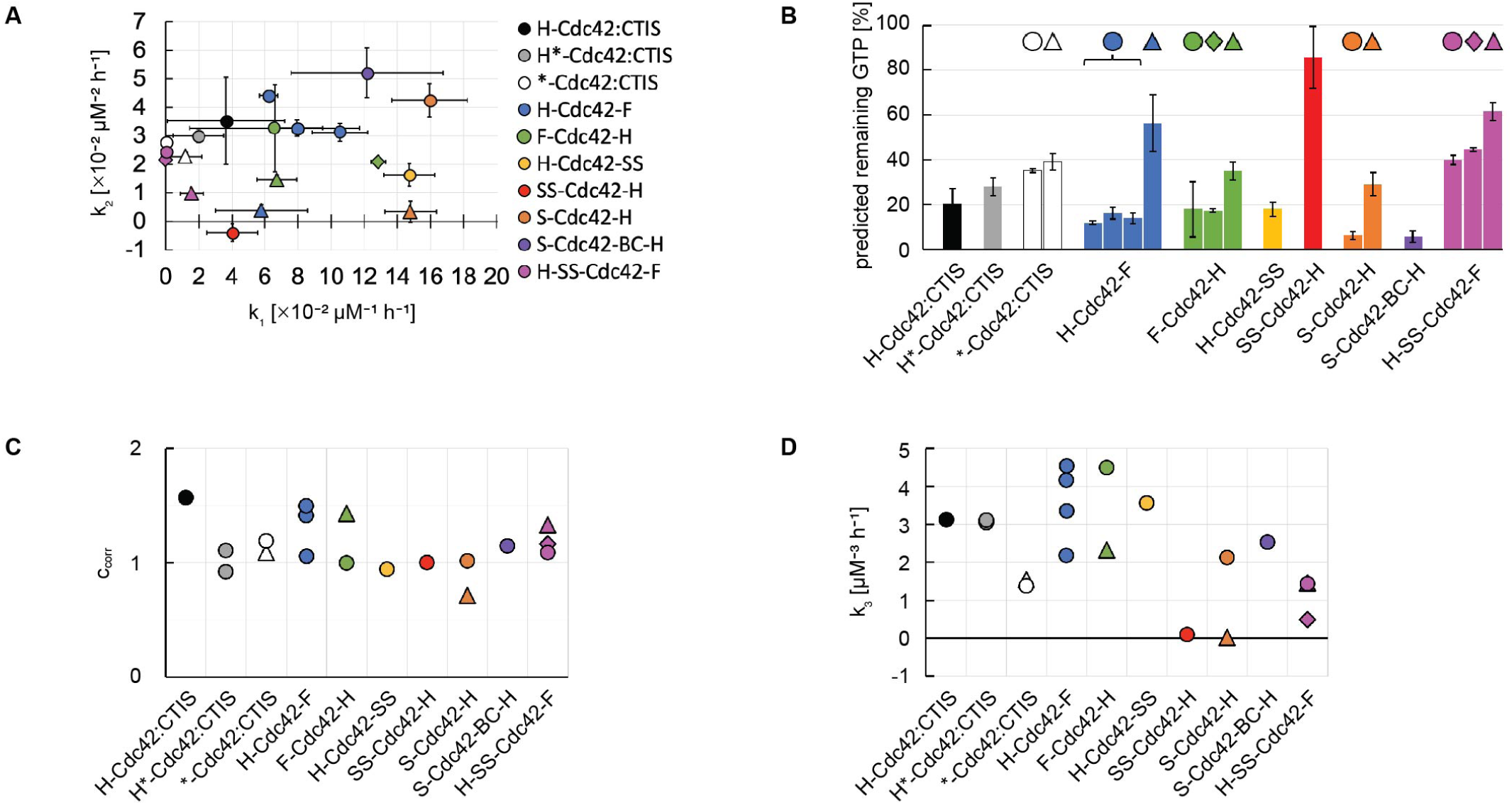
Summary graphs of Cdc42’s GTPase activity and interaction with Cdc24. (A) GTP hydrolysis cycling rates *k*_1_ and *k*_2_. (B) Predicted amount of remaining GTP for Cdc42 constructs (5 µM) after an incubation time of 1.5 h, calculated using rates *k*_1_, *k*_2_ shown in (A). (C) correction factors *c*_*corr*_. (D) Cdc42-Cdc24 interaction rate *k*_3_. (A-D): Data points of the same colour refer to the rates obtained from separate measurements/assays. Data points of the same colour but a distinct shape (circle, triangle, rhombus) represent different purification batches of the same construct.

**S4 Table 1.**
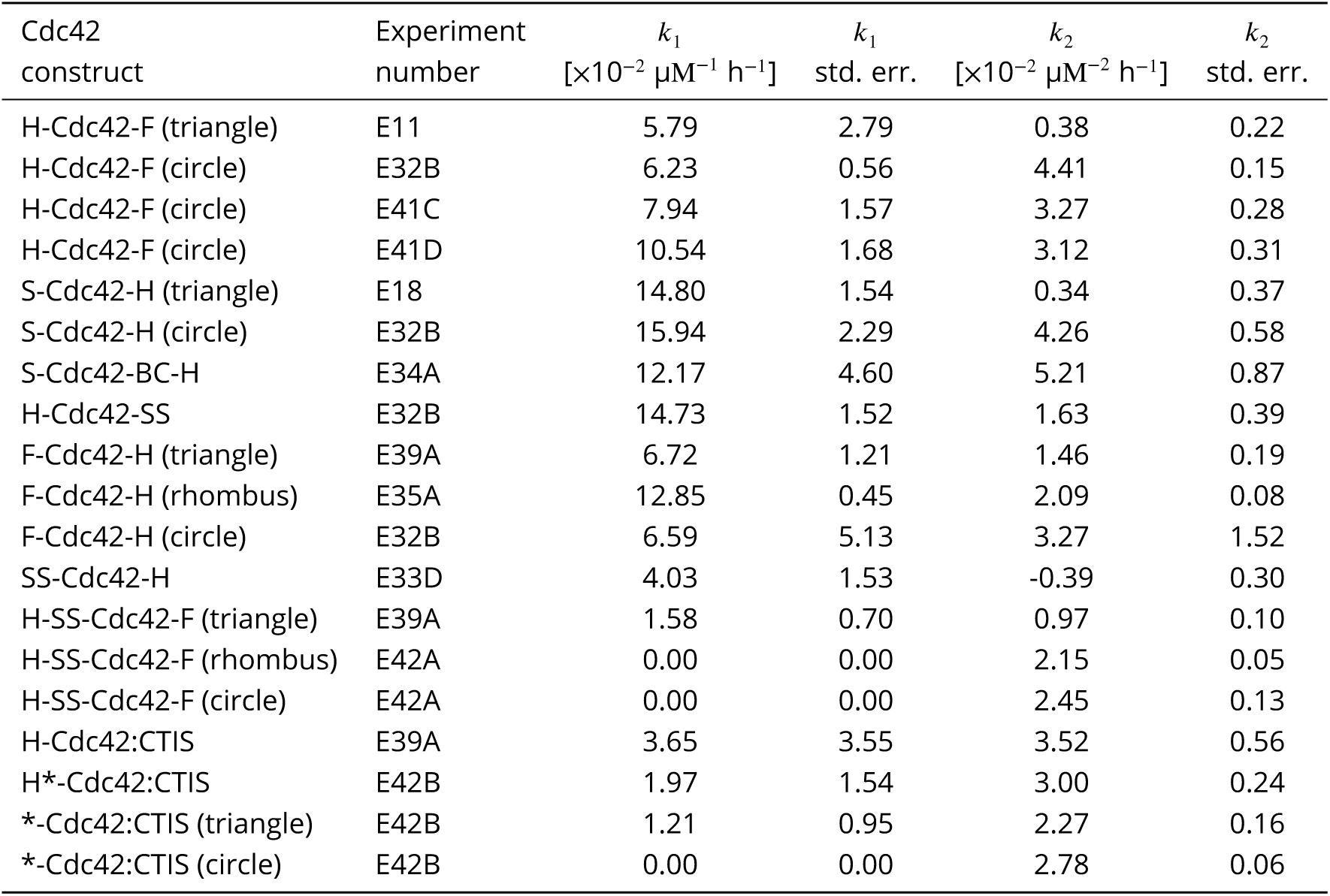
GTP hydrolysis cycling rates *k*_1_ and *k*_2_ of Cdc42. Purification batches are indicated in brackets.

| Cdc42<br>construct | Experiment<br>number | $k_1$<br>[$\times 10^{-2} \mu\text{M}^{-1} \text{h}^{-1}$] | $k_1$<br>std. err. | $k_2$<br>[$\times 10^{-2} \mu\text{M}^{-2} \text{h}^{-1}$] | $k_2$<br>std. err. |
| --- | --- | --- | --- | --- | --- |
| H-Cdc42-F (triangle) | E11 | 5.79 | 2.79 | 0.38 | 0.22 |
| H-Cdc42-F (circle) | E32B | 6.23 | 0.56 | 4.41 | 0.15 |
| H-Cdc42-F (circle) | E41C | 7.94 | 1.57 | 3.27 | 0.28 |
| H-Cdc42-F (circle) | E41D | 10.54 | 1.68 | 3.12 | 0.31 |
| S-Cdc42-H (triangle) | E18 | 14.80 | 1.54 | 0.34 | 0.37 |
| S-Cdc42-H (circle) | E32B | 15.94 | 2.29 | 4.26 | 0.58 |
| S-Cdc42-BC-H | E34A | 12.17 | 4.60 | 5.21 | 0.87 |
| H-Cdc42-SS | E32B | 14.73 | 1.52 | 1.63 | 0.39 |
| F-Cdc42-H (triangle) | E39A | 6.72 | 1.21 | 1.46 | 0.19 |
| F-Cdc42-H (rhombus) | E35A | 12.85 | 0.45 | 2.09 | 0.08 |
| F-Cdc42-H (circle) | E32B | 6.59 | 5.13 | 3.27 | 1.52 |
| SS-Cdc42-H | E33D | 4.03 | 1.53 | -0.39 | 0.30 |
| H-SS-Cdc42-F (triangle) | E39A | 1.58 | 0.70 | 0.97 | 0.10 |
| H-SS-Cdc42-F (rhombus) | E42A | 0.00 | 0.00 | 2.15 | 0.05 |
| H-SS-Cdc42-F (circle) | E42A | 0.00 | 0.00 | 2.45 | 0.13 |
| H-Cdc42:CTIS | E39A | 3.65 | 3.55 | 3.52 | 0.56 |
| H*-Cdc42:CTIS | E42B | 1.97 | 1.54 | 3.00 | 0.24 |
| *-Cdc42:CTIS (triangle) | E42B | 1.21 | 0.95 | 2.27 | 0.16 |
| *-Cdc42:CTIS (circle) | E42B | 0.00 | 0.00 | 2.78 | 0.06 |

**S4 Table 2.** Correction factors *c*_*corr*_ and Cdc42-Cdc24 interaction rate *k*_3_. Purification batches are indicated in brackets.

| Cdc42<br>construct | Experiment<br>number | $c_{corr}$ | $k_3$<br>[$\mu\text{M}^{-3} \text{h}^{-1}$] | $k_3$<br>std. err. |
| --- | --- | --- | --- | --- |
| SS-Cdc42-H | E33D | 1.00 | 0.10 | 0.05 |
| H-Cdc42-F (circle) | E39B | 1.41 | 2.18 |  |
| H-Cdc42-F (circle) | E33C | 1.49 | 4.16 |  |
| H-Cdc42-F (circle) | E33F | 1.46 | 3.36 | 0.68 |
| H-Cdc42-F (circle) | E42D | 1.06 | 4.54 | 0.00 |
| S-Cdc42-H (triangle) | E21 | 0.71 | 0.01 |  |
| S-Cdc42-H (circle) | E33C | 1.01 | 2.13 |  |
| S-Cdc42-BC-H | E34B | 1.15 | 2.53 | 0.39 |
| H-Cdc42-SS | E33C | 0.94 | 3.56 |  |
| F-Cdc42-H (circle) | E33C | 1.00 | 4.49 |  |
| F-Cdc42-H (triangle) | E39B | 1.43 | 2.32 |  |
| H-SS-Cdc42-F (triangle) | E39B | 1.33 | 1.46 |  |
| H-SS-Cdc42-F (rhombus) | E42D | 1.16 | 0.49 |  |
| H-SS-Cdc42-F (circle) | E42D | 1.09 | 1.43 |  |
| H-Cdc42:CTIS | E39B | 1.57 | 3.13 |  |
| H*-Cdc42:CTIS | E42C | 0.92 | 3.04 | 0.89 |
| H*-Cdc42:CTIS | E42D | 1.10 | 3.11 |  |
| *-Cdc42:CTIS (triangle) | E42D | 1.09 | 1.54 |  |
| *-Cdc42:CTIS (circle) | E42D | 1.19 | 1.38 |  |

## Supplement S5

**S5 Figure 1.**
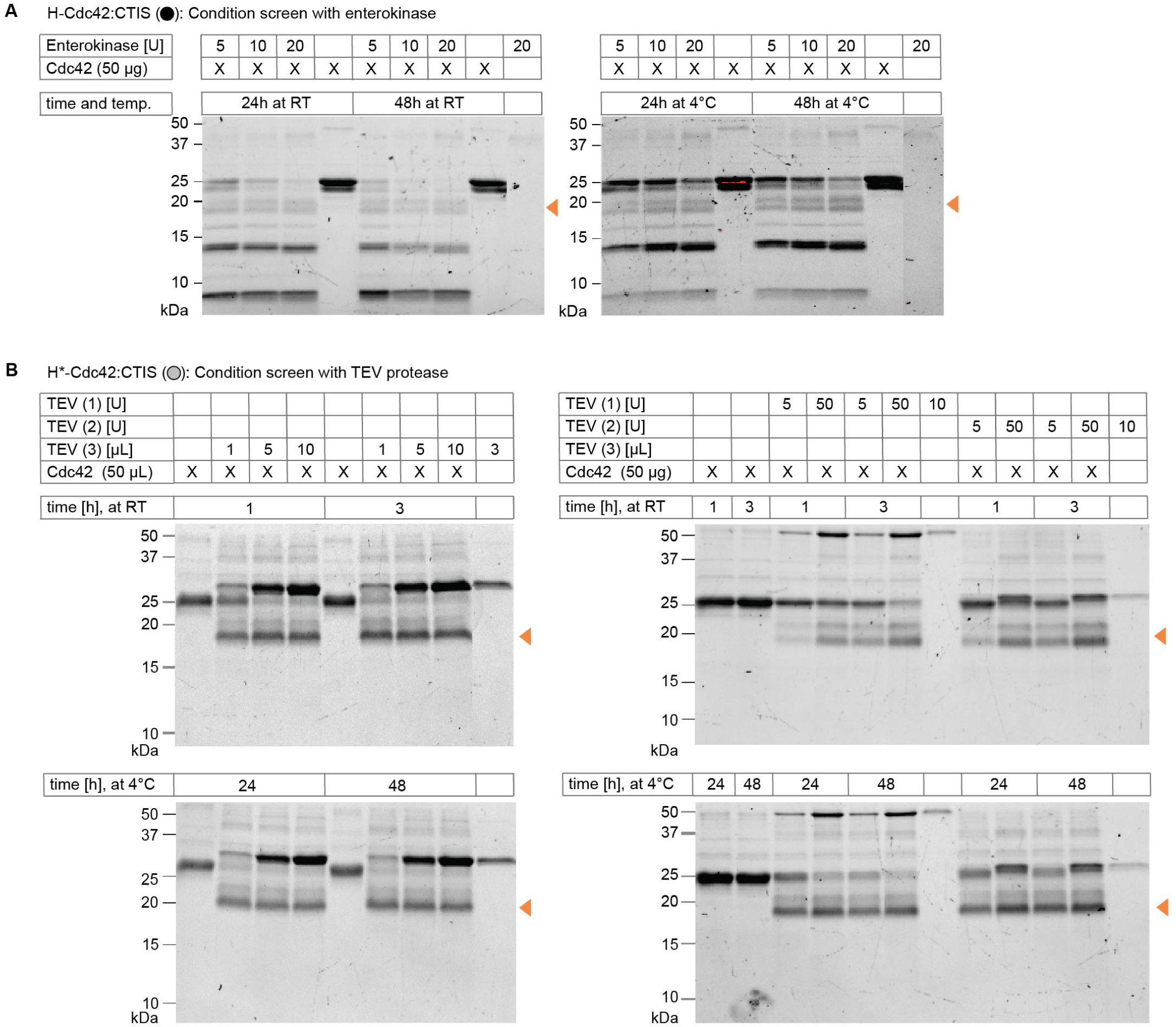
TEV protease, but not enterokinase, is a suitable cleavage enzyme for Cdc42. (A) Cleavage reactions of enterokinase with Cdc42 (H-Cdc42:CTIS) lead mainly to undesired cleavage products. The images shown are SDS-Page. (B) Cleavage reactions of TEV protease with Cdc42 (H*-Cdc42:CTIS) lead to the desired cleavage products. Images shown are SDS-Page of cleavage reaction products of reactions using two commercially available TEV proteases (TEV (1), TEV (2)) and one enzyme that was purified in-house (TEV (3), 125 µM) and H*-Cdc42:CTIS (63 μM).

**S5 Figure 2.**
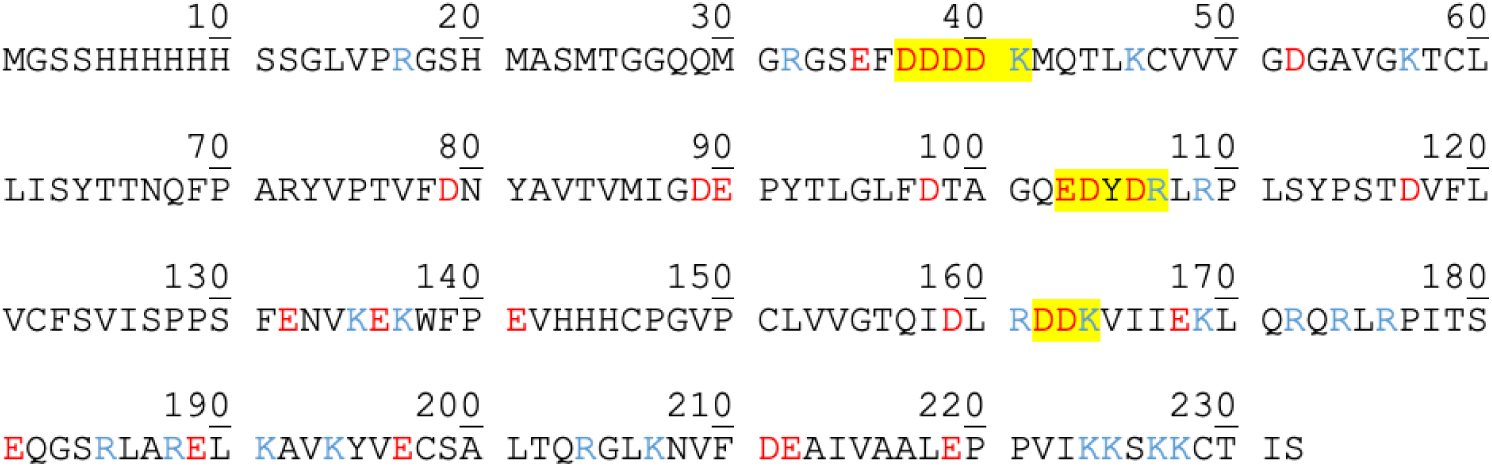
Sequence of H-Cdc42:CTIS (25.7 kDa). Acidic AAs are shown in red and basic AAs are shown in blue. Potential cleavage sites for Enterokinase are highlighted in yellow. Cleavage after ‘…DDDDK’ (site 1) results in the desired products of 4.4 and 21.3 kDa, cleavage after ‘…EDYDR’ (site 2) results in fragments of 11.7 and 14.0 kDa, and cleavage after ‘…DDK’ (site 3) results in 18.2 and 7.5 kDa pieces. Cleavage at site 2 in combination with cleavage at site 3 results in fragments of sizes 18.2, 14.0, 11.7, 7.5, and 6.5 kDa, which partially matches observed fragments on SDS-Page (S5 Fig. 1a).

## Supplement S6

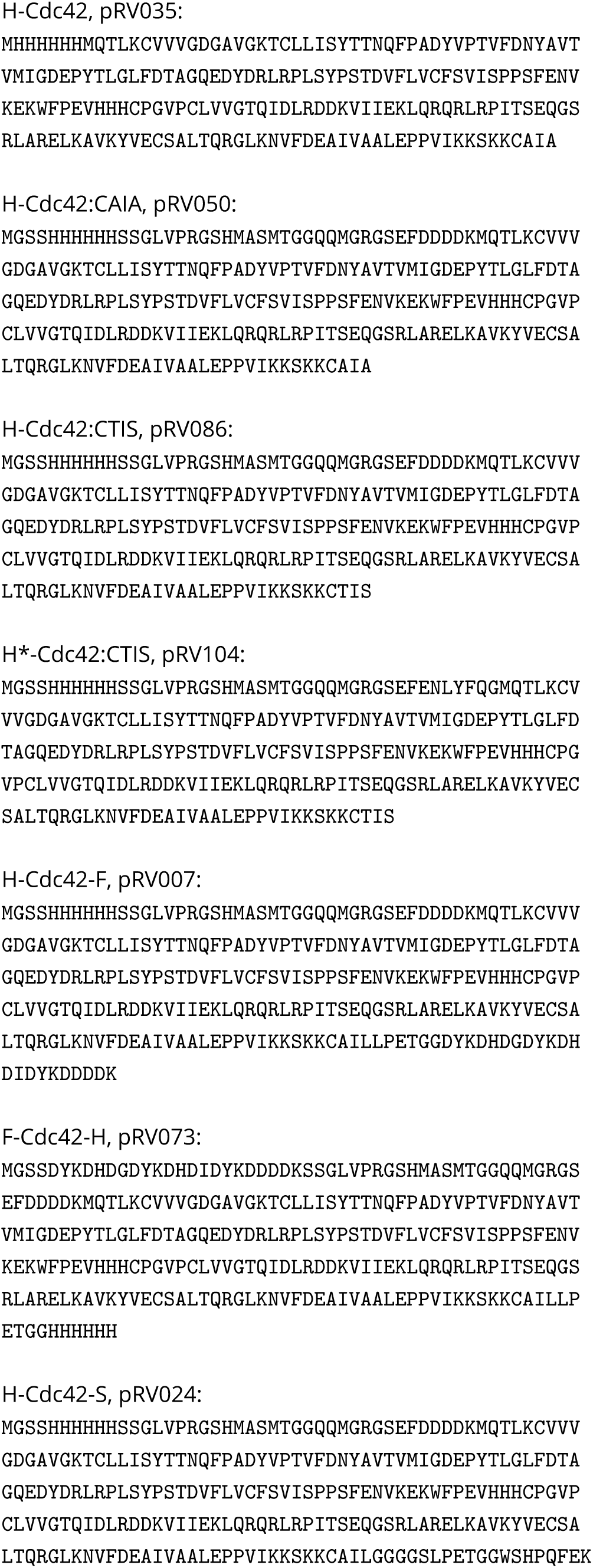
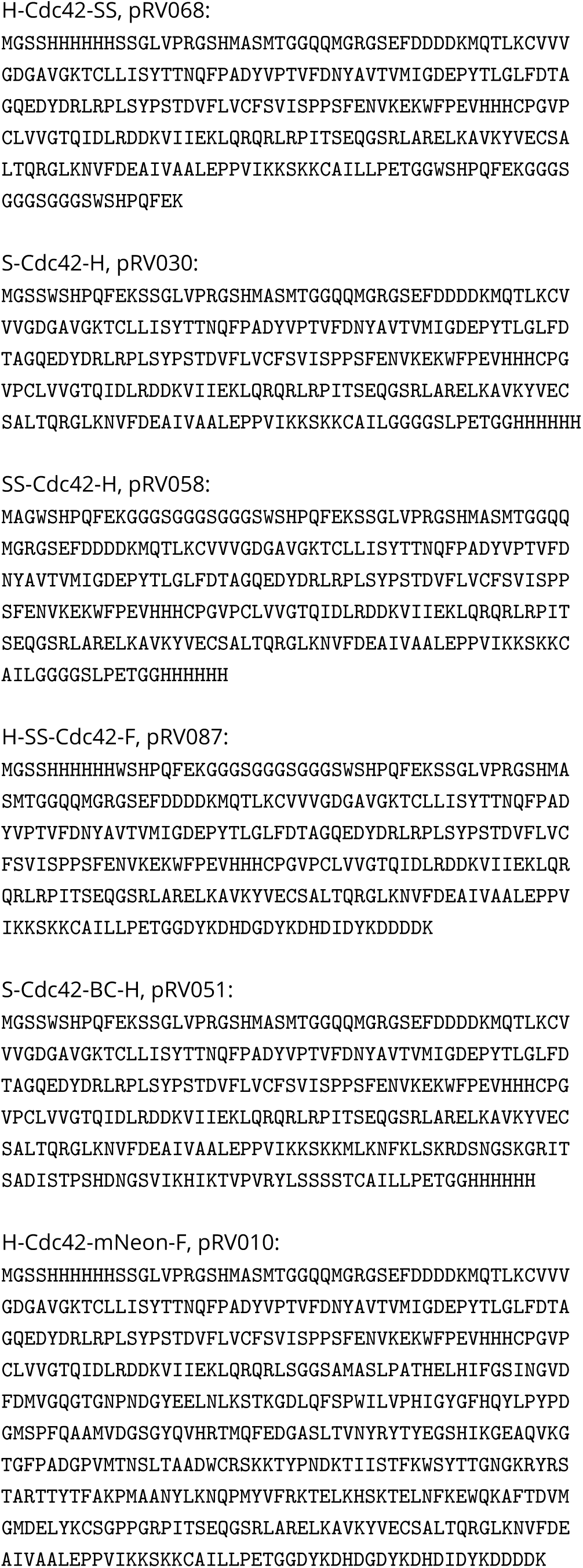
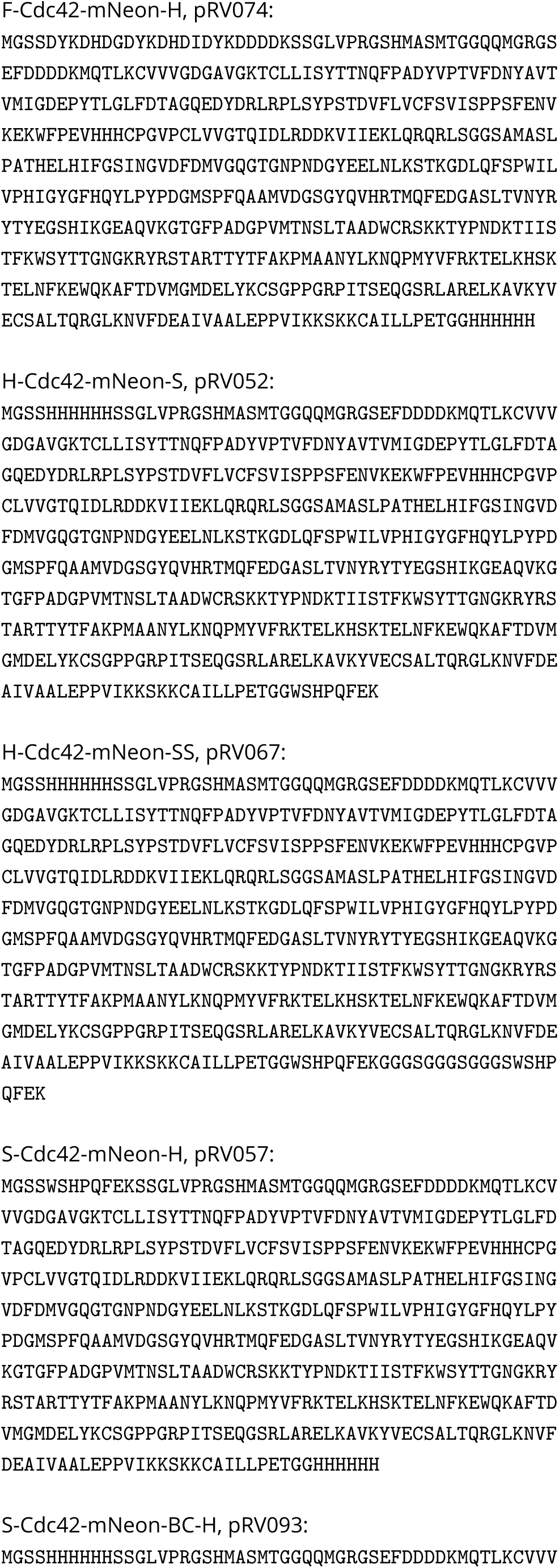
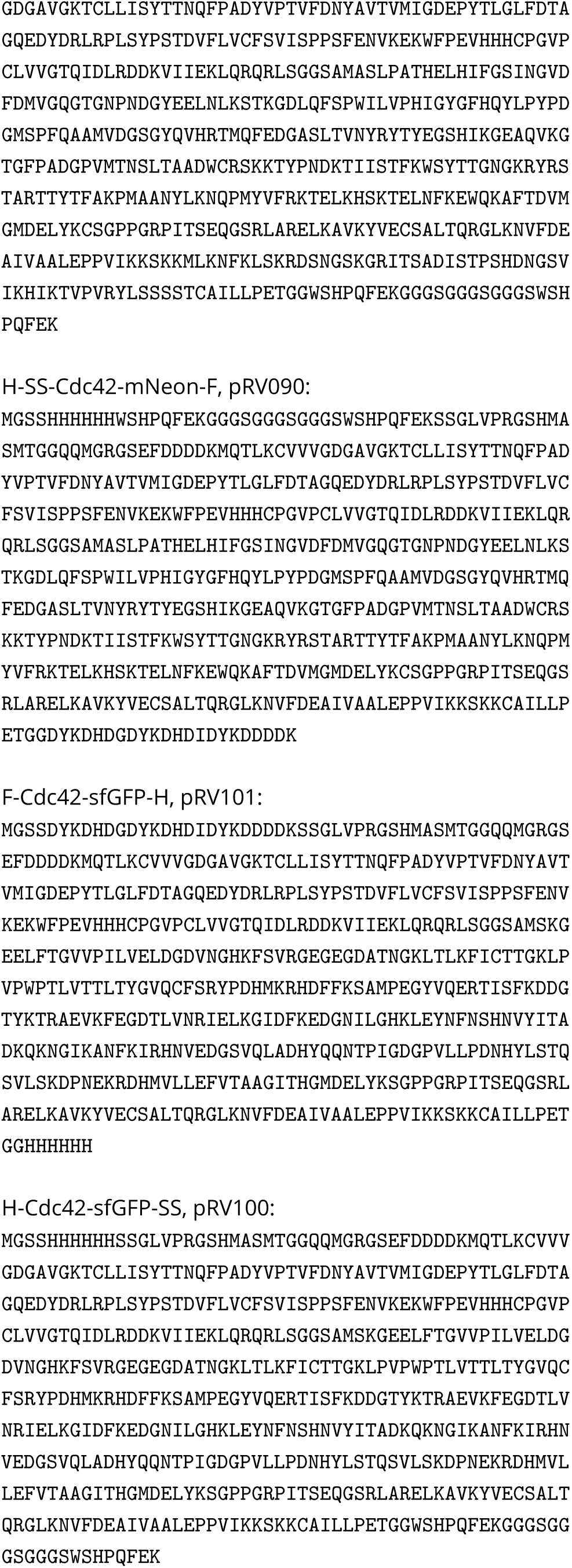
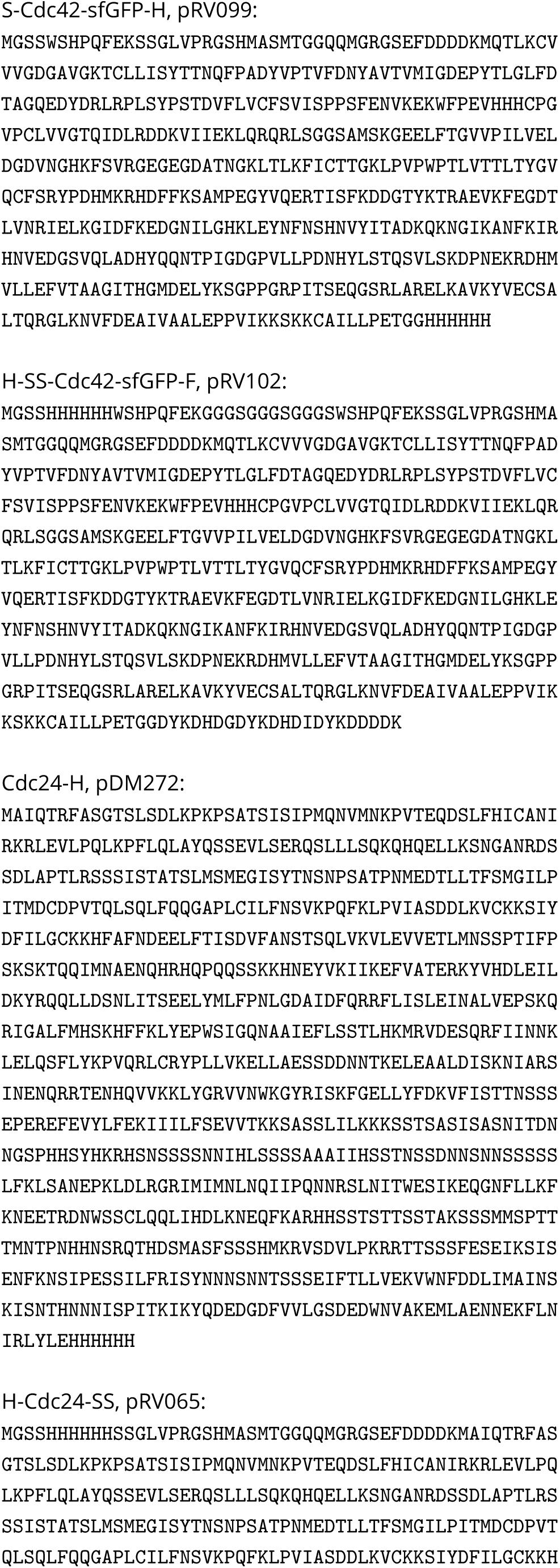
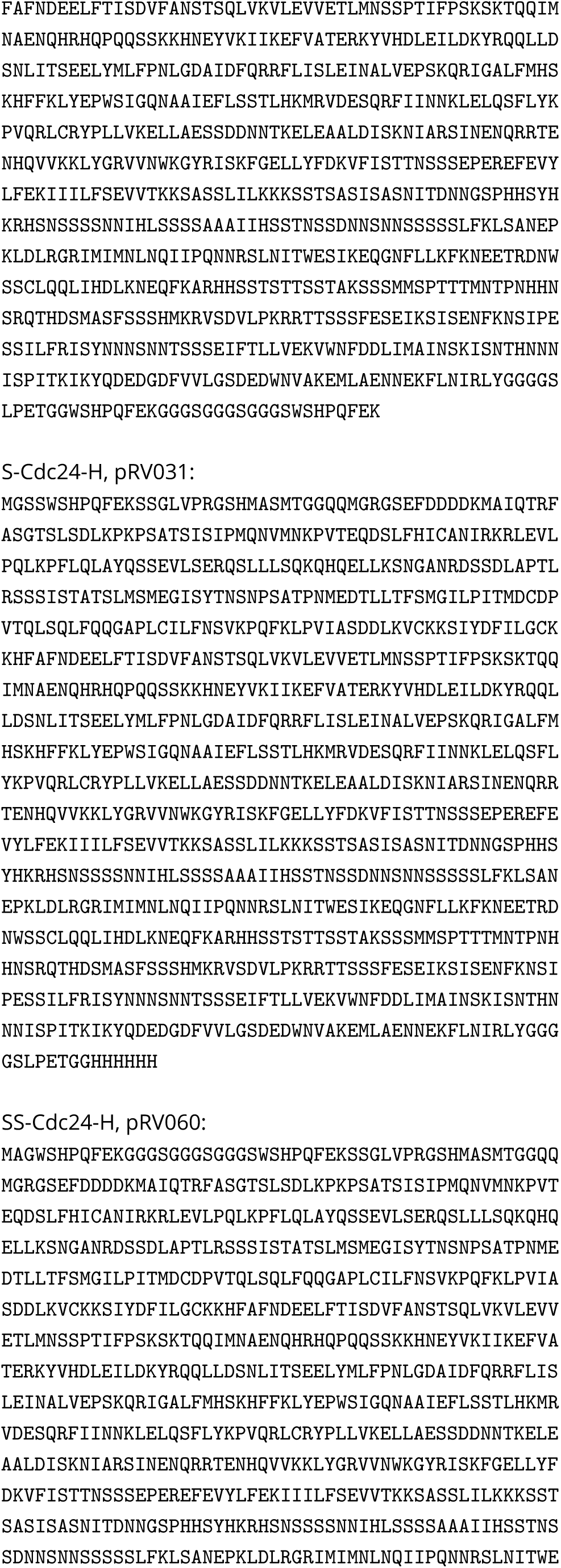
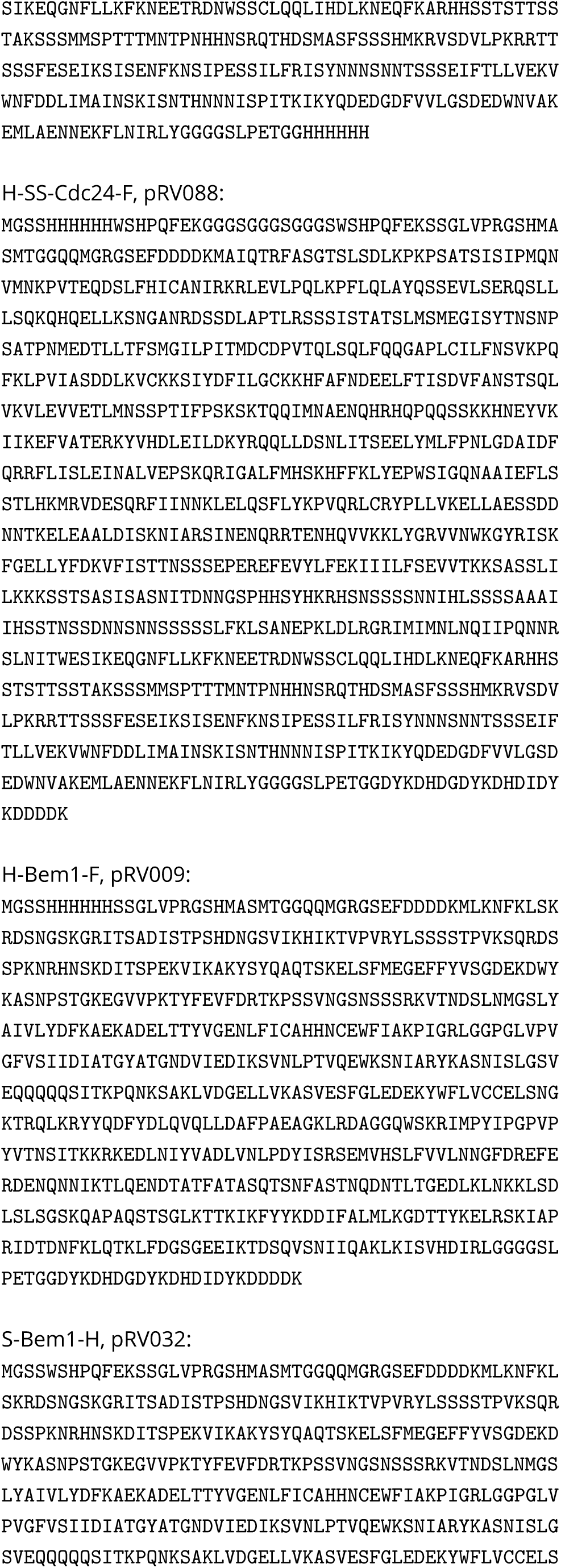
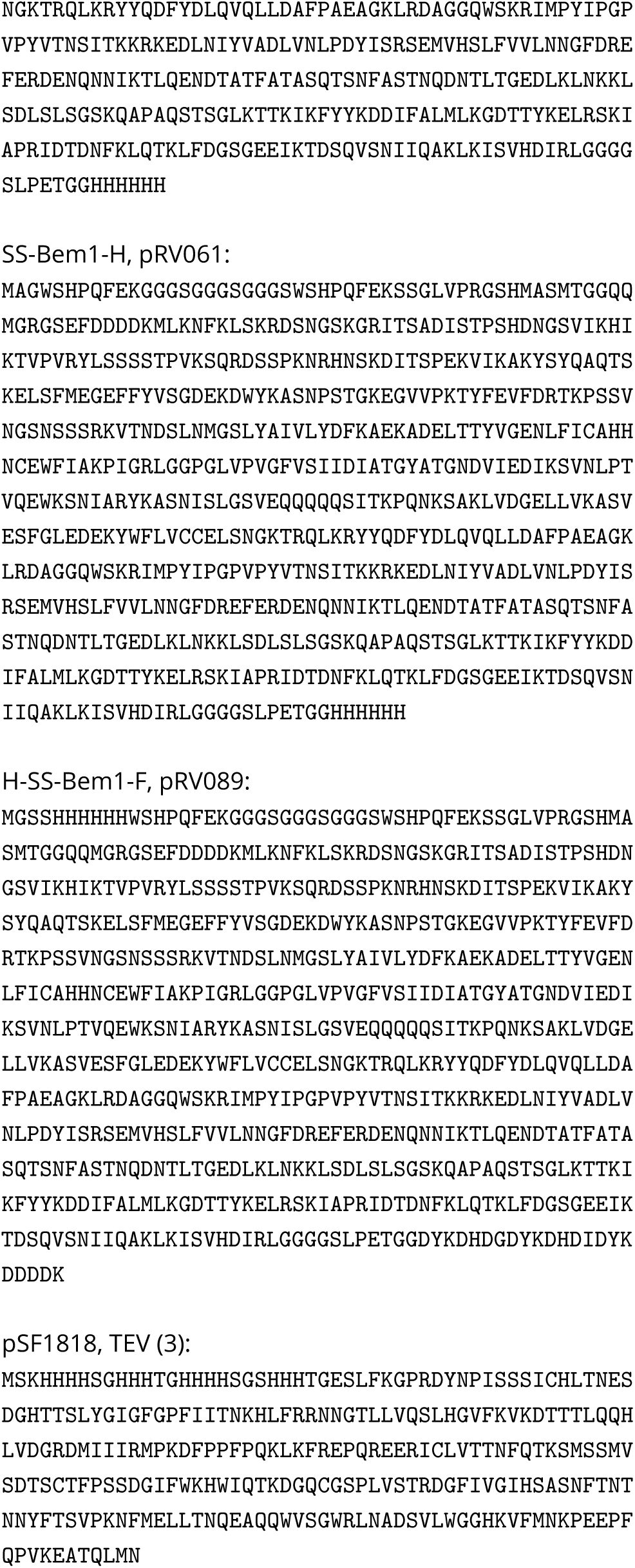

## Supplement S7

**S7 Table 1.**
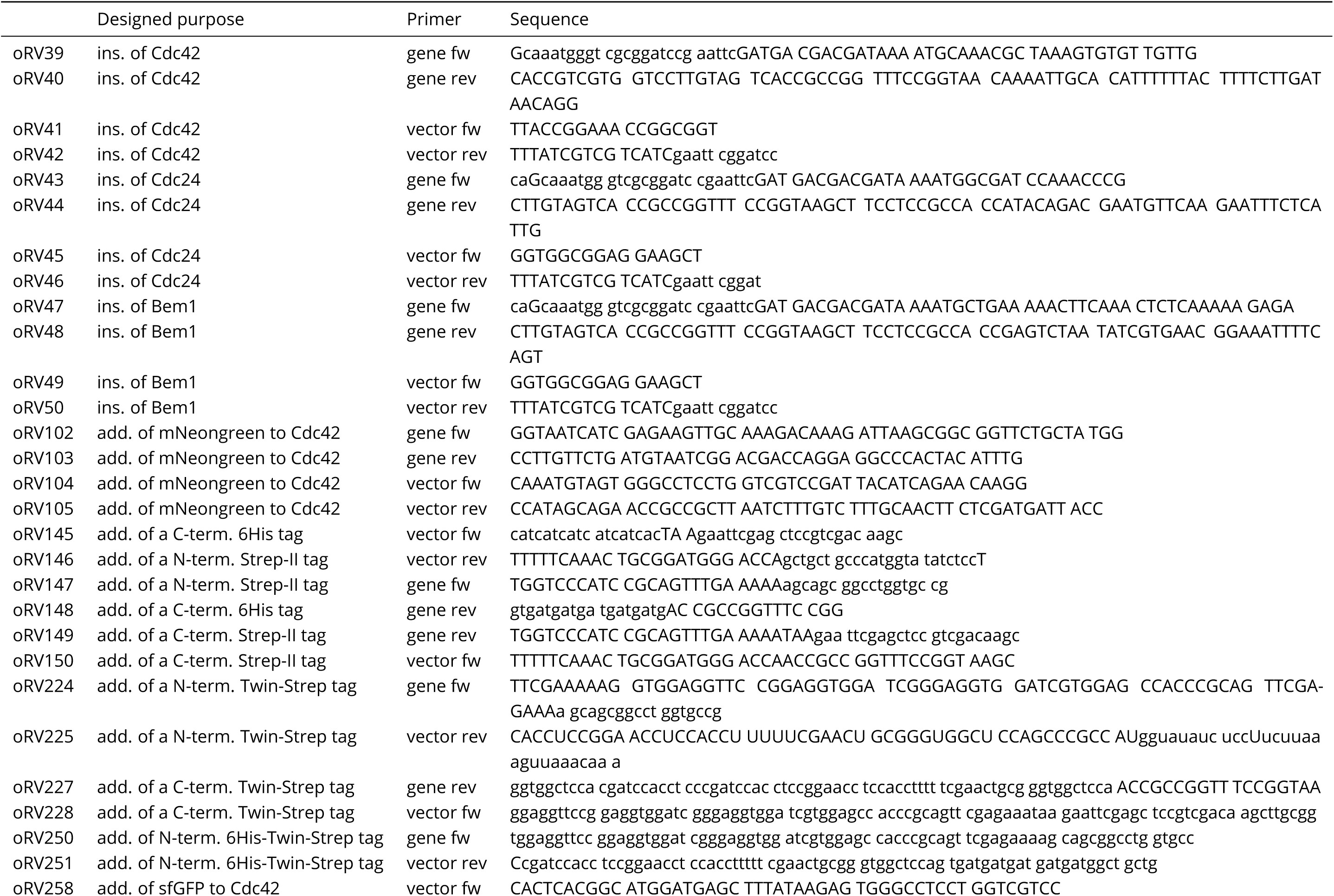
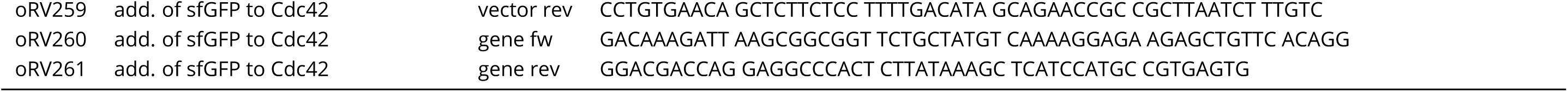
Primer overview. List of primers used to generate plasmids of Tab 2 and 3. Abbreviations: ins: insertion; add: addition; fw: forward; rev: reverse.

## Footnotes

1 Newer columns (bought 2020 or later) required a higher amount of imidazole in the lysis and washing buffer, as indicated by the recommendation ‘use 20-40 mM imidazole in sample and binding buffer for highest purity’ on the column package. For these columns the amount of imidazole in lysis and His-AC washing buffer was increased to 50 mM.

